# Differential conformational dynamics in two type-A RNA-binding domains drive the double-stranded RNA recognition and binding

**DOI:** 10.1101/2023.08.06.552166

**Authors:** Firdousi Parvez, Devika Sangpal, Harshad Paithankar, Zainab Amin, Jeetender Chugh

## Abstract

TAR RNA binding protein (TRBP) has emerged as a key player in the RNA interference (RNAi) pathway, wherein it binds to different pre-miRNAs and siRNAs, each varying in sequence and/or structure. We hypothesize that TRBP displays dynamic adaptability to accommodate heterogeneity in target RNA structures. Thus, it is crucial to ascertain the role of intrinsic and RNA-induced protein dynamics in RNA recognition and binding. We have previously elucidated the role of intrinsic and RNA-induced conformational exchange in the double-stranded RNA-binding domain 1 (dsRBD1) of TRBP in shape-dependent RNA recognition. The current study delves into the intrinsic and RNA-induced conformational dynamics of the TRBP-dsRBD2 and then compares it with the dsRBD1 study carried out previously. Remarkably, the two domains exhibit differential binding affinity to a 12 bp dsRNA owing to the presence of critical residues and structural plasticity. Further, we report that dsRBD2 depicts constrained conformational plasticity when compared to dsRBD1. Although, in the presence of RNA, dsRBD2 undergoes induced conformational exchange within the designated RNA-binding regions and other residues, the amplitude of the motions remains modest when compared to those observed in dsRBD1. We propose a dynamics-driven model of the two tandem domains of TRBP, substantiating their contributions to the versatility of dsRNA recognition and binding.

**Significance Statement:** Exploring the intricacies of RNA-protein interactions by delving into dynamics-based measurements not only adds valuable insights into the mechanics of RNA-protein interactions but also underscores the significance of conformational dynamics in dictating the functional outcome in such tightly regulated biological processes. In this study, we measure intrinsic and RNA-induced conformational dynamics in the second dsRBD, i.e., TRBP-dsRBD2, and compare the same with that carried out in the first dsRBD (TRBP-dsRBD1) of TRBP protein, a key player of the RNAi pathway. The study unveils the differential conformational space accessible to the two domains of TRBP, even though they both adopt a canonical dsRBD fold, thereby affecting how they interact with target RNAs.

## Introduction

RNA-binding proteins (RBPs) play a crucial role in every aspect of RNA biology, including folding, splicing, processing, transport, and localization (Dreyfuss et al., 2002; Fu et al., 2016; Jiang and Baltimore, 2016; Kim et al., 2009; Wilkinson and Shyu, 2001). RBPs can be broadly classified into single-stranded RNA-binding proteins (ssRBPs) and double-stranded RNA-binding proteins (dsRBPs). While ssRBPs are generally sequence-specific as the interaction points include exposed nucleobases (Auweter et al., 2006; Daubner et al., 2013); dsRBPs tend to target a particular structural fold, especially the A-form RNA duplex via phosphate backbone and sugar hydroxyl protons (Masliah et al., 2012). dsRBPs interact with highly structured dsRNAs via their double-stranded RNA-binding motifs/domains (dsRBMs/dsRBDs).

dsRBDs are 65-68 amino acid long domains (Johnston et al., 1992) that contain a highly conserved secondary structure (α_1_-β_1_-β_2_-β_3_-α_2_), where the two α-helices are packed against an antiparallel β-sheet formed by three β-strands (Bycroft et al., 1995; Kharrat et al., 1995). All the dsRBDs have three common RNA-binding regions: (1) the middle of helix α_1_ (E residue), (2) the N-terminal residues of helix α_2_ (KKxAK motif), and (3) the loop between β_1_ and β_2_ strands (GPxH motif), forming a canonical RNA-binding surface that interacts with the consecutive minor, major, and minor groove of a dsRNA (Masliah et al., 2012; Tian et al., 2004). dsRBDs interact with dsRNAs in a non-sequence-specific manner, targeting the sugar 2’-OH groups (dsRBDs do not target dsDNAs) and phosphate backbone (Bevilacqua and Cech, 1996). They are often arranged in a modular fashion to create highly versatile RNA-binding surfaces and have been classified into type A and B (Fierro-Monti and Mathews, 2000; Johnston et al., 1992; Krovat and Jantsch, 1996; Masliah et al., 2012). Type A dsRBDs are highly homologous (> 59%) to the consensus sequence and are involved in RNA-binding, whereas type B has only conserved C-terminal end and helps in stabilizing the protein-RNA/protein complex (Green et al., 1995; Krovat and Jantsch, 1996; Laraki et al., 2008).

dsRBDs, both within and across proteins, exhibit significant variations in their binding modes and affinities towards specific RNA targets. For example, *X. laevis* RNA-binding protein A (XlrbpA) has 3 dsRBDs: xl1 (type A), xl2 (type A), and xl3 (type B), where the xl2 is only able to bind the dsRNA *in vitro* (Krovat and Jantsch, 1996). In Protein Kinase R (PKR), these two types of dsRBDs also demonstrate a differential dynamic behavior, i.e., dsRBD1 (type A) shows plasticity on µs and ps-ns timescale, while dsRBD2 (type B) predominantly depicts dynamics on the ps-ns timescale dynamics (Fierro-Monti and Mathews, 2000; Nanduri et al., 2000). Furthermore, the average order parameter (S^2^; the degree of motion of the backbone N-H vector in a cone, where S^2^ = 1 indicates limited flexibility and 0 indicates maximum flexibility) for the individual α-helices and β-sheets is observed to be lower for dsRBD1 than for dsRBD2, suggesting the latter as a more rigid domain (Fierro-Monti and Mathews, 2000; Nanduri et al., 2000). TAR RNA binding protein (TRBP) harbours two type A (Krovat and Jantsch, 1996) dsRBDs, i.e., dsRBD1 and dsRBD2, where the latter shows a significantly stronger binding affinity (four times) for pre-miR-155 (Benoit et al., 2013). Additionally, studies have reported that dsRBD2 of TRBP displays a higher binding affinity for siRNA and HIV trans-activation response (TAR) RNA (Daviet et al., 2000; Yamashita et al., 2011). It is intriguing to notice that the two type-A dsRBDs originating from the same protein demonstrate a markedly different binding affinity for target RNAs across various cases, as elucidated above. Recent investigations have suggested that the dissimilar behavior of dsRBDs in Dicer (Wostenberg et al., 2012), PKR (Nanduri et al., 2000), and double-stranded-RNA-binding protein 4 (DRB4) (Chiliveri et al., 2017) might be attributed to protein dynamics. Although variations in the binding affinity for the two type-A dsRBDs of TRBP have been documented for several target RNAs, a comprehensive exploration of the differences in the conformational dynamics between these two domains remains to be undertaken.

In this study, we have investigated the role of differential conformational dynamics of the two type A dsRBD domains of TRBP2 (isoform 1 of TRBP) in RNA recognition and binding. First discovered as a TAR RNA-binding protein (39 kDa) involved in HIV-I replication (Gatignol et al., 1991), TRBP was later found to be indispensable to the RNAi pathway (Kim et al., 2014). It is involved in Dicer-mediated pre-miRNA/pre-siRNA cleavage and recruitment of Argonaute protein (Chendrimada et al., 2005). Being a part of the RNA-induced silencing (RISC) complex, TRBP helps in the guide strand selection (Noland et al., 2011; Noland and Doudna, 2013). The guide strand of the mature miRNA directly interacts with the target mRNA to regulate its expression, while the passenger strand is cleaved off by the RNA-induced silencing complex (RISC) complex (Leuschner et al., 2006; Matranga et al., 2005). TRBP and its homologs across different species such as Loquacious (Loqs in *D. melanogaster*), R2D2 (*D. melanogaster*), DRB1-3,5 (*A. thaliana*), RNAi defective 4 (RDE-4 in *C. elegans*), Xlrbpa (*X. laevis*) (Eckmann and Jantsch, 1997), and the PKR activator (PACT) protein (mammals) (Peters et al., 2001) have conserved arrangement of three consecutive dsRBDs; of which, the two N-terminal domains (type A) are known to bind dsRNAs, whereas the third domain is known for protein-protein interactions. We have recently established the role of TRBP2-dsRBD1 dynamics in dsRBD-dsRNA interactions and proposed that dsRBD1 adopts a conformationally dynamic structure to recognize a set of topologically different dsRNA structures (Paithankar et al., 2022). The µs timescale motions were found to be present all along the dsRBD1 backbone with higher frequency motions (*k*_ex_ > 50 kHz) in the RNA-binding sites. The data suggested that the presence of conformational exchange in the µs timescale could help the dsRBD1 to dynamically tune itself for targeting conformationally distinct dsRNA substrates.

In this work, we have compared TRBP2-dsRBD1 with TRBP2-dsRBD2 in terms of the structure, dynamics (intrinsic and RNA-induced), and dsRBD-dsRNA interactions in the two type-A domains. We have measured motions in the TRBP2-dsRBD2 at ps-ns and µs-ms timescale dynamics by NMR relaxation dispersion experiments in apo-state and studied its perturbation in the presence of an A-form duplex RNA. The 12 bp duplex RNA used for the study is called the D12 RNA derived from the miR-16-1 (Figure 1F and Table S1). We also compared the apo- and RNA-bound conformational dynamics measured in dsRBD2 with that of dsRBD1. Based on our observations, we propose that the differential protein dynamics and its perturbation in the presence of RNA in the two dsRBDs enables them to recognize a variety of RNA substrates and may lead to diffusion along the length of the RNA.

**Figure 1:**
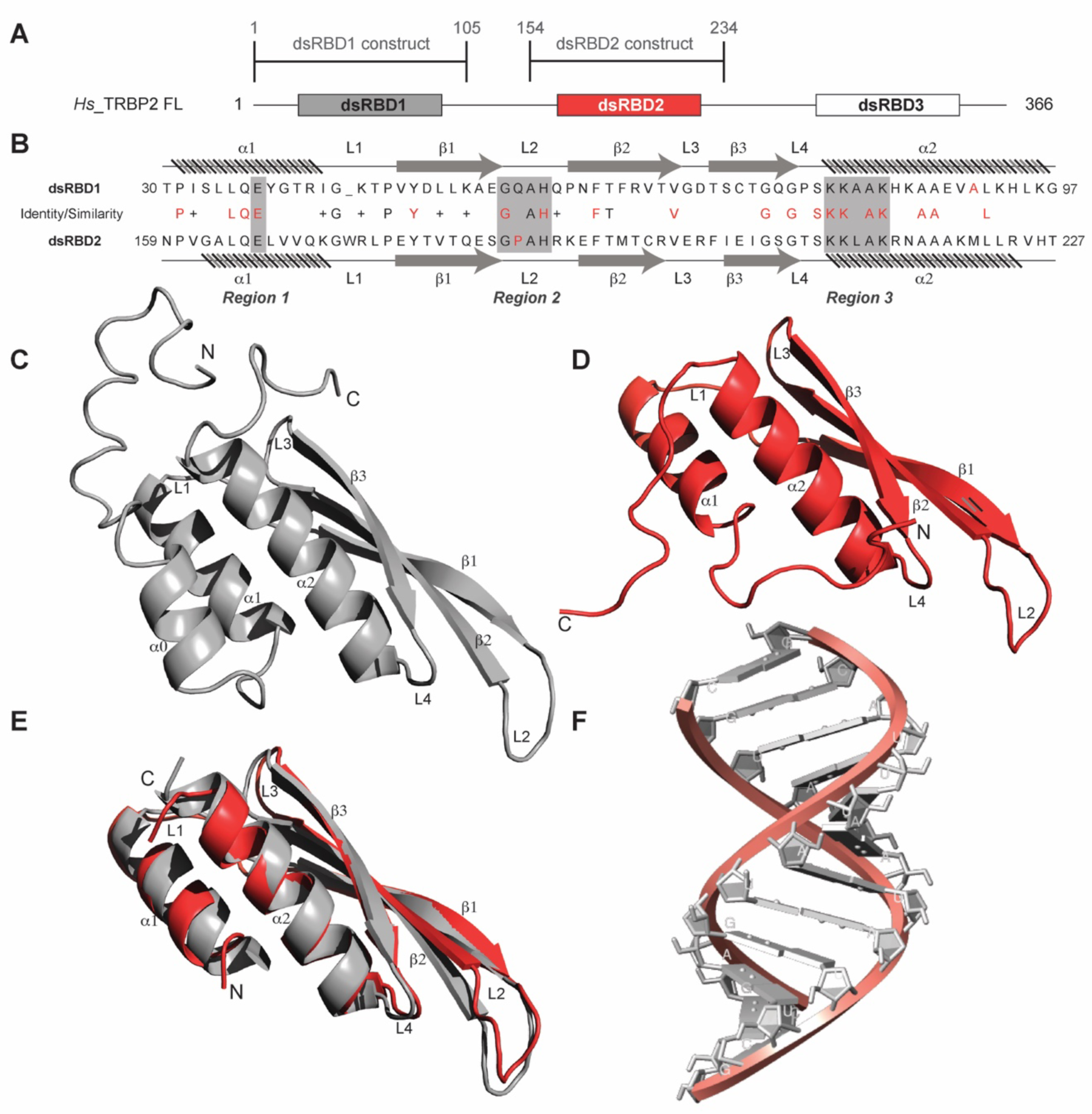
(A) dsRBD constructs [TRBP2-dsRBD1 (1–105 aa) and TRBP2-dsRBD2 (154–234 aa] of Human TRBP2 full-length protein used in this study. (B) Sequence alignment of the two constructs mentioned in (A). CS-Rosetta structures of (C) TRBP2-dsRBD1 (Paithankar et al., 2018), (D) TRBP2-dsRBD2, (E) an alignment of core residues of dsRBD1 and dsRBD2, (F) Model structure of 12 bp duplex D12 RNA.

## Results and Discussion

### Structural and dynamical comparison of apo TRBP2-dsRBD1 and TRBP2-dsRBD2

Primary sequence comparison of TRBP2-dsRBD1 and TRBP2-dsRBD2 domain constructs (Figure 1A) revealed 30% identity and 38% similarity. The consensus for dsRNA-binding has been marked in red in Figure 1B. The two reported RNA-binding regions 1 (E) and 3 (KKxAK) of both the domains were matched to the consensus (Figure 1B). While RNA-binding region 2 (GPxH) (Figure 1B, Region 2) of TRBP2-dsRBD2 was an exact match to the consensus, dsRBD1 harbored a mutation in region 2 (P56Q). Proline is a rigid amino acid with one less dihedral angle; it imparts flexibility to the backbone by causing secondary structure breaks (Imai and Mitaku, 2005; Krieger ^e^t al., 200^5^). Thus, the presence of conserved Pro186 in the β_1_-β_2_ loop of dsRBD2 (Figure 1B, Region 2) may perturb the β_1_-β_2_ loop region plasticity, thereby making it more accessible to the incoming RNA partner. Additionally, dsRBD2 contains a KR-helix motif in the α_2_-helix (Figure 1B, Region 3) known to increase its binding affinity, as reported earlier by Daviet *et al*. (Daviet et al., 2000). Owing to these tightly conserved RNA-binding regions and the presence of additional KR-helix motif, dsRBD2 could make stronger contact with RNA.

The SEC-MALS study of TRBP2-dsRBD2 in the experimental conditions used showed a single monomeric species in solution (Figure S2). The ^1^H-^15^N HSQC spectrum (Figure S3) of TRBP2-dsRBD2 indicated a well-folded protein. It (154–234 aa) was compared to the previously assigned spectrum of the TRBP2-D1D2 construct (19–228 aa) (Benoit and Plevin, 2012) to transfer the backbone amide resonance assignments. Resonance assignments were further confirmed using a set of double and triple resonance experiments, and a total of 73 non-proline residues were assigned as shown in the representative ^1^H-^15^N HSQC (Figure S3). Overall, 89% ^1^H (151/169), 74% ^13^C (124/168), and 87% ^15^N (73/84) resonances from the backbone, and 43% ^1^H (174/402) and 25% ^13^C (59/238) from the side chains were assigned. Similar to the CS-Rosetta structure calculated previously for dsRBD1 (Figure 1C) by our group (Paithankar et al., 2018), the CS-Rosetta structure calculated for the dsRBD2 (Figure 1D) matched well with the previously reported structures in terms of characteristic dsRBD fold. The core residues (159-227 aa) of dsRBD2 were found to adapt the characteristic dsRBD αβββα fold with an RMSD of 1.284 Å when compared to the previously reported solution structure of dsRBD2 (2CPN; (Yamashita et al., 2011)). An alignment of core structures between dsRDB1 and dsRBD2 (Figure 1E) yielded an RMSD of 0.894 Å, indicating a close match between the two domains. Despite their core length being identical (69 aa), the individual secondary structure spans were found to be different, as listed in Supplementary Table 3. The flexible regions, like loops 1 and 2 depicted in Figures 1B and 1E, are longer in dsRBD2, whereas the structured regions β_2_, β_3_, and α_2_ are longer in dsRBD1. Most importantly, loop 2 (β_1_-β_2_ loop) — critical for RNA-binding (Masliah et al., 2012)— is equal to the canonical length in dsRBD2, while in dsRBD1, it is shorter by 1 residue (Figure 1E).

Comparing the MD simulation data of the core residue-specific RMSF plots of these two type A dsRBD domains (Figure 2A), we observe that the average RMSF of the core domain of dsRBD2 is 0.22 nm, which is almost half as compared to dsRBD1 RMSF average of 0.43. This, in turn, indicates that dsRBD2 is overall more rigid among the two. The most interesting fact is that loop 2 (shown in Figure 1B) of dsRBD2 is very flexible compared to the rest of the core, with an average RMSF of 0.49 nm (more than double the core average), as depicted in Figure 2A. Hence, we propose that this long, conserved, and dynamic loop 2 might help dsRBD2 to establish strong contacts in the minor grooves of dsRNA partners. An effect of conserved insertions in the β_1_-β_2_ loop on RNA binding has been previously discussed in the crystal structure of full-length *Arabidopsis* HEN1 in complex with a small RNA duplex (Huang et al., 2009). The MD simulation data of RMSD against the simulation time of TRBP2-dsRBD2 showed that the protein remained stable during the course of triplicate simulations, as seen previously in the case of dsRBD1 (Figure 2B).

**Figure 2:**
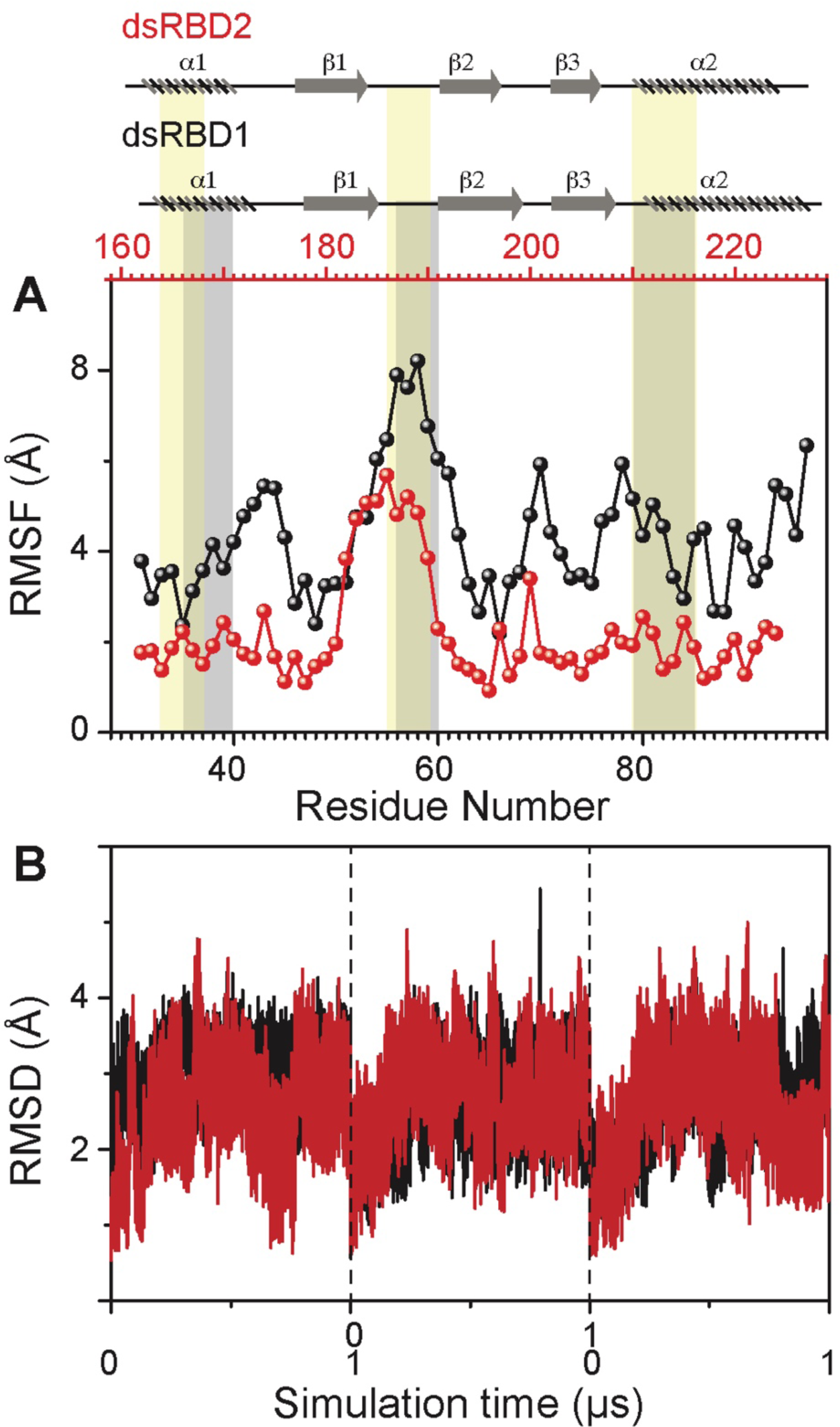
(A) RMSF profile (by C_α_) of core dsRBD1 (black) and dsRBD2 (red). The secondary structure for the two domains has been shown on the top, and three RNA-binding regions in dsRBD1 and dsRBD2 have been highlighted using vertical grey and yellow bars, respectively. (B) RMSD of profiles of dsRBD1 (black) and dsRBD2 (red) over 1 μs simulation time. RMSD values for data measured in triplicate have been separated by vertical dashed lines.

To gain insights into the picosecond to nanosecond (ps-ns) timescale motions of apo-dsRBD2, NMR spin relaxation data (*R*_1_, *R*_2_, and [^1^H]-^15^N NOE) was recorded on 600 and 800 MHz NMR spectrometers. The set of plots for *R*_1_, *R*_2_, and [^1^H]-^15^N NOE at both 600 and 800 MHz magnetic fields showed the expected trend (Figure S4). *R*_1_ in the structured residues showed lower values at a higher magnetic field and similar values in the terminal regions at the two magnetic fields (Figure S4A). The increase of core *R*_1_ rates (decreasing *T*_1_) with increasing magnetic field indicates that most of the core residues lie on the right-hand side of the *T*_1_ minima in the standard plot of *T*_1_ vs. ω*τ*_c_, where ω is the frequency of motions, *τ*_c_ and is rotational correlation time (Bloembergen et al., 1948). The core residues had average *R*_1_ rates of 1.43 ± 0.05 s^-1^ (Figure 3). The *R*_2_ rates (Figure S4B) of a few N-terminal (N159) and core residues (A163, V169, R174, E177, V180, R189, M194, R197, V198, G206, G207, G208, K210, L212, 221, 224 and A227) marginally increased (0.1–3 s^-1^) when measured at 800 MHz than when measured at 600 MHz, suggesting a very insignificant presence of the *R*_ex_ component in the linewidth of these residues in the experimental conditions. The dsRBD2 core N- and C-terminal ends (S151-E157, N159, T227, V228, L230, A232) and the loop residues (E177, G185, E191, S209, K210) exhibited relatively higher flexibility as indicated by lower *R*_2_ rates in Figure 3B and nOe values in Figure 3C. A few residues lying in different secondary structures have been shown in Figure 3B, for e.g.: Q165, L167, and V168 in α_1_; T193 and T195 in β_2_; L212, A213, N216, and L223 in α_2_ exhibited only slightly higher (> 1 s^-1^) than the average *R*_2_ rate (10.92 ± 0.37 s^-1^), indicating a thin conformational exchange in the secondary structured regions.

**Figure 3:**
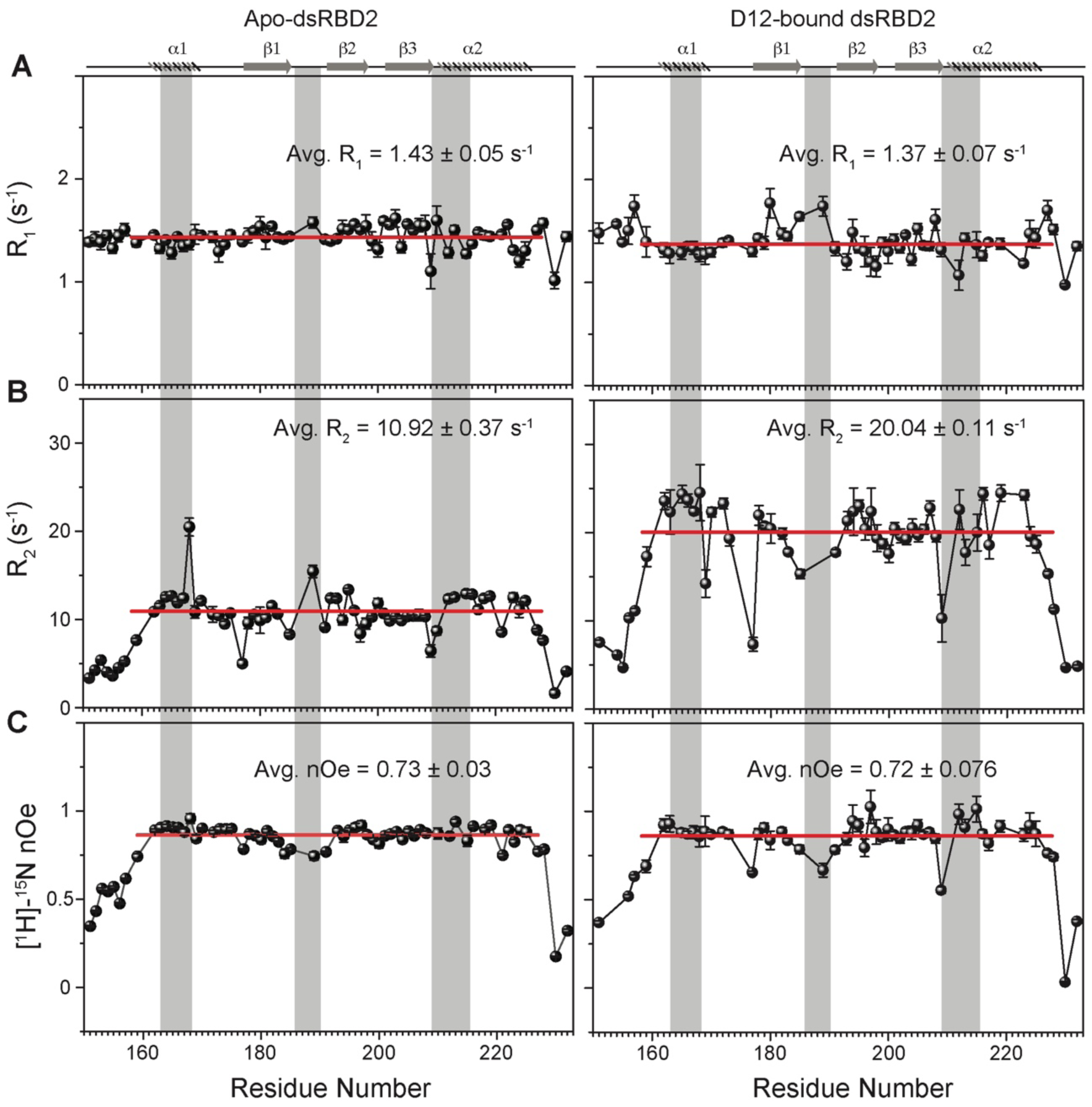
Spin relaxation parameters (A) R_1_, (B) R_2_, and (C) [^1^H]-^15^N nOe plotted against the residues for (left panel) apo TRBP2-dsRBD2 and (right panel) D12-bound TRBP2-dsRBD2. Experiments were recorded on a 600 MHz NMR spectrometer at 298 K. The secondary structure of TRBP2-dsRBD2 has been mentioned at the top, and the RNA-binding region of the protein has been marked in grey vertical columns. Average R_1_, R_2_, and [^1^H]-^15^N-nOe of the core residues (159-227 aa) at 600 MHz is depicted in the green bar, and at 800 MHz is depicted in the red bars.

The core average *R*_1_ (dsRBD1 = 1.52 ± 0.03 s^-1^; dsRBD2 = 1.43 ± 0.05 s^-1^) and *R*_2_ rates (dsRBD1 = 9.83 ± 0.17 s^-1^; dsRBD2 = 10.92 ± 0.37 s^-1^) for the two domains were very similar (Figure S15 of (Paithankar et al., 2022)). In contrast, dsRBD1 (Figure S15C of (Paithankar et al., 2022)) had significantly lower average heteronuclear steady-state nOe values (0.63 ± 0.02) than dsRBD2 (0.73 ± 0.03) shown in Figure 3C, indicating a more rigid core at the picosecond timescale motions in dsRBD2, thereby supporting the earlier observations made in the MD simulation study (Figure 2).

The extended model-free analysis of the relaxation data for the two core domains allowed us to compare the order parameters S^2^, *R*_ex_ components, and global tumbling time (*τ*_c_). The overall order parameters for dsRBD1 followed the expected trend of higher values in the secondary structured regions and lower values in the terminals and loops (Figure S5 top panel). Interestingly, the secondary structural motifs α_1_, loop 2, and α_2_ (refer to Figure 1B & D), containing RNA-binding regions of dsRBD2, showed a higher rigidity than dsRBD1. The presence of conserved Pro in loop 2 (L2) of dsRBD2 and a longer length of L2 in dsRBD2 might be responsible for the additional observed flexibility in dsRBD2, as also observed in the MD simulations (Figure 2 and Table S3). Loop 1 (W173, L175) of dsRBD2 containing the stabilizing tryptophan (absent in dsRBD1) also showed higher rigidity than dsRDB1, corroborated by the low RMSF values for residues in this region from the MD simulation study. The global rotational correlation time (*τ*_c_) of dsRBD2 was 7.3 ns, and that of dsRBD1 was 7.64 ns, suggesting a more compact structure of dsRBD2 than dsRBD1. This result corroborates the CD-based determination of the melting point of the two constructs. While TRBP2-dsRDB2 exhibited a higher T_m_ (55°C), TRBP2-dsRBD1 melted at 45°C (data not shown), thus suggesting a stronger network of interactions in TRBP2-dsRDB2 (Yamashita ^e^t al., 201^1^). Indeed, it has been shown previously that the tryptophan (W173) present in the α_1_-β_1_ of dsRBD2 (W is absent in dsRBD1 at this position) induced local hydrophobic and cation-π interactions with K171, E199, R200, F201, and V225, thereby enhancing the overall thermal stability of dsRBD2 (Yamashita et al., 2011).

Only 7 residues of dsRBD2 could fit into a model-free model containing the exchange term (*R*_ex_) as opposed to 11 residues in dsRBD1, suggesting that there is significantly lower conformational flexibility at the μs-ms timescale in dsRBD2 shown in Figure S5 bottom panel. While dsRBD2 depicted a low frequency (< 2 s^-1^) contribution to the line width of resonances in the RNA-binding regions (E166, L212, and N216, and L223), dsRBD1 had a larger contribution (2–8 s^-1^) to the line width of resonances present all along the backbone (Figure S5 bottom panel). In summation, the plasticity at the ps-ns timescale is present in both the dsRBDs of TRBP, while the amplitude of motions is found to be largely restricted for the dsRBD2.

Similar to dsRBD1, dsRBD2 does not exhibit motions at a slower ms timescale as probed by the CPMG relaxation dispersion experiments shown in Figure S6. No dispersion was observed in the *R*_2eff_ rates with increasing ϖ_CPMG_, suggesting the absence of motions sensitive to this experiment (Paithankar et al., 2022).

The heteronuclear adiabatic relaxation dispersion (HARD) NMR experiment (Mangia et al., 2010; Traaseth et al., 2012) was used to study NMR spin relaxation in a rotating frame. The dispersion in relaxation rates is created by changing the shape of a hyperbolic secant (HSn, where n = stretching factor) adiabatic pulse that is used to create the spin-lock. HARD experiments are sensitive to the conformational exchange processes occurring on the 10 μs–10 ms timescales. The *R_1ρ_* and *R_2ρ_* relaxation rates showed that with the increase in applied spin-lock field strength from HS1 to HS8, the *R_1ρ_* rates increased (Figure 4A & C) and the *R_2ρ_* rates decreased (Figure 4B & D) for both dsRBD1 and dsRBD2. The extent of dispersion was much less in the core residues of dsRBD2, indicating the absence of higher conformational dynamics in the dsRBD2. The *R_1ρ_* and *R_2ρ_* rates were then fit to Bloch-McConnell equations by using the geometric approximation approach (Chao and Byrd, 2016) to extract the rate of exchange (*k*_ex_) and the excited state population (*p*_B_) (Figures 5A). The number of residues with *k*_ex_ > 5,000 Hz and > 10% *p*_B_ (red and green, medium and big spheres) (Figure 5C) differed vastly between the two domains. Only 3 residues with such conformational exchange in dsRBD2 were identified: R200 (loop 3), L212 and R224 (α_2_); whereas in dsRBD1, the number is 15, distributed along the entire backbone of the protein (Figure 6 of (Paithankar et al., 2022)), similar to PKR-dsRBD1, where 75% of the residues from dsRBM1 showed *R*_ex_ > 1 s^-1^ (Nanduri et al., 2000). Interestingly, all three conserved RNA-binding regions in dsRBD1: α_1_ (L35, Q36), β_1_-β_2_ loop (E54), and α_2_ (N61) were brimming with a *k*_ex_ > 50,000 Hz. To sum it up, most of the dsRBD2 core showed slow conformational exchange (*k*_ex_ < 5000 Hz), depicted in blue small spheres, while the dsRBD1 domain was undergoing significantly faster conformational exchange (*k*_ex_ > 5000 Hz). Taken together, the intrinsic *k*_ex_ profile of TRBP2-dsRBD1 and TRBP-dsRBD2 revealed the presence of 10 μs–10 ms timescale conformational dynamics distributed throughout the core dsRBD rather than being localized in the RNA binding regions; however, the amplitude of motions was found significantly smaller in dsRBD2.

**Figure 4:**
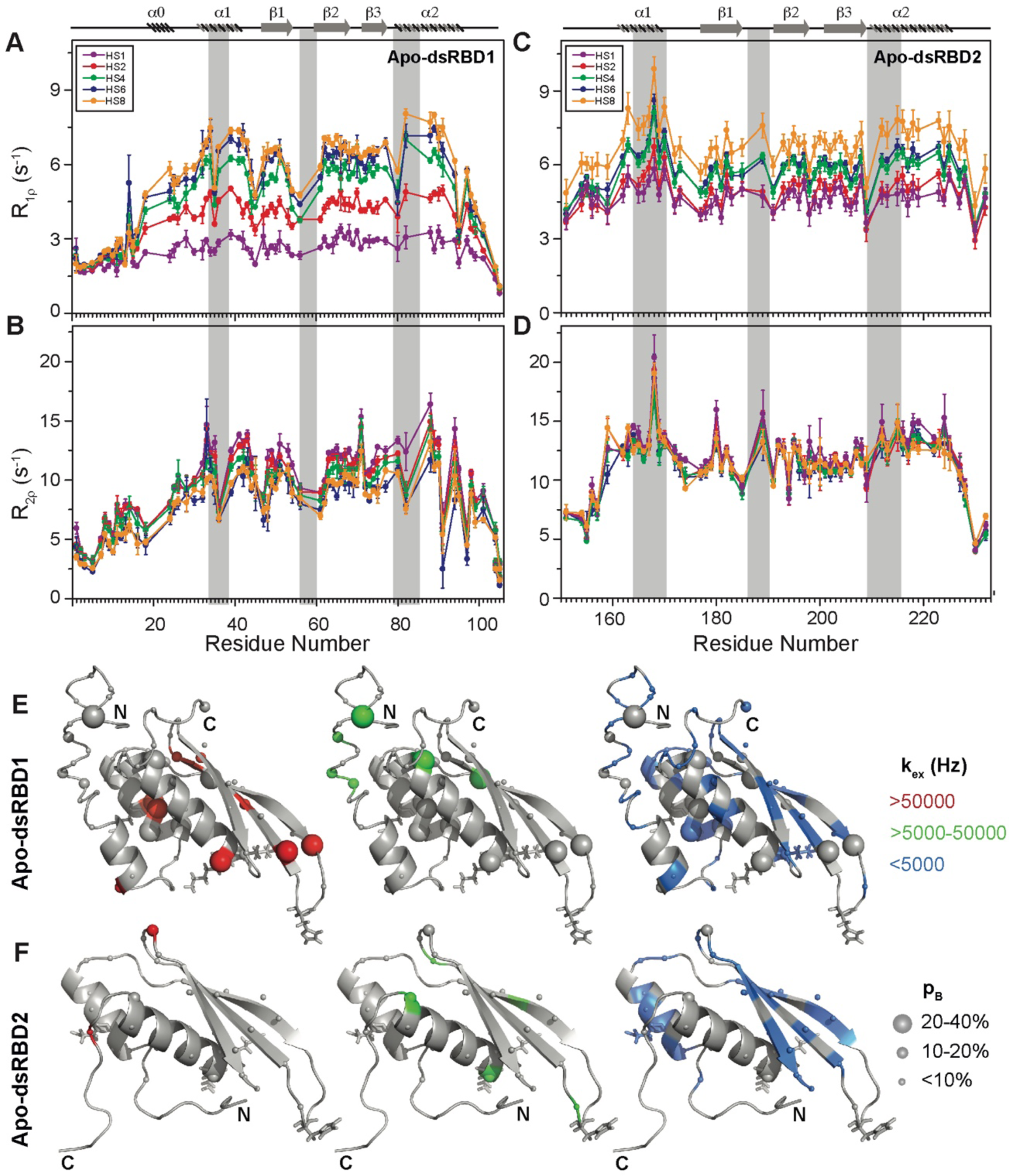
(A) R_1ρ_ data, (B) R_2ρ_ data, recorded on apo ^15^N-TRBP2-dsRBD1, (C) R_1ρ_ data, (B) R_2ρ_ data, recorded on apo ^15^N-TRBP2-dsRBD2 using Heteronuclear Adiabatic Relaxation Dispersion (HARD) experiments on a 600 MHz NMR spectrometer at 298 K, plotted against residue numbers. An increase in the spin-lock field strength is achieved by an increase in the stretching factor of the adiabatic pulse used to create the spin lock, n (in HSn). The secondary structure has been depicted on the top, and three RNA-binding regions have been highlighted using vertical grey bars. Mapping of conformational exchange parameters (rate of exchange between the ground state and excited state (*k*_ex_), and excited state population (*p*_B_)) obtained by fitting the above-described data to a two-state model on the CS-Rosetta structures of (E) apo-dsRBD1, and (F) apo-dsRBD2. Residues have been marked in different colors to highlight the distribution of *k*_ex_ values, and the diameters of the sphere indicate the extent of *p*_B_ along the protein backbone. The RNA-binding residues have been depicted in stick mode.

**Figure 5:**
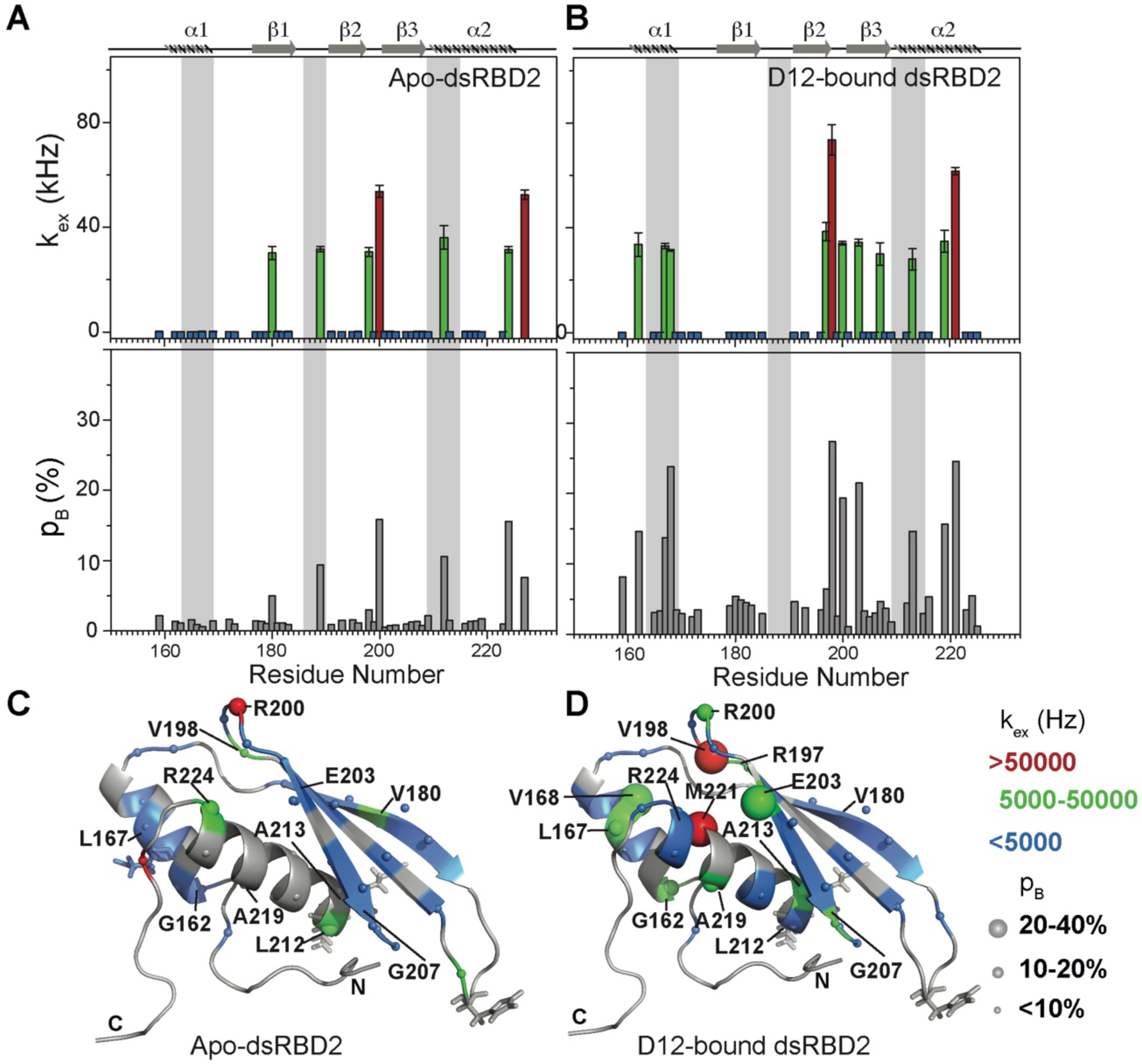
Conformational exchange in (A) apo- and (B) D12-bound-TRBP2-dsRBD2. Top panel: Rate of exchange between the ground state and excited state (*k*_ex_); Bottom panel: excited state population (*p*_B_) as obtained by the geometric approximation method, using the HARD experiment, plotted against residue numbers. Mapping of core *k*_ex_, and *p*_B_, on the CS-Rosetta structure of apo TRBP2-dsRBD2, as extracted for (C) apo TRBP2-dsRBD2 and (D) D12-bound TRBP2-dsRBD2. Different colors highlight the distribution of *k*_ex_ values, and the sphere’s diameter indicates the extent of *p*_B_ along the protein backbone. The RNA-binding residues have been depicted in stick mode.

**Figure 6:**
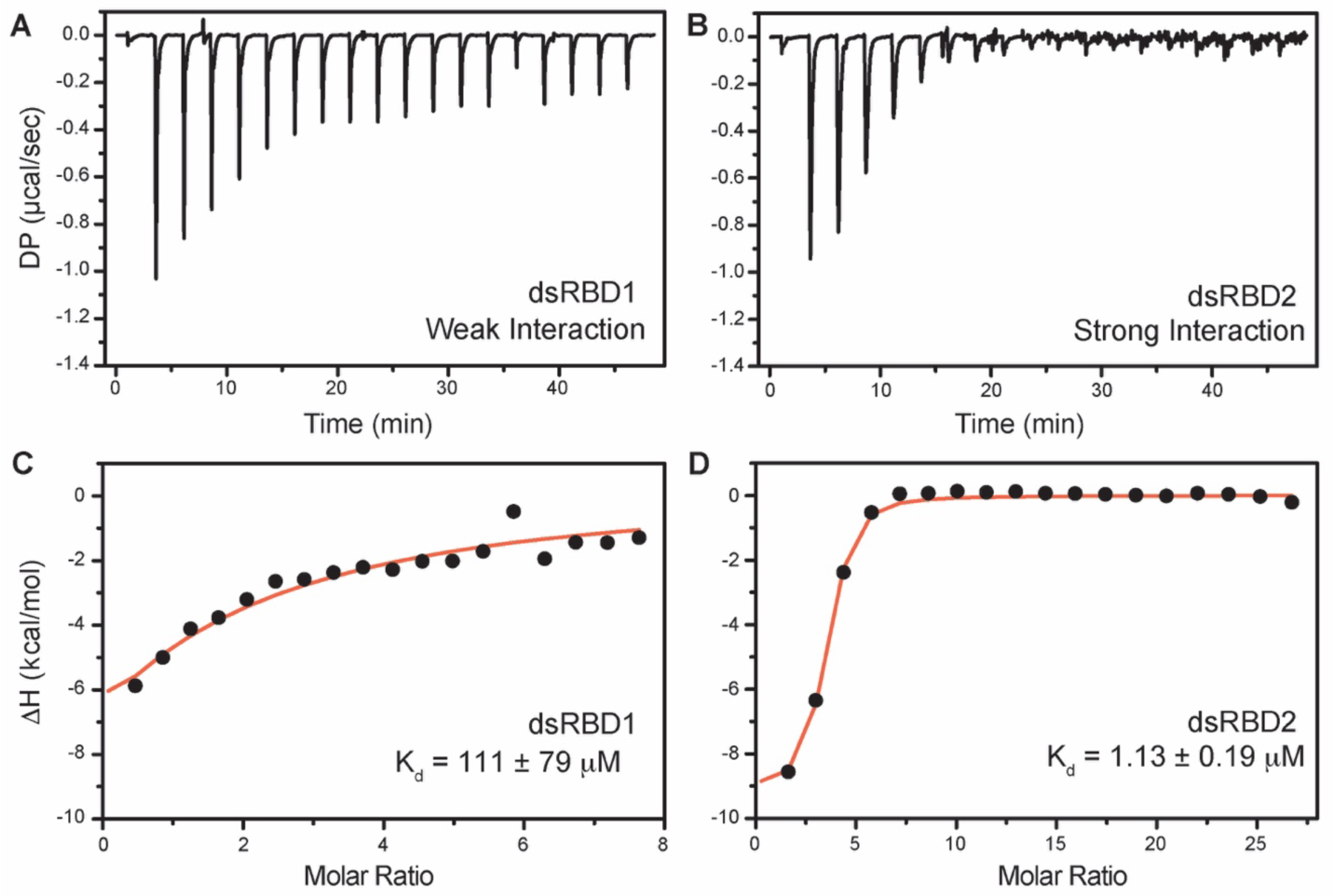
ITC-based binding study of D12 duplex RNA with TBRP2-dsRBD1 and TRBP2dsRBD2. Top panel: the raw differential potential for each injection is plotted against the titration time for (A) TBRP2-dsRBD1 & (B) TBRP2-dsRBD2. Bottom panel: the integrated heat (enthalpy change) upon each injection (black dots) and the data fit for a single set of binding sites (red line) plotted against per mole of injectant for (C) TBRP2-dsRBD1 & (D) TBRP2-dsRBD2.

A similar extensive fast and slower timescale conformational dynamics has also been studied in the case of DRB4 protein (Chiliveri et al., 2017), where the first domain was found to have a large conformational exchange when compared with that of the second domain. The conformational plasticity of dsRBD1 enables it to initiate primary interaction with various structurally and sequentially different dsRNAs in the TRBP2 protein (Paithankar et al., 2022).

### dsRBD2 has a higher affinity for a short double-stranded A-form RNA duplex

Target RNA sequences were designed based on the fact that TRBP interacts with miR-16-1 duplex (miRbase accession no. MI0000070) (Yan et al., 2019). Three mutants of miR-16-1 were created to perturb the RNA shapes as described previously (Paithankar et al., 2022). Briefly, wt miR-16 (22–23 bp) has a bulge (unpaired uridine) and an internal loop (A:A mismatch), miR-16-1-M has only A:A mismatch, (ii) miR-16-1-B has only U-bulge, and (iii) miR-16-1-D has neither the bulge nor internal loop forming a perfect duplex (Figure 1 of (Paithankar et al., 2022)). It has been established that the TRBP2-dsRBD1 is able to recognize a set of topologically different dsRNA structures owing to its high conformational plasticity (Paithankar et al., 2022). In the current study, we have characterized the interaction of TRBP2-dsRBD2 with the same set of topologically different dsRNAs (wt miR-16-1 and mutants) used for TRBP2-dsRBD1 through ^1^H-^15^N HSQC-based NMR titrations shown in Figure S7. The NMR investigation regarding the interaction of TRBP-dsRBD2 with the wt miR-16-1 and the three mutant dsRNAs revealed only chemical shift perturbations (< 0.1 ppm) in the presence of four topologically different RNAs as shown in Figure S7. This indicated that the tertiary structure/fold of the protein remained unperturbed upon RNA binding. The fact that the structure remains unperturbed is consistent with the previous findings reported by Yamashita et al. (Yamashita et al., 2011), where the authors compared the solution structure of TRBP-dsRBD2 structure with the crystal structure of GC10RNA-bound TRBP-dsRBD2 (PDB ID: 3ADL) (Yang et al., 2010) and concluded that the structure of the dsRBD remains unperturbed upon RNA-binding. However, TRBP-dsRBD2, in some cases, has shown the presence of chemical shift perturbation <1.0 ppm at the three RNA-binding regions upon binding with RNA (Benoit et al., 2013; Masliah et al., 2018), indicating subtle conformational changes at the RNA-binding interface (Yamashita et al., 2011; Yang et al., 2010). Furthermore, there were two intriguing observations in the ^1^H-^15^N HSQC-based titrations. First, the amide signals were getting broadened at as low as 0.1 RNA equivalents, suggesting the slow-to-intermediate timescale of binding as has also been observed previously by our and other research groups in TRBP2 dsRBD1, Staufen dsRBD3, hDus2 dsRBD, MLE dsRBD2, DBR4 dsRBD1, PKR dsRBD, and Dicer dsRBD (Ankush Jagtap et al., 2019; Bou-Nader et al., 2019; Chiliveri et al., 2017; Paithankar et al., 2022; Ramos et al., 2000; Ucci et al., 2007; Wostenberg et al., 2012; Yadav et al., 2019). Second, not only the reported RNA-binding residues but the entire backbone was undergoing line broadening, suggesting the presence of RNA-induced motions in the entire backbone. Since the protein is not saturated with the RNAs at 0.1 RNA equivalents (considering a reported K_d_ = 1.7 μM for a 22 bp dsRNA (Acevedo et al., 2015), [protein] = 50 μM, [RNA] = 5 μM, fraction bound of protein < 10%) we can rule out an increase in the size due to the formation of a stable protein-RNA complex causing line-broadening. Upon excess addition of RNA (to 1 equivalents of miR-16-1-M), the backbone amide signals were not recovered (Figure S7 top left panel), indicating the RNA-protein interaction does not seem to come out of the local minima of slow-to-intermediate exchange regime.

The amide NMR signals were, however, partially recovered by shortening the length of the RNA duplex to 12 bp D12 RNA. The line broadening was delayed till 0.35 equivalents of D12 RNA as compared to 0.1 equivalents for the longer RNAs (Figure S8). The excessive line broadening all along the backbone and recovery of the same by reducing the length of the RNA hints towards the presence of the phenomenon of protein diffusing over the length of the RNA, as has been reported by Koh *et. al.* earlier using smFRET (Koh et al., 2013). Additionally, the phenomenon of diffusion along the length of RNA has been hinted at in other dsRBPs like Loqs-PD dsRBD (Tants et al., 2017), MLE dsRBD (Ankush Jagtap et al., 2019), DRB4 dsRBD1 (Chiliveri et al., 2017), and RDE4 (Chiliveri and Deshmukh, 2014). To extract the equilibrium dissociation constant (K_d_) for the D12-TRBP-dsRBD2 interaction, residue-wise peak intensities were plotted against the RNA concentration and tried to fit to the binding isotherm for one-site binding (Figure S9). Due to extensive line broadening, there was a lack of data at the inflection point, affecting the data fitting. Hence, the fitted parameters had large errors and remained inconclusive and thus are not reported here.

The ITC-based study was performed with D12 duplex RNA (Figure 6) to get the equilibrium dissociation constant (K_d_) of the two N-terminal dsRBDs, change in enthalpy upon RNA binding (ΔH), and stoichiometry (n) (Table S2). The differential potential against the time plot indicates that RNA did not get saturated with TRBP2-dsRBD1 (Figure 6A) while it got saturated with TRBP2-dsRBD2 as early as at the eighth injection (Figure 6B). The TRBP2-dsRBD1 data could not be fit conclusively to any binding model (K_d_ has large errors in this data) as depicted in Figure 6C. Our studies showed that TRBP2-dsRBD2 binds to D12 RNA with an average K_d_ of 1.18 ± 0.32 μM (Figure 6D and table S2), while TRBP2-dsRBD1 did not show any significant heat exchange (weak binding) during the titration. The integrated change in injection enthalpy (ΔH) versus the molar ratio of the reactants yielded an average ΔH of –10.12 ± 0.43 kcal/mol, suggesting an enthalpy-driven binding event. The entropy ΔS of the binding event was negative, indicating the absence of hydrophobic interaction between the D10 RNA and dsRBD2. ITC study was repeated for TRBP2-dsRBD1 with as high as 22 folds excess of protein, but RNA remained unsaturated. These results strongly suggest that TRBP2-dsRBD2 binds more efficiently to a small perfect A-form duplex, D12 RNA, than TRBP2-dsRBD1.

### Dynamics perturbations of TRBP2-dsRBD2 at various timescales in the presence of RNA ligand

Minute structural changes in TRBP2-dsRBD2 in the presence of RNA necessitated a deeper insight into the perturbations of conformational dynamics at multiple timescales. Backbone ^1^H-^15^N dynamics measurements on TRBP2-dsRBD2 were carried out in the limiting concentration (RNA:Protein = 0.05) of short duplex D12RNA as an increase in the RNA length and an increase in the RNA concentration, both caused line-broadening impacting dynamics measurements. In the presence of D12 RNA, the average *R*_1_ rates (apo = 1.43 ± 0.05 s^-1^; bound = 1.37 ± 0.07 s^-1^) and nOe values (apo = 0.73 ± 0.03; bound = 0.72 ± 0.076) remained unperturbed, while there was a significant increase in the average *R*_2_ rates (apo = 10.92 ± 0.37 s^-1^; bound = 20.04 ± 0.11 s^-1^) (Figure 3). The apparent increase in the *R*_2_ rates (= *R*_2_* + *R*_ex_) hinted towards a perturbation in the μs-ms timescale dynamics, to which only *R*_2_ rates are sensitive. This phenomenon could be attributed to either an increase in intrinsic *R*_2_* resulting from an increase in the residence time of the RNA on the protein, or an increase in the *R*_ex_ component caused by a chemical exchange between apo- and D12-bound state, or RNA-induced conformational exchange in the protein. Interestingly, such a perturbation in *R*_2_ rates was not found for the dsRBD1 upon RNA-binding (Paithankar et al., 2022). The core N- and C-terminal residues (S151, Q154, Q155, S156, E157, N159, V225, V228, L230, A232), a few residues in the vicinity of loop regions (E177, E183, G185, E191, S209) harbored a lower than average bound-state *R*_2_ rates than the rest of the core indicating a faster motion induced in these residues (Figure 3). On adding RNA, the *R*_2_ rates and nOe values of the N- and C-terminal residues (S156, E157, T227, V228, L230, A232), β1 (E177), the loop 2 region (E183, G185, 189, E191), and the loop 4 region (S209) decreased, indicating further enhanced flexibility in one of the vital RNA-binding region (Figure 3).

The extended model-free analysis of apo and RNA-bound TRBP2-dsRBD2 suggested that the anisotropic (ellipsoid) diffusion model was the best fit for the global motion of them both. The global rotational correlation time (*τ*_c_) of core TRBP2-dsRBD2 in the presence of RNA was 10.9 ns as against 7.3 for the apo-dsRBD2 core, indicating an apparent increase in molecular weight. The overall higher S^2^ values for TRBP2-dsRBD2 indicated a rigidification of the backbone amide vectors in the presence of D12 RNA (Figure S10). A few residues in the α_1_ (E166) and α_2_ (L212, N216, L223) regions exhibited a *R*_ex_ component > 1 s^-1^, lying in the vicinity of the reported RNA-binding regions 1 and 3. The number of residues with a significant (> 2 s^-1^) *R*_ex_ — implying the presence of μs-ms timescale motions — increased in the case of bound-state. *R_ex_* was found to be induced in the α_1_ (V168), β_1_ (Y178), β_2_ (T195), β_3_ (I204), and α_2_ (N216, L223) region, thereby indicating that not only the RNA-binding residues but the rest of the core might play a role while interacting with dsRNA substrates. The presence of *R*_ex_ at multiple sites (in addition to RNA-binding regions) rules out the chemical exchange between apo- and D12-bound state contributing to *R*_ex_, thereby implying a presence of RNA-induced conformational exchange is the predominant contributor to *R*_ex_ and to increased line broadening (or apparent *R*_2_ rates, discussed in the NMR-based titration paragraph).

The effective transverse relaxation rates, *R*_2eff_, for TRBP2-dsRBD2, were plotted against the CPMG frequencies (ϖ_CPMG_) in the presence of D12 RNA (Figure S11). None of the residues in either condition showed effective dispersion in the *R*_2eff_ rates with the increase in ϖ_CPMG_, suggesting motions in the timescale sensitive to this experiment remain absent in D12RNA-bound TRBP2-dsRBD2.

Most importantly, HARD NMR experiments recorded for TRBP2-dsRBD2 in the presence of D12-RNA showed fascinating results (Figure S12 and 4). In the presence of RNA, there was massive induction of 10 μs–10 ms timescale conformational dynamics as reflected by the increase in the number of residues having significant *k*_ex_. The extent of enhancement in *k*_ex_ varied along the backbone of the core protein. For instance, a *k*_ex_ of 5000–50,000 kHz with 10–20% *p*_B_ (green spheres) was observed in α_1_ (G162, L167), loop3 (R200), and α_2_ (A213, A219) and 20–40% *p*_B_ in α_1_ (V168), and β_3_ (E203), depicted by the medium and big green spheres respectively (Figures 4B and 4D). The residues V198 in loop 3 and M221 in the α_2_ region exhibited the presence of the highest frequency of motion with a *k*_ex_ > 50,000 Hz. Intriguingly, among these residues, L167, V168 (α_1_), and A213 (α_2_) lie in the reported RNA-binding regions 1 and 3, respectively. L167 and V168 are adjacent to the key RNA binding residue E165 (region 1), which interacts directly with the RNA minor groove. A213 precedes the important KR-helix motif in the α_2_ region of dsRBD2. The K214 and R215 residues make ionic interactions with the negatively charged phosphate backbone of the RNA major groove. The interaction between the RNA and the RNA-binding residues of the protein might induce conformational exchange within the nearby residues like in the case of G162 (region 1), A219 and M221 (region 2). Additionally, residues like R197 (β_2_); V198 and R200 (L3); E203 and G207 (β_3_) are further from RNA binding regions with *k*_ex_ > 5000 Hz. Thus, the ligand affected not only the RNA-binding residues but the entire protein.

Notable perturbations in terms of Δ*k*_ex_ (*k*_bound_ – *k*_apo_) > 10,000 Hz were mapped on the structure of the protein (Figure 7). There was an enhancement of *k*_ex_ in the residues lying in α_1_ (G162, L167), loop 3 (V198), β_3_ (E203 and G207), and α_2_ (A213, A219) region. Concomitantly, suppression of exchange was observed in β_1_ (V180), loop 3 (R200), and α_2_ (L212, R224) regions. A tantalizing relay of exchange was seen between two residue pairs in close spatial proximity. For instance, exchange was induced in V198 and quenched in R200 in the loop 3 region. Similarly, in the case of A213 and L212 falling in the N-terminal RNA-binding region 3, and A219 and R224 lying in the C-terminal of α_2_, the former of the pairs underwent an increase in *k*_ex_ while the latter observed a decrease. Thus, the enhancement of RNA-induced conformational exchange was accommodated by allosteric quenching of the same. Altogether, there was a significant induction of 10 μs–10 ms timescale motions over the entire backbone of TRBP2-dsRBD2 in the presence of RNA. This induction of motions could be ascribed to either an exchange between the apo- and RNA-bound state of the protein, conformational dynamics in the RNA-bound state of the protein, or the previously reported diffusion motion of the protein on the dsRNA (Koh et al., 2013). Interestingly, in TRBP2-dsRBD2, the extent of dynamics perturbation, in terms of Δ*k*_ex_, was significantly lower (10–50 kHz) than TRBP-dsRBD1 where at least 10 residues underwent a Δ*k*_ex_ > 50 kHz in the presence of small dsRNA. Moreover, a relay of exchange was evident only in loop 3 and α_2_ regions of dsRBD2 and not in α_1_ region, which was observed earlier in dsRBD1. Summing up, in the presence of RNA, dsRBD2 undergoes enhanced conformational exchange in the 10 μs– 10 ms timescale. When compared to dsRBD1, dsRBD2 samples limited conformational space both in the absence and presence of D12-RNA.

**Figure 7:**
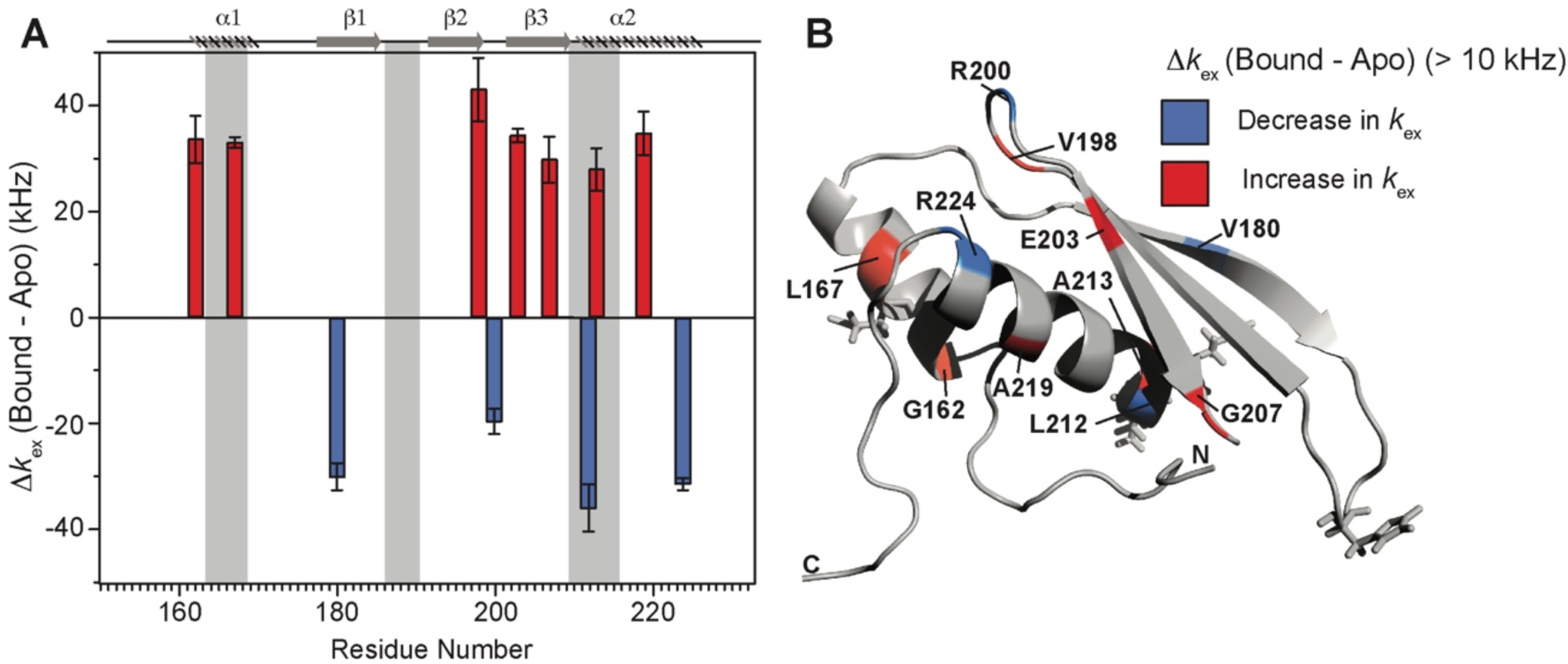
Conformational exchange perturbations in core TRBP2-dsRBD2 in the presence of D12 RNA. (A) Δ*k*_ex_ (D12-bound – apo) TRBP2-dsRBD2 plotted against residue numbers. The secondary structure has been shown on the top, and three RNA-binding regions have been highlighted using vertical grey bars. Only residues having significant perturbation (Δ*k*_ex_ > 10 kHz) have been plotted, where an increase is shown in red, and a decrease is shown in blue, (B) An increase in *k*_ex_ (red) and a decrease (blue) in the presence of D12 RNA indicated on the backbone of the CS-Rosetta structure of apo TRBP2-dsRBD2. The RNA-binding residues have been depicted in stick mode in the tertiary structure.

Thus, our study has led us to propose a model involving two tandem type A dsRBDs in TRBP2. According to our model, these dsRBDs work synergistically to recognize, bind, and assist associated proteins (like Dicer) in processing the incoming dsRNA ligand (Figure 8). The first dsRBD, TRBP2-dsRBD1, has remarkable flexibility, which allows it to recognize a wide range of structurally and sequentially diverse dsRNAs. On the other hand, TRBP2-dsRBD2 firmly adheres to the RNA ligand due to its conserved RNA-binding stretches and relatively high rigidity. During their interaction with the RNA, both dsRBDs experience enhanced conformational changes on a fast timescale (picoseconds to nanoseconds) and a moderate timescale (microseconds to milliseconds), as observed through nuclear spin relaxation and rotating frame relaxation dispersion measurements. These dynamic changes likely enable the dsRBDs to diffuse along the length of the RNA, which explains the broadening of the signals observed during titration with longer RNAs. This diffusion movement is likely crucial for other associated proteins, such as Dicer (Fareh et al., 2016; Lee and Doudna, 2012; Wilson et al., 2015), to carry out precise microRNA biogenesis. By understanding these processes, we can gain insights into the intricate mechanisms involved in RNA processing, which has implications for different biological processes.

**Figure 8:**
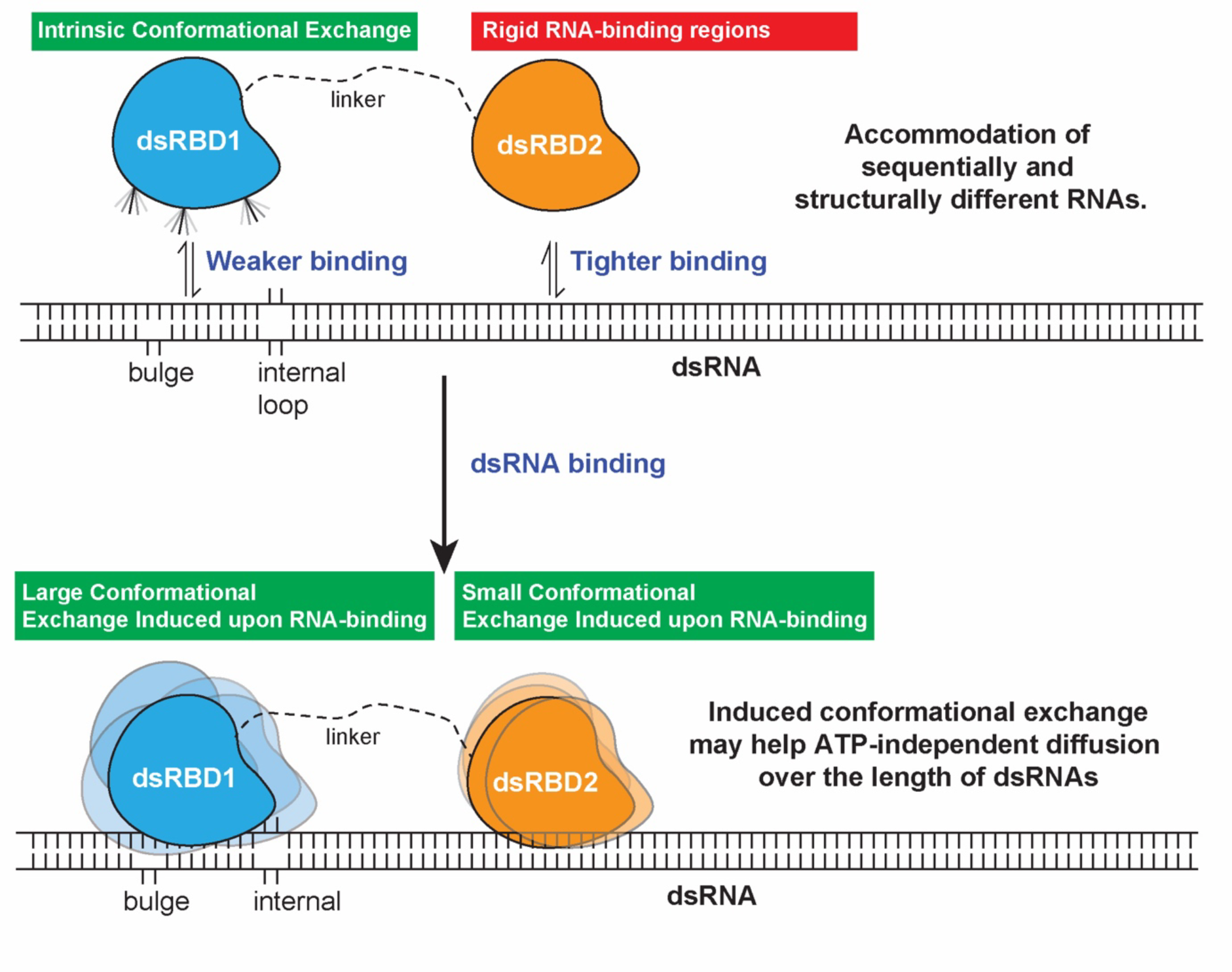
The model proposed for the two type-A dsRBDs in TRBP2 protein. dsRBD2 with rigid and conserved RNA binding regions is able to bind the RNA tightly, whereas dsRBD1 with high conformational exchange is able to recognize different RNA structures (often with bulges and internal loops). Following this, the two dsRBDs upon contacting the RNA undergoes enhanced conformational exchange at different extent. This enhanced conformational exchange coupled with differential binding affinity towards dsRNA might enable the tandem dsRBDs to move along the backbone of the RNA molecule, leading to ATP-independent diffusion.

## Conclusions

It has been long established that despite engaging with a wide array of dsRNA molecules exhibiting diverse structures and sequences, dsRBDs remain unaffected in their own structural conformation. However, the more significant question that has persisted is whether protein dynamics itself plays a crucial role in these intriguing interactions. Though variability in the binding affinity for the two type-A dsRBDs of TRBP has been reported, the specific contribution of conformational dynamics has remained unexplored until now. In the present investigation, we have exclusively delved into the role of conformational dynamics of the two type-A dsRBDs of TRBP2 in RNA recognition and binding. Our findings have unveiled that TRBP2-dsRBD2 samples a limiting conformational space in solution, and it is dsRBD1 that recognizes sequentially and structurally diverse RNA substrates through its high conformational plasticity. While dsRBD1 explores and engages dsRNA via its dynamically interactive RNA-binding surface, dsRBD2 holds the RNA in position via stronger canonical contacts. Once bound to the RNA, the ensuing conformational exchange in both dsRBD1 and dsRBD2 might facilitate the domains to diffuse over the length of the RNA freely, thus playing a pivotal role in assisting Dicer-mediated differential cleavage of RNA. Thus, this study not only adds valuable insights into the mechanics of RNA-protein interactions but also underscores the significance of conformational dynamics in dictating the functional outcome in such intricate biological processes.

Exploring the intricacies of RNA-protein interactions by delving into dynamics-based measurements across diverse RNA shapes within small RNA duplexes could offer invaluable insights into the differential recognition and binding by dsRBD1 and dsRBD2. The experimental challenge lies in the meticulous design of RNA sequences that maintain duplex stability amid the presence of bulges and internal loops of varying lengths and sequences. The task is further complicated by managing RNA length judiciously, as longer sequences can introduce line-broadening in NMR spectroscopy. Navigating these complexities in the future promises to unravel the subtleties of the RNA-protein interactions at a molecule level.

## Materials and Methods

### Protein overexpression and purification

The TRBP2-dsRBD1 (1–105 aa) and TRBP2-dsRBD2 (154–234 aa) cDNA clones were obtained as a kind gift from N. L. Prof. Jennifer Doudna (University of California, Berkeley, CA, USA). The residue numbers in the two constructs (Figure 1A) have been mentioned in reference to the full-length TRBP2 sequence (1–366 aa, Uniprot ID: Q15633-1). The cDNA for TRBP2-dsRBD2 was cloned in pHMGWA vector (Amp^R^), and was expressed as a fusion protein having N-terminal His_6_-Maltose binding protein (MBP) tag-TEV protease cleavage site followed by the protein of interest. Treatment with TEV protease during purification resulted in non-native Ser-Asn-Ala residues at the N-terminal to TRBP2-dsRBD2 (154–234), which were excluded from all the NMR-based dynamics studies. For the NMR experiments, ^15^N-labeled and ^15^N-^13^C labeled (as required) TRBP2-dsRBD2 protein was prepared using ^15^NH_4_Cl and ^13^C-glucose (Cambridge Isotope Laboratories) as a sole source of nitrogen and carbon, respectively, in M9 minimal medium.

TRBP2-dsRBD1 was overexpressed and purified as previously described (Paithankar et al., 2018). Briefly, the pHMGWA plasmid containing the cDNA for the TRBP2-dsRBD2 was transformed into *E. coli* BL21 (DE3) cells, and plated on an LB agar plate (containing 100 μg/mL ampicillin) and incubated at 37 °C for 12 h. A single isolated colony was inoculated into 20 mL LB broth (containing 100 μg/mL ampicillin) and incubated for 12 h at 37 °C, 225 rpm to initiate a starter culture that was eventually used to inoculate 2 L LB broth (containing 100 μg/mL ampicillin) and incubated at 37 °C, 225 rpm till the OD_600_ reached to 0.8–1.0. For induction, IPTG was added at the final concentration of 1 mM, and the culture was further incubated at 37 °C, 225 rpm for another 8 h. The cells were harvested by centrifuging the culture at 4500 *g*, 4 °C for 20 min and resuspended in 25 mL of buffer A (20 mM Tris-Cl, pH 7.5, 500 mM NaCl, 10% glycerol, 5 mM DTT, 10 mM imidazole). To the resuspended cells, lysozyme (Sigma-Aldrich) was added to a final concentration of 50 μg/mL and incubated in ice for 30 min. Post-incubation, 1% Triton X-100, 100 μl of 1 mM PMSF, and 10 X Protease Inhibitor Cocktail (PIC, Roche) (2 mL per 4 g of cell pellet) were added to the cell suspension. The partially lysed cells were sonicated using an ultrasonic sonicator microprobe at 60% amplitude, with 5 s ON and 10 s OFF pulse, for a period of 60 min in an ice bath for complete lysis. The cell lysate was centrifuged at 15,000 x *g* for 2 h, at 4°C, to obtain the total soluble protein (TSP). The TSP was then circulated through a pre-equilibrated (buffer A) Ni-NTA column (HisTrap, 5 ml, GE HealthCare) for 4 h at 4 °C. After equilibrating with the TSP, the column was washed with 40 CVs of buffer A containing 30 mM imidazole to remove the impurities. The fusion protein was eluted with the elution buffer B (buffer A containing 300 mM imidazole). Nucleic acids were removed from the eluted protein using polyethyleneimine (PEI) precipitation. To remove imidazole, the protein solution was dialyzed against cleavage buffer C (20 mM Tris-Cl, pH 7.5, 25 mM NaCl, 10% glycerol, 5 mM DTT). His_6_-MBP-tag was cleaved using TEV protease at a final concentration of 1:100 (protease: protein) at 4 °C for 18 h with intermittent mixing. The completion of the cleavage reaction was tested and confirmed by SDS-PAGE analysis. The cleavage mix was then passed through a sulphopropyl sepharose column (HiPrep SP FF 16/10 20ml, GE HealthCare) and washed with buffer C. TRBP2-dsRBD2 protein was eluted from the cation-exchange column using a salt gradient ranging from 25 mM to 1 M NaCl in the buffer C. To further purify, the protein was subjected to size-exclusion chromatography using sephacryl S-100 HR 16/60 column (GE HealthCare). The final purified protein was concentrated to 1 mM using Amicon (3 kDa cutoff, Merck) and exchanged with NMR buffer D (10 mM sodium phosphate, pH 6.4, 100 mM NaCl, 1 mM EDTA, 5 mM DTT) before recording any experiment.

### Design and preparation of RNA

RNA duplexes miR-16-1-A, miR-16-1-M, miR-16-1-B, and miR-16-1-D were designed and procured as mentioned previously (Paithankar et al., 2022). The shorter 12 bp RNA duplex RNA oligo (D12) was designed from miR-16-1-D and was procured from either Integrated DNA Technologies (Coralville, IA, USA) or GenScript Biotech Corporation (Piscataway NJ, USA). All the RNA sequences used in this study have been listed in Supplementary Table S1.

The preparation of RNA duplexes and the confirmation of their formation was performed, as described previously (Paithankar et al., 2022). Briefly, the respective guide and passenger RNA listed in Supplementary Table S1 were mixed together in a 1:1 ratio, followed by denaturation at 90°C for 10 min and then cooling at 4°C for 30 min. The annealing of all four RNAs (miR-16-1-A, miR-16-1-M, miR-16-1-B, and miR-16-1-D) (Paithankar et al., 2022) and D12 RNA was confirmed by ^1^H NMR by observing the imino proton signals (Figure S1). The annealed samples were maintained in buffer D for all data measurements.

### Size-Exclusion Chromatography – Multiple Angle Light Scattering

Size-exclusion chromatography coupled with multiple angle light scattering (SEC-MALS) experiments were performed using an S75 column (Superdex 75 10/300 GL 24 ml, GE Healthcare), Agilent HPLC system (Wyatt Dawn HELIOs II) and a refractive index detector (Wyatt Optilab T-rEX). The system was first calibrated by injecting 100 μl of 30 μM Bovine Serum Albumin solution (ThermoScientific). Post calibration, 100 μl protein samples were injected (in duplicate) at a concentration of 0.8 mM for TRBP2-dsRBD2. The respective molar mass values of the peaks were calculated using the Zimm model in ASTRA software version 7 (Wyatt Technologies).

### Isothermal titration calorimetric binding assays

All isothermal titration calorimetry (ITC) experiments were performed using a MicroCal PEAQ-ITC calorimeter (Malvern Panalytical, Malvern, UK) operating at 25°C. The final RNAs and protein solutions used for the assays were prepared in buffer D. The D12 dsRNA was used at a concentration of 10 or 20 μM in the sample cell. TRBP2-dsRBD1 concentration was varied from 5– 19 folds of RNA, whereas, in the case of TRBP2-dsRBD2, it was varied from 10–18 folds. The first injection was 0.4 μl (discarded for data analysis), which was followed by eighteen 2 μl injections. All the ITC data was measured in triplicate.

Data were fitted with a single-site binding model using the MicroCal PEAQ-ITC analysis software (Malvern Panalytical, Malvern, UK) to extract the equilibrium dissociation constant (K_d_), stoichiometry (n), and change in enthalpy (ΔH). The final values of the thermodynamic parameters are given as the average of triplicate measurements (Supplementary Table S2).

### NMR Spectroscopy

All the NMR experiments were recorded at 298 K either on: 1) Ascend^TM^ Bruker AVANCE III HD 14.1 Tesla (600 MHz) NMR spectrometer equipped with a quad-channel (^1^H/^13^C/^15^N/^19^F) Cryoprobe (in-house); or on 2) Ascend^TM^ Bruker Avance AV 18.89 Tesla (800 MHz) NMR spectrometer equipped with a triple-channel (^1^H/^13^C/^15^N) Cryoprobe and a Broad Band Inverse probe (located at National Facility for High-Field NMR at TIFR, Mumbai). The ^1^H-^15^N HSQC spectrum was collected with 2048 and 128 points and 12 ppm and 28 ppm spectral width in ^1^H and ^15^N dimensions, respectively, giving an acquisition time of 100 ms in the direct dimension. An inter-scan delay of 1.0 s and 4 scans were used on a 1 mM ^15^N-labeled TRBP2-dsRBD2 sample in a 5 mm Shigemi tube (Shigemi Co., LTD., Tokyo, Japan) (Takeda et al., 2011). ^1^H-^15^N TOCSY-HSQC (mixing times = 60, 80, and 120 ms) and ^1^H-^15^N NOESY-HSQC (mixing times = 150, 300, and 400 ms) were recorded on a 600 MHz NMR spectrometer on ^15^N-labeled TRBP2-dsRBD2 for the backbone assignment of TRBP-dsRBD2. Further, triple resonance experiments like HNCO, HNCACO, HNCA, HNCOCA, HNN, CBCANH, and CBCA(CO)NH were carried out to make the sequential connections using 1.2 mM of ^15^N-^13^C-labeled TRBP2-dsRBD2 sample in a 5 mm Shigemi tube. All the NMR data were processed via TopSpin/NMRPipe (Delaglio et al., 1995) and were analyzed in SPARKY/CARA (Keller, 2004; Lee et al., 2015).

For NMR-based titration assays, ^1^H-^15^N-HSQCs were measured on TRBP2-dsRBD2 (50 μM) with increasing concentrations of duplex RNAs from 0.05**–**0.2 equivalents (in the case of D12 RNA 0.05–5 equivalents) of the protein. After every RNA addition, the protein was allowed to equilibrate for 30 mins before acquiring ^1^H-^15^N HSQC. Peak intensities were plotted against the RNA:protein concentration for each residue and were fit to the one-site binding isotherm using the below equation (Williamson, 2013):

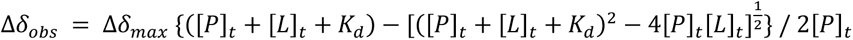

where, Δδ_obs_ is the change in the observed peak intensity from the apo state, Δδ_max_ is the maximum peak intensity change on saturation with ligand, [P]_t_ and [L]_t_ are the total concentration of protein and ligand, respectively, and K_d_ is the equilibrium dissociation constant.

Apo-TRBP-dsRBD2 nuclear spin relaxation experiments (*R*_1_, *R*_2,_ and [^1^H]-^15^N nOe) were recorded at two different field strengths (600 and 800 MHz NMR spectrometers) on a 1 mM ^15^N-TRBP2-dsRBD2 sample in a 5 mm Shigemi tube. ^15^N longitudinal relaxation rates (*R*_1_) were recorded with 8 inversion recovery delays of 10, 30*, 50, 100, 200, 300, 450*, and 600 ms. ^15^N transverse relaxation rates (*R*_2_) were recorded with 8 CPMG (Carr-Purcell-Meiboom-Gill) delays of 17, 34*, 51, 68, 85, 102, 136*, and 170 ms. Steady-state [^1^H]-^15^N heteronuclear nOe experiments were recorded with and without ^1^H saturation with a relaxation delay of 5 s.

All D12-bound TRBP-dsRBD2 nuclear spin relaxation experiments were recorded in a similar fashion as the apo-protein on a 1 mM ^15^N-TRBP2-dsRBD2 in the presence of 50 μM D12 RNA in a 3 mm NMR tube. The ^15^N-*R*_1_ rates were measured with 5 inversion recovery delays of 10, 30*, 70, 150, and 600 ms on 600 MHz spectrometer and 10, 70*, 150, 300, and 600* ms on 800 MHz spectrometer. The ^15^N-*R*_2_ rates were measured with 5 CPMG delays of 17, 51, 85, 136*, and 170 ms on 600 MHz spectrometer and with 4 CPMG delays of 17, 34*, 51, and 68 ms on 800 MHz spectrometer with a CPMG loop length of 17 ms.

For ^15^N relaxation dispersion measurement, a constant time CPMG (Carr-Purcell-Meiboom-Gill) experiment (Tollinger et al., 2001) was recorded on a 600 MHz NMR spectrometer. CPMG relaxation dispersion experiments for apo and D12-bound ^15^N-TRBP2-dsRBD2 were acquired separately, consisting of 15 data points with *v_cpmg_* values of 25, 50, 75, 125*, 175, 275, 375*, 525, 675, 825*, and 1000 Hz at a constant relaxation time – T*_relax_* (40 ms).

Heteronuclear Adiabatic Relaxation Dispersion (HARD) experiments (Mangia et al., 2010; Traaseth et al., 2012) were recorded on apo and D12-bound ^15^N-TRBP2-dsRBD2 on the 600 MHz NMR spectrometer, as described previously (Paithankar et al., 2022). The relaxation delays used for *R*_1π_ were 0, 16, 32, 64, 96, and 128 ms for apo-; and 0, 16, 32, 64, and 112 ms for D12-bound ^15^N TRBP2-dsRBD2. For *R*_2π_ experiments, the relaxation delays used were 0, 16, 32, and 64 ms for apo and 0, 16, and 32 ms for D12-bound ^15^N-TRBP2-dsRBD2. *R*_1_ experiments were acquired similarly to *R*_1π_ and *R*_2π_ experiments without using the adiabatic pulse during evolution. The delays used for the *R*_1_ experiment were 16, 48, 96, 160, 224, 320, 480, and 640 ms for both apo- and RNA-bound protein samples.

All relaxation experiments were measured using a single scan interleaving method, and the order of delays was randomized. The time periods marked with an asterisk have been recorded in duplicates for error estimation. An inter-scan delay of 2.5 s was used for all the above-mentioned relaxation experiments (unless mentioned otherwise).

### NMR Relaxation Data Analysis

NMR spectra were processed with either TopSpin 3.5pl6 or NMRPipe and visualized through SPARKY (version 3.115). For the backbone resonance assignment, CARA (http://cara.nmr.ch/doku.php) was used. All the input NMR spectra were fed in the form of either ucsf or 3rrr file formats. Manual peak picking was done for double and triple-resonance NMR experiments using a synchroscope in CARA. Further identification, confirmation, and assignment of residues were done in a stripscope mode. The chemical shifts of the assigned residues were used as inputs in the CS-Rosetta program in the BMRB server (https://csrosetta.bmrb.wisc.edu/csrosetta/submit) to get an ensemble of the 10 lowest energy structures the structure.

Different relaxation rates like *R_1_*/*R_2_*/*R_1ρ_*/*R_2ρ_* were calculated using mono-exponential decay fitting of peak height against the corresponding set of relaxation delays in a Mathematica script (Spyracopoulos, 2006). Steady-state [^1^H]-^15^N nOes for individual residues were calculated as a ratio of the corresponding residue peak height in the spectra recorded in the presence and absence of ^1^H saturation.

Additional analysis of ^15^N-relaxation data (*R_1_*, *R_2_*, [^1^H]-^15^N nOe) was conducted using the extended model-free formalism via Relax v4.0.3 software (Morin et al., 2014) in a similar fashion carried out previously (Paithankar et al., 2022). The structure of TRBP-dsRBD2 available with PDB ID 2CPN was employed for this analysis.

The effective transverse relaxation rates (R*_2eff_*) from CPMG relaxation dispersion experiments, the *R_1_*, *R_1ρ_*, and *R_2ρ_* relaxation rates obtained from the HARD experiments, the rotational correlation time, and the subsequent *k*_ex_, and *p*_A_ were calculated by fitting the *R_1_*, *R_1ρ_*, and *R_2ρ_* data to a two-state model using numerical fittings as described previously (Paithankar et ^a^l., 2022^;^ Spyracopoulos, 2006). The errors in different fitted parameters were obtained using 500 Monte-Carlo simulations in addition to the duplicate relaxation data points.

The relaxation rates have been depicted in different contexts across the manuscript and the figure legends describe the corresponding context. The raw data has been reported in the Supplementary Information as tables.

### Molecular dynamics simulations

All the ensemble cluster analysis was done using UCSF Chimera (Pettersen et al., 2004) on the CS-Rosetta (Lange et al., 2012; Shen et al., 2010) structure of TRBP2-dsRBD2 to select the lowest energy structures. CS-Rosetta structure was used since the previously reported structure (PDB ID: 2CPN) did not have the extended C-terminal sequence used in this study (2CPN has an amino acid length of 150-225, while the construct used in this study is from 154-234). Moreover, the buffer conditions of previously reported structures differed from what is used in this study. The lowest energy structures in the ensemble were used as the seed structure for apo TRBP2-dsRBD2 molecular dynamics (MD) simulation. Briefly, GROMACS V.2019.6 (https://www.gromacs.org) was used for all the atomistic MD simulations. All-atom additive CHARMM36 protein force field was used to build the topologies (Huang et al., 2017), and initial structure solvation was done TIP3P water molecules in a cubic simulation box with a minimum distance of 1.2 nm from protein in all directions. Microsecond timescale simulations (1 μs) were set up in triplicates in a similar way as described previously (Paithankar et al., 2022). All analysis was performed using GROMACS and VMD.

## Acknowledgments

The authors acknowledge N.L. Prof. Jennifer Doudna (University of California, Berkeley) for the TRBP plasmids. The authors also thank Prof. Gianluigi Veglia (University of Minnesota, Minnesota) and Dr. Fa-An Chao (National Cancer Institute, Bethesda, Maryland) for active discussions while setting up HARD experiments and data analysis. F.P. acknowledges Dr. Radha Chauhan, Ms. Jyotsana, and Ms. Sangeeta for help with SEC-MALS experiments. The authors acknowledge the High-Field NMR facility at IISER-Pune (co-funded by DST-FIST and IISER Pune). J.C. acknowledges extramural funding from the Science and Engineering Research Board, Govt. of India (EMR/2015/001966), Department of Biotechnology, Govt. of India (BT/PR24185/BRB/10/1605/2017), and the generous funding from IISER Pune. F.P. is grateful to DBT-JRF, Govt. of India, for providing a fellowship. Z.A. and H.P. thank IISER Pune for the fellowship.

## References

Acevedo R, Orench-Rivera N, Quarles KA, Showalter SA. 2015. Helical Defects in MicroRNA Influence Protein Binding by TAR RNA Binding Protein. Plos One 10:e0116749. doi:10.1371/journal.pone.0116749

Ankush Jagtap PK, Müller M, Masiewicz P, von Bülow S, Hollmann NM, Chen P-C, Simon B, Thomae AW, Becker PB, Hennig J. 2019. Structure, dynamics and roX2-lncRNA binding of tandem double-stranded RNA binding domains dsRBD1,2 of Drosophila helicase Maleless. Nucleic Acids Res 47:gkz125-. doi:10.1093/nar/gkz125

Auweter SD, Fasan R, Reymond L, Underwood JG, Black DL, Pitsch S, Allain FH. 2006. Molecular basis of RNA recognition by the human alternative splicing factor Fox-1. EMBO J 25:163–173. doi:10.1038/sj.emboj.7600918

Benoit MPMH, Imbert L, Palencia A, Pérard J, Ebel C, Boisbouvier J, Plevin MJ. 2013. The RNA-binding region of human TRBP interacts with microRNA precursors through two independent domains. Nucleic Acids Res 41:4241–4252. doi:10.1093/nar/gkt086

Benoit MPMH, Plevin MJ. 2012. Backbone resonance assignments of the micro-RNA precursor binding region of human TRBP. Biomol NMR Assign 7:229–233. doi:10.1007/s12104-012-9416-8

Bevilacqua PC, Cech TR. 1996. Minor-Groove Recognition of Double-Stranded RNA by the Double-Stranded RNA-Binding Domain from the RNA-Activated Protein Kinase PKR †. Biochemistry 35:9983–9994. doi:10.1021/bi9607259

Bloembergen N, Purcell EM, Pound RV. 1948. Relaxation Effects in Nuclear Magnetic Resonance Absorption. Phys Rev 73:679–712. doi:10.1103/physrev.73.679

Bou-Nader C, Barraud P, Pecqueur L, Pérez J, Velours C, Shepard W, Fontecave M, Tisné C, Hamdane D. 2019. Molecular basis for transfer RNA recognition by the double-stranded RNA-binding domain of human dihydrouridine synthase 2. Nucleic Acids Res 47:gky1302-. doi:10.1093/nar/gky1302

Bycroft M, Grünert S, Murzin AG, Proctor M, Johnston DS. 1995. NMR solution structure of a dsRNA binding domain from Drosophila staufen protein reveals homology to the N-terminal domain of ribosomal protein S5. EMBO J 14:3563–3571. doi:10.1002/j.1460-2075.1995.tb07362.x

Chao F-A, Byrd RA. 2016. Geometric Approximation: A New Computational Approach To Characterize Protein Dynamics from NMR Adiabatic Relaxation Dispersion Experiments. J Am Chem Soc 138:7337–7345. doi:10.1021/jacs.6b02786

Chendrimada TP, Gregory RI, Kumaraswamy E, Norman J, Cooch N, Nishikura K, Shiekhattar R. 2005. TRBP recruits the Dicer complex to Ago2 for microRNA processing and gene silencing. Nature 436:740–744. doi:10.1038/nature03868

Chiliveri SC, Aute R, Rai U, Deshmukh MV. 2017. DRB4 dsRBD1 drives dsRNA recognition in Arabidopsis thaliana tasi/siRNA pathway. Nucleic Acids Res 45:gkx481-. doi:10.1093/nar/gkx481

Chiliveri SC, Deshmukh MV. 2014. Structure of RDE-4 dsRBDs and mutational studies provide insights into dsRNA recognition in the Caenorhabditis elegans RNAi pathway. Biochem J 458:119–130. doi:10.1042/bj20131347

Daubner GM, Cléry A, Allain FH-T. 2013. RRM–RNA recognition: NMR or crystallography…and new findings. Curr Opin Struct Biol 23:100–108. doi:10.1016/j.sbi.2012.11.006

Daviet L, Erard M, Dorin D, Duarte M, Vaquero C, Gatignol A. 2000. Analysis of a binding difference between the two dsRNA-binding domains in TRBP reveals the modular function of a KR-helix motif. Eur J Biochem 267:2419–2431. doi:10.1046/j.1432-1327.2000.01256.x

Delaglio F, Grzesiek S, Vuister GW, Zhu G, Pfeifer J, Bax A. 1995. NMRPipe: A multidimensional spectral processing system based on UNIX pipes. J Biomol NMR 6:277–293. doi:10.1007/bf00197809

Dreyfuss G, Kim VN, Kataoka N. 2002. Messenger-RNA-binding proteins and the messages they carry. Nat Rev Mol Cell Biol 3:195–205. doi:10.1038/nrm760

Eckmann CR, Jantsch MF. 1997. Xlrbpa, a Double-stranded RNA-binding Protein Associated with Ribosomes and Heterogeneous Nuclear RNPs. J Cell Biol 138:239–253. doi:10.1083/jcb.138.2.239

Fareh M, Yeom K-H, Haagsma AC, Chauhan S, Heo I, Joo C. 2016. TRBP ensures efficient Dicer processing of precursor microRNA in RNA-crowded environments. Nat Commun 7:13694. doi:10.1038/ncomms13694

Fierro-Monti I, Mathews MB. 2000. Proteins binding to duplexed RNA: one motif, multiple functions. Trends Biochem Sci 25:241–246. doi:10.1016/s0968-0004(00)01580-2

Fu Y, Zhao X, Li Z, Wei J, Tian Y. 2016. Splicing variants of ADAR2 and ADAR2-mediated RNA editing in glioma. Oncol Lett 12:788–792. doi:10.3892/ol.2016.4734

Gatignol A, Buckler-White A, Berkhout B, Jeang K-T. 1991. Characterization of a Human TAR RNA-Binding Protein That Activates the HIV-1 LTR. Science 251:1597–1600. doi:10.1126/science.2011739

Green SR, Manche L, Mathews MB. 1995. Two Functionally Distinct RNA-Binding Motifs in the Regulatory Domain of the Protein Kinase DAI. Mol Cell Biol 15:358–364. doi:10.1128/mcb.15.1.358

Huang J, Rauscher S, Nawrocki G, Ran T, Feig M, Groot BL de, Grubmüller H, MacKerell AD. 2017. CHARMM36m: an improved force field for folded and intrinsically disordered proteins. Nat Methods 14:71–73. doi:10.1038/nmeth.4067

Huang Y, Ji L, Huang Q, Vassylyev DG, Chen X, Ma J-B. 2009. Structural insights into mechanisms of the small RNA methyltransferase HEN1. Nature 461:823–827. doi:10.1038/nature08433

Imai K, Mitaku S. 2005. Mechanisms of secondary structure breakers in soluble proteins. BIOPHYSICS 1:55–65. doi:10.2142/biophysics.1.55

Jiang S, Baltimore D. 2016. RNA-binding protein Lin28 in cancer and immunity. Cancer Lett 375:108–113. doi:10.1016/j.canlet.2016.02.050

Johnston DS, Brown NH, Gall JG, Jantsch M. 1992. A conserved double-stranded RNA-binding domain. Proc Natl Acad Sci 89:10979–10983. doi:10.1073/pnas.89.22.10979

Keller RLJ. 2004. The Computer Aided Resonance Assignment Tutorial, First. ed. CANTINA Verlag.

Kharrat A, Macias MJ, Gibson TJ, Nilges M, Pastore A. 1995. Structure of the dsRNA binding domain of E. coli RNase III. EMBO J 14:3572–3584. doi:10.1002/j.1460-2075.1995.tb07363.x

Kim VN, Han J, Siomi MC. 2009. Biogenesis of small RNAs in animals. Nat Rev Mol Cell Biol 10:126–139. doi:10.1038/nrm2632

Kim Y, Yeo J, Lee JH, Cho J, Seo D, Kim J-S, Kim VN. 2014. Deletion of Human tarbp2 Reveals Cellular MicroRNA Targets and Cell-Cycle Function of TRBP. Cell Rep 9:1061–1074. doi:10.1016/j.celrep.2014.09.039

Koh HR, Kidwell MA, Ragunathan K, Doudna JA, Myong S. 2013. ATP-independent diffusion of double-stranded RNA binding proteins.

Krieger F, Möglich A, Kiefhaber T. 2005. Effect of Proline and Glycine Residues on Dynamics and Barriers of Loop Formation in Polypeptide Chains. J Am Chem Soc 127:3346–3352. doi:10.1021/ja042798i

Krovat BC, Jantsch MF. 1996. Comparative Mutational Analysis of the Double-stranded RNA Binding Domains of Xenopus laevis RNA-binding Protein A*. J Biol Chem 271:28112–28119. doi:10.1074/jbc.271.45.28112

Lange OF, Rossi P, Sgourakis NG, Song Y, Lee H-W, Aramini JM, Ertekin A, Xiao R, Acton TB, Montelione GT, Baker D. 2012. Determination of solution structures of proteins up to 40 kDa using CS-Rosetta with sparse NMR data from deuterated samples. Proc Natl Acad Sci 109:10873–10878. doi:10.1073/pnas.1203013109

Laraki G, Clerzius G, Daher A, Melendez-Peña C, Daniels S, Gatignol A. 2008. Interactions between the double-stranded RNA-binding proteins TRBP and PACT define the Medipal domain that mediates protein-protein interactions. RNA Biol 5:92–103. doi:10.4161/rna.5.2.6069

Lee HY, Doudna JA. 2012. TRBP alters human precursor microRNA processing in vitro. RNA 18:2012–2019. doi:10.1261/rna.035501.112

Lee W, Tonelli M, Markley JL. 2015. NMRFAM-SPARKY: enhanced software for biomolecular NMR spectroscopy. Bioinformatics 31:1325–1327. doi:10.1093/bioinformatics/btu830

Leuschner PJF, Ameres SL, Kueng S, Martinez J. 2006. Cleavage of the siRNA passenger strand during RISC assembly in human cells. EMBO Rep 7:314–320. doi:10.1038/sj.embor.7400637

Mangia S, Traaseth NJ, Veglia G, Garwood M, Michaeli S. 2010. Probing Slow Protein Dynamics by Adiabatic R 1ρ and R 2ρ NMR Experiments. J Am Chem Soc 132:9979–9981. doi:10.1021/ja1038787

Masliah G, Barraud P, Allain FH-T. 2012. RNA recognition by double-stranded RNA binding domains: a matter of shape and sequence. Cell Mol Life Sci 70:1875–1895. doi:10.1007/s00018-012-1119-x

Masliah G, Maris C, König SL, Yulikov M, Aeschimann F, Malinowska AL, Mabille J, Weiler J, Holla A, Hunziker J, Meisner-Kober N, Schuler B, Jeschke G, Allain FH. 2018. Structural basis of siRNA recognition by TRBP double-stranded RNA binding domains. EMBO J 37:e97089. doi:10.15252/embj.201797089

Matranga C, Tomari Y, Shin C, Bartel DP, Zamore PD. 2005. Passenger-Strand Cleavage Facilitates Assembly of siRNA into Ago2-Containing RNAi Enzyme Complexes. Cell 123:607– 620. doi:10.1016/j.cell.2005.08.044

Morin S, Linnet TE, Lescanne M, Schanda P, Thompson GS, Tollinger M, Teilum K, Gagné S, Marion D, Griesinger C, Blackledge M, d’Auvergne EJ. 2014. relax: the analysis of biomolecular kinetics and thermodynamics using NMR relaxation dispersion data. Bioinformatics 30:2219–2220. doi:10.1093/bioinformatics/btu166

Nanduri S, Rahman F, Williams BRG, Qin J. 2000. A dynamically tuned double-stranded RNA binding mechanism for the activation of antiviral kinase PKR. EMBO J 19:5567–5574. doi:10.1093/emboj/19.20.5567

Noland CL, Doudna JA. 2013. Multiple sensors ensure guide strand selection in human RNAi pathways. RNA 19:639–648. doi:10.1261/rna.037424.112

Noland CL, Ma E, Doudna JA. 2011. siRNA Repositioning for Guide Strand Selection by Human Dicer Complexes. Mol Cell 43:110–121. doi:10.1016/j.molcel.2011.05.028

Paithankar H, Jadhav PV, Naglekar AS, Sharma S, Chugh J. 2018. 1H, 13C and 15N resonance assignment of domain 1 of trans-activation response element (TAR) RNA binding protein isoform 1 (TRBP2) and its comparison with that of isoform 2 (TRBP1). Biomol NMR Assign 12:189–194. doi:10.1007/s12104-018-9807-6

Paithankar H, Tarang GS, Parvez F, Marathe A, Joshi M, Chugh J. 2022. Inherent conformational plasticity in dsRBDs enables interaction with topologically distinct RNAs. Biophys J 121:1038– 1055. doi:10.1016/j.bpj.2022.02.005

Peters GA, Hartmann R, Qin J, Sen GC. 2001. Modular Structure of PACT: Distinct Domains for Binding and Activating PKR. Mol Cell Biol 21:1908–1920. doi:10.1128/mcb.21.6.1908-1920.2001

Pettersen EF, Goddard TD, Huang CC, Couch GS, Greenblatt DM, Meng EC, Ferrin TE. 2004. UCSF Chimera—A visualization system for exploratory research and analysis. J Comput Chem 25:1605–1612. doi:10.1002/jcc.20084

Ramos A, Grünert S, Adams J, Micklem DR, Proctor MR, Freund S, Bycroft M, Johnston DS, Varani G. 2000. RNA recognition by a Staufen double-stranded RNA-binding domain. EMBO J 19:997–1009. doi:10.1093/emboj/19.5.997

Shen Y, Bryan PN, He Y, Orban J, Baker D, Bax A. 2010. De novo structure generation using chemical shifts for proteins with high-sequence identity but different folds. Protein Sci 19:349–356. doi:10.1002/pro.303

Spyracopoulos L. 2006. A suite of Mathematica notebooks for the analysis of protein main chain 15N NMR relaxation data. J Biomol NMR 36:215. doi:10.1007/s10858-006-9083-0

Takeda M, Hallenga K, Shigezane M, Waelchli M, Löhr F, Markley JL, Kainosho M. 2011. Construction and performance of an NMR tube with a sample cavity formed within magnetic susceptibility-matched glass. J Magn Reson 209:167–173. doi:10.1016/j.jmr.2011.01.005

Tants J-N, Fesser S, Kern T, Stehle R, Geerlof A, Wunderlich C, Juen M, Hartlmüller C, Böttcher R, Kunzelmann S, Lange O, Kreutz C, Förstemann K, Sattler M. 2017. Molecular basis for asymmetry sensing of siRNAs by the Drosophila Loqs-PD/Dcr-2 complex in RNA interference. Nucleic Acids Res 45:gkx886-. doi:10.1093/nar/gkx886

Tian B, Bevilacqua PC, Diegelman-Parente A, Mathews MB. 2004. The double-stranded-RNA-binding motif: interference and much more. Nat Rev Mol Cell Biol 5:1013–1023. doi:10.1038/nrm1528

Tollinger M, Skrynnikov NR, Mulder FAA, Forman-Kay JD, Kay LE. 2001. Slow Dynamics in Folded and Unfolded States of an SH3 Domain. J Am Chem Soc 123:11341–11352. doi:10.1021/ja011300z

Traaseth NJ, Chao F-A, Masterson LR, Mangia S, Garwood M, Michaeli S, Seelig B, Veglia G. 2012. Heteronuclear Adiabatic Relaxation Dispersion (HARD) for quantitative analysis of conformational dynamics in proteins. J Magn Reson 219:75–82. doi:10.1016/j.jmr.2012.03.024

Ucci JW, Kobayashi Y, Choi G, Alexandrescu AT, Cole JL. 2007. Mechanism of Interaction of the Double-Stranded RNA (dsRNA) Binding Domain of Protein Kinase R with Short dsRNA Sequences †. Biochemistry 46:55–65. doi:10.1021/bi061531o

Wilkinson MF, Shyu A. 2001. Multifunctional regulatory proteins that control gene expression in both the nucleus and the cytoplasm. BioEssays 23:775–787. doi:10.1002/bies.1113

Williamson MP. 2013. Using chemical shift perturbation to characterise ligand binding. Prog Nucl Magn Reson Spectrosc 73:1–16. doi:10.1016/j.pnmrs.2013.02.001

Wilson RC, Tambe A, Kidwell MA, Noland CL, Schneider CP, Doudna JA. 2015. Dicer-TRBP Complex Formation Ensures Accurate Mammalian MicroRNA Biogenesis. Mol Cell 57:397–407. doi:10.1016/j.molcel.2014.11.030

Wostenberg C, Lary JW, Sahu D, Acevedo R, Quarles KA, Cole JL, Showalter SA. 2012. The Role of Human Dicer-dsRBD in Processing Small Regulatory RNAs. PLoS ONE 7:e51829. doi:10.1371/journal.pone.0051829

Yadav DK, Zigáčková D, Zlobina M, Klumpler T, Beaumont C, Kubíčková M, Vaňáčová Š, Lukavsky PJ. 2019. Staufen1 reads out structure and sequence features in ARF1 dsRNA for target recognition. Nucleic Acids Res 48:2091–2106. doi:10.1093/nar/gkz1163

Yamashita S, Nagata T, Kawazoe M, Takemoto C, Kigawa T, Güntert P, Kobayashi N, Terada T, Shirouzu M, Wakiyama M, Muto Y, Yokoyama S. 2011. Structures of the first and second double-stranded RNA-binding domains of human TAR RNA-binding protein. Protein Sci 20:118–130. doi:10.1002/pro.543

Yan L, Liang M, Hou X, Zhang Y, Zhang H, Guo Z, Jinyu J, Feng Z, Mei Z. 2019. The role of microRNA-16 in the pathogenesis of autoimmune diseases: A comprehensive review. Biomed Pharmacother 112:108583. doi:10.1016/j.biopha.2019.01.044

Yang SW, Chen H-Y, Yang J, Machida S, Chua N-H, Yuan YA. 2010. Structure of Arabidopsis HYPONASTIC LEAVES1 and Its Molecular Implications for miRNA Processing. Structure 18:594–605. doi:10.1016/j.str.2010.02.006

