## Supplementary Information for "Differential conformational dynamics in two type-A RNA-binding domains drive the double-stranded RNA recognition and binding"

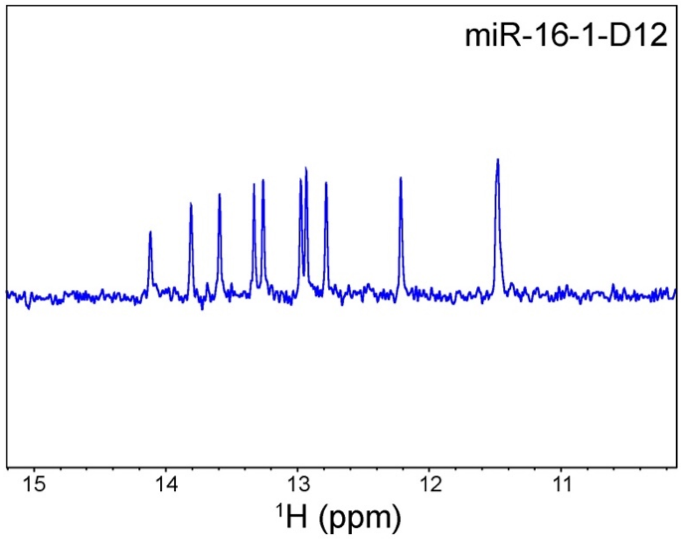


**Figure S1**: $^{1}$H-NMR spectrum of the imine region of the annealed D12 RNA indicating the duplex formation.

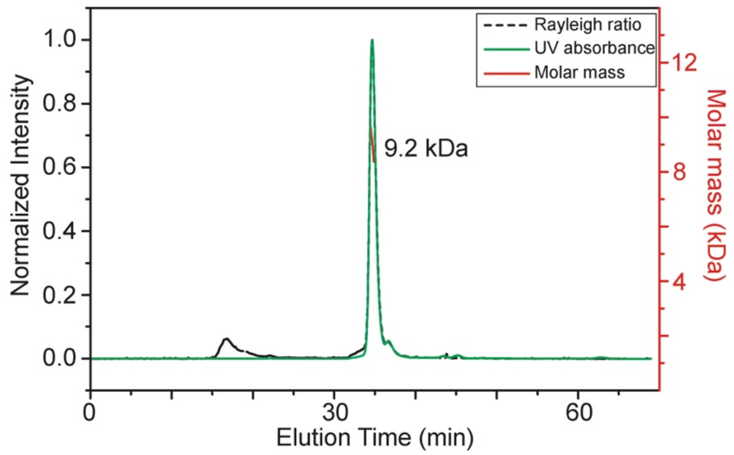


**Figure S2**: SEC-MALS elution profile for TRBP2-dsRDB2 showing that the protein remains monomeric in the buffer conditions used for NMR studies.

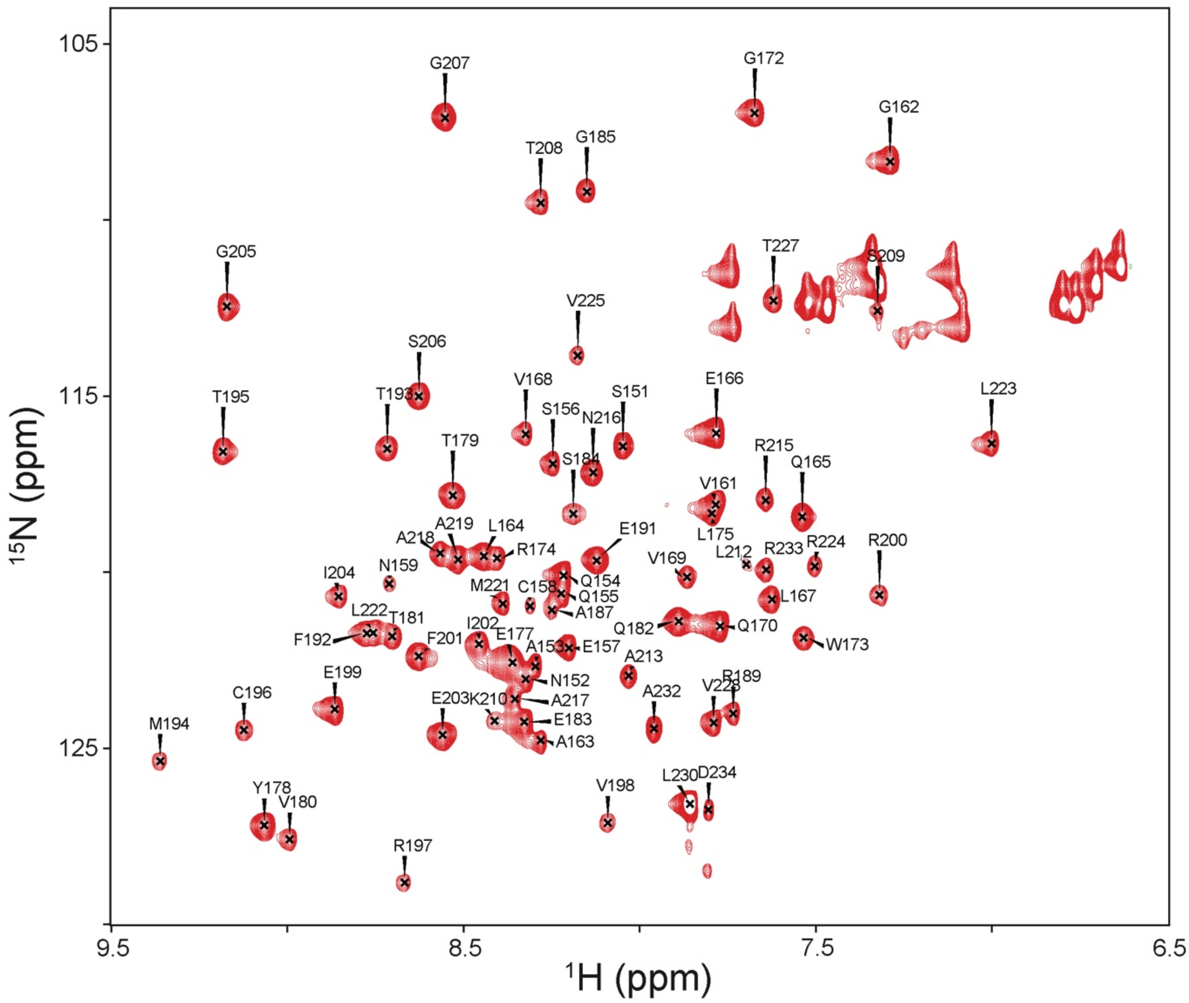


**Figure S3**: Backbone resonance assignments for TRBP2-dsRBD2 marked on the $^{1}H$-$^{15}N$ HSQC recorded on 600 MHz NMR spectrometer at 298 K in buffer D, pH 6.4.

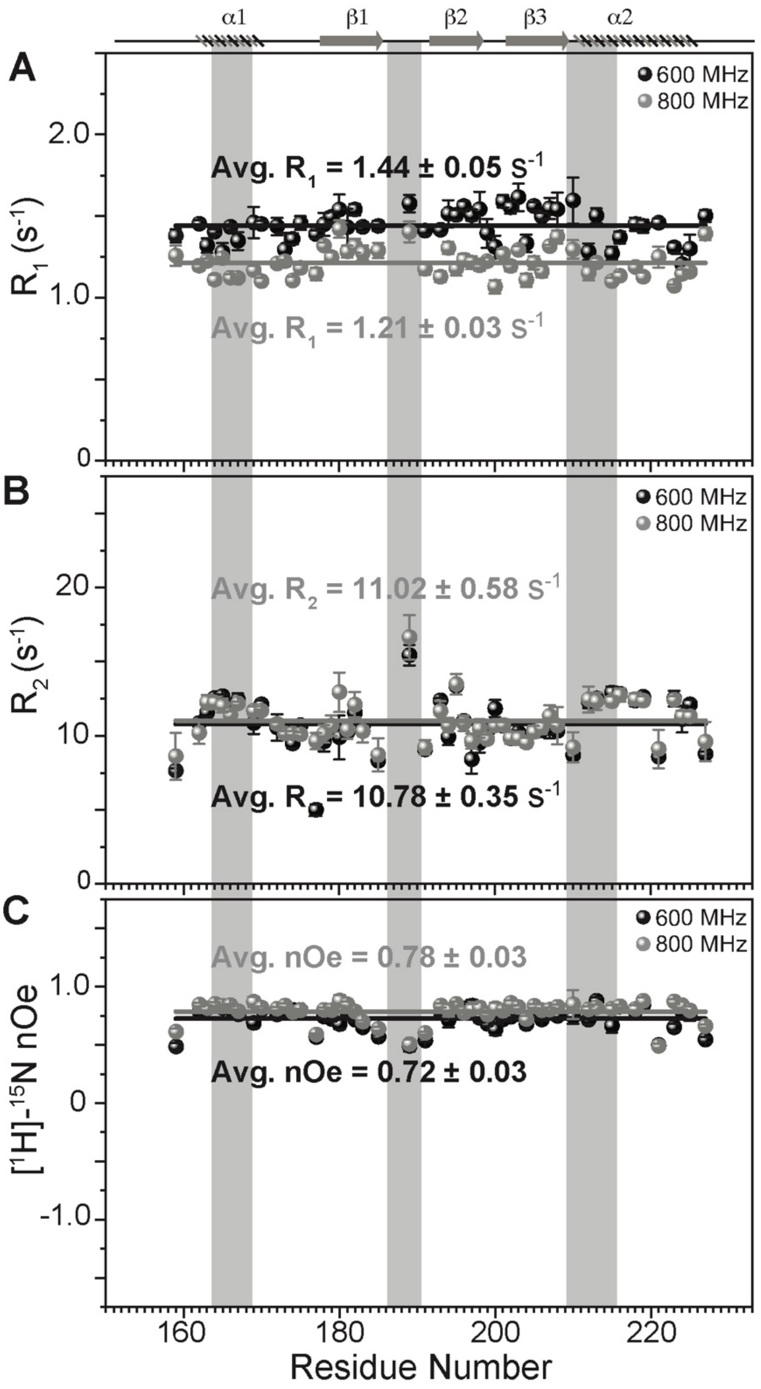


**Figure S4**: A) Longitudinal relaxation rates ($R_1$), B) transverse relaxation rates ($R_2$), and heteronuclear [$^1$H]-$^{15}$N-nOe, as measured for common residues of TRBP2-dsRDB2 on 600 MHz (black) and 800 MHz (grey) magnetic fields at 298 K plotted against residue numbers for both fields. Average $R_1$, $R_2$, and [$^1$H]-$^{15}$N-nOe of the core residues (159-227 aa) at 600 MHz is depicted in the green bar, and at 800 MHz is depicted in the grey bar (average is calculated only for the common residues that could be analyzed between the data measured at two magnetic fields). The secondary structure of the protein has been shown on the top, and three RNA-binding regions have been highlighted using vertical grey bars

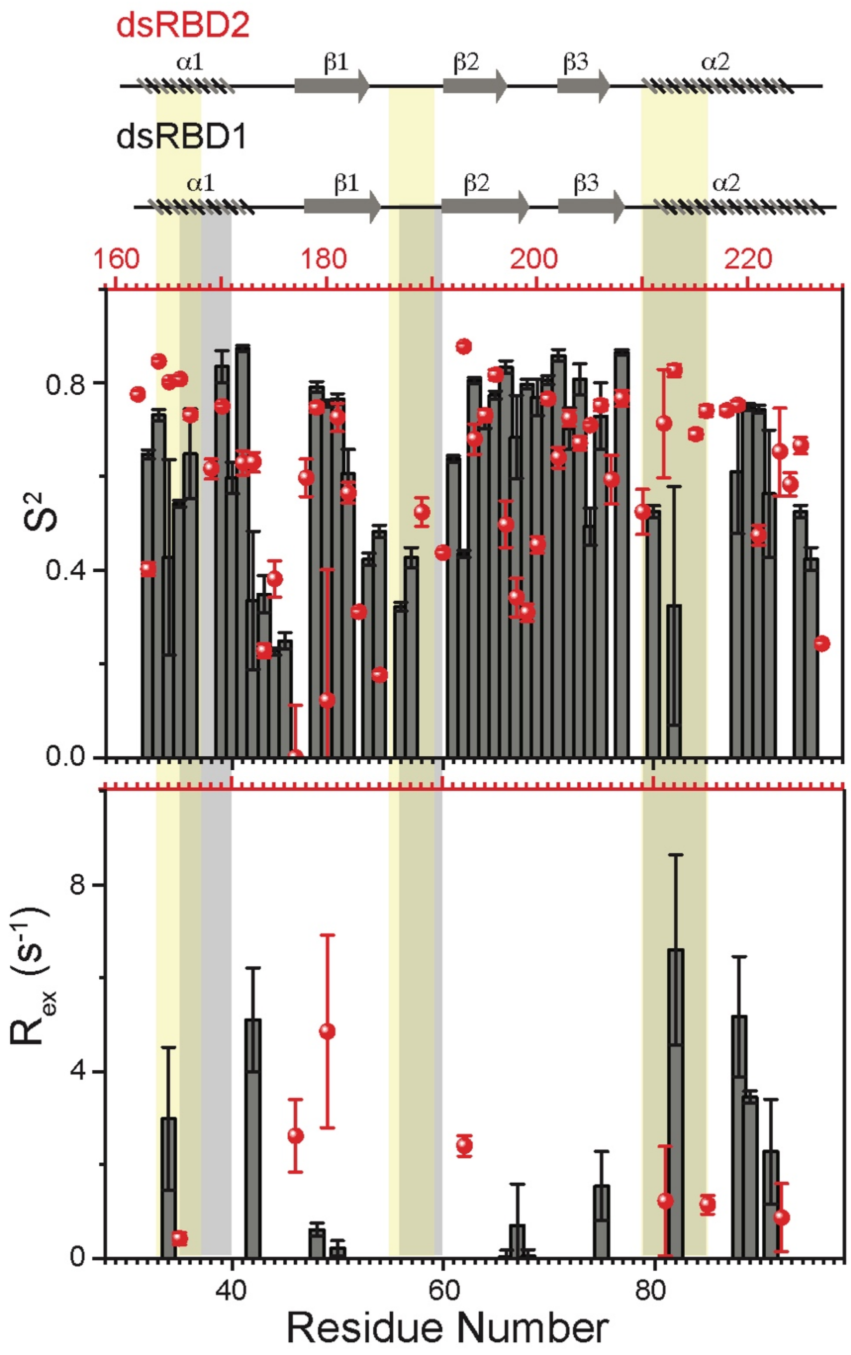


**Figure S5**: Top-panel: Order parameters ($S^2$), and bottom-panel: $R_{ex}$ as calculated using model-free fitting of the fast relaxation data for core residues of TRBP2-dsRBD1 (histograms) and TRBP2-dsRBD2 (red scatter) plotted against residue numbers. The secondary structure of both proteins has been shown on the top, and three RNA-binding regions have been highlighted using vertical grey bars (for TRBP2-dsRBD1) and vertical yellow bars (for TRBP2-dsRBD2).

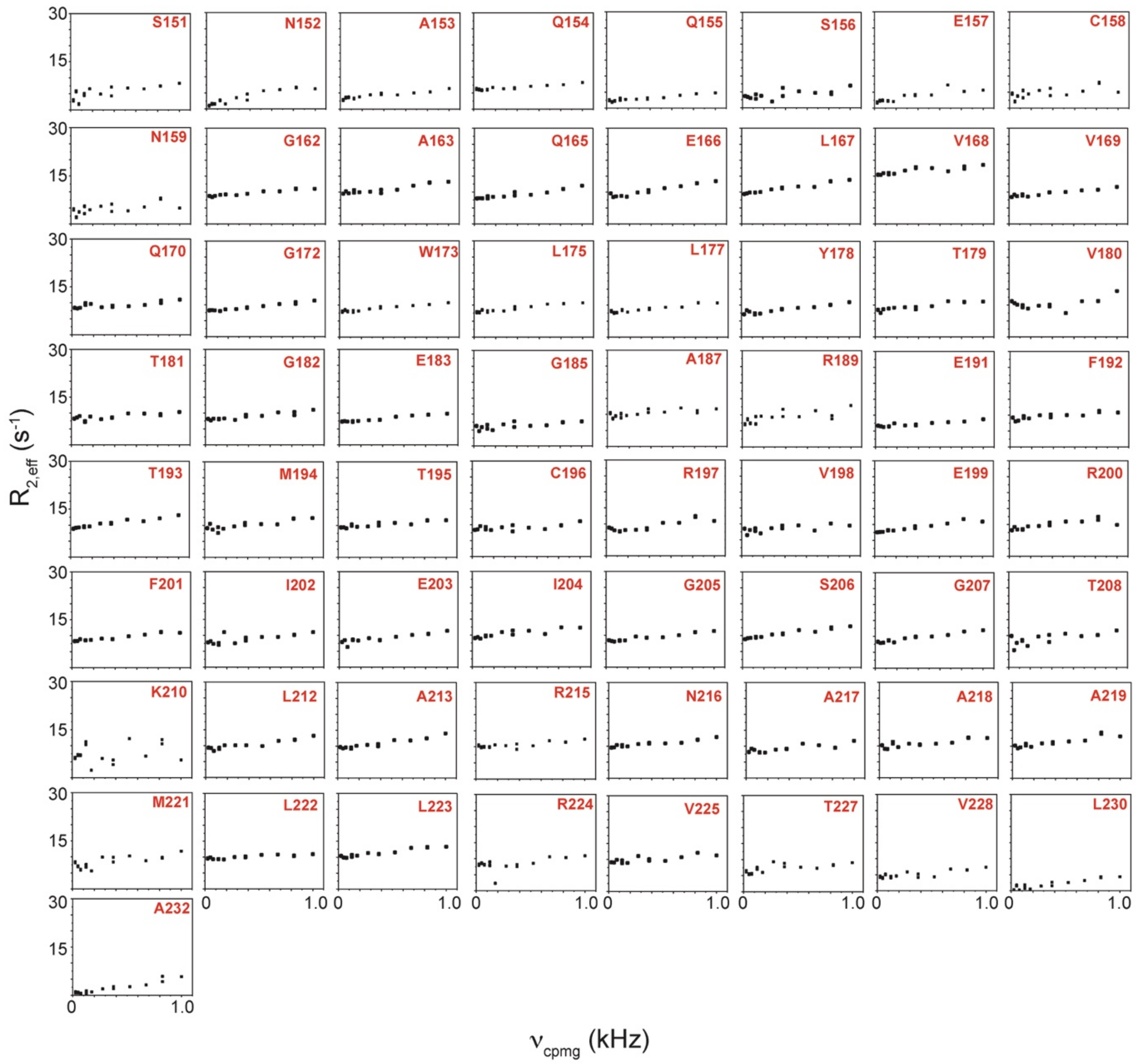


**Figure S6**: $R_{2,\text{eff}}$ rates plotted against $\nu_{\text{CPMG}}$ for the 65 non-overlapping residues of apo TRBP2-dsRBD2 measured at 600 MHz at 298 K. Residue names and respective positions in the TRBP2 protein have been mentioned in each plot.

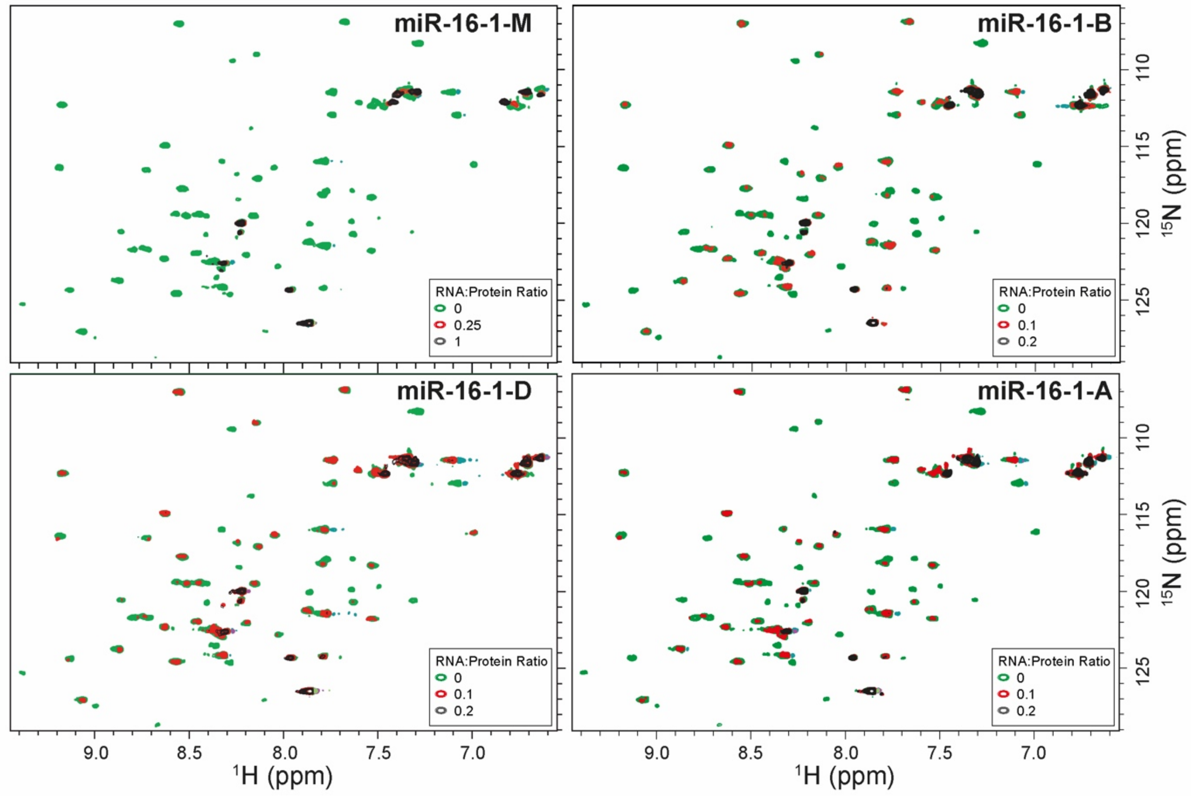


**Figure S7**: Titration of $^{15}$N-TRBP2-dsRBD2 with miR-16-1-M (An overlay of $^{1}$H-$^{15}$N -HSQC spectra of apo-protein (green) RNA:protein (R:P) = 0.25:1 (red) and (R:P) = 1:1 (black)); miR-16-1-B, miR-16-1-D, and miR-16-1-A (wt) [Overlaid $^{1}$H-$^{15}$N -HSQC spectra of apo-protein (green), (R:P) = 0.1:1 (red) and (R:P) = 0.2:1 (black)]. All the data was measured on a 600 MHz NMR spectrometer at 298 K in buffer D, pH 6.4.

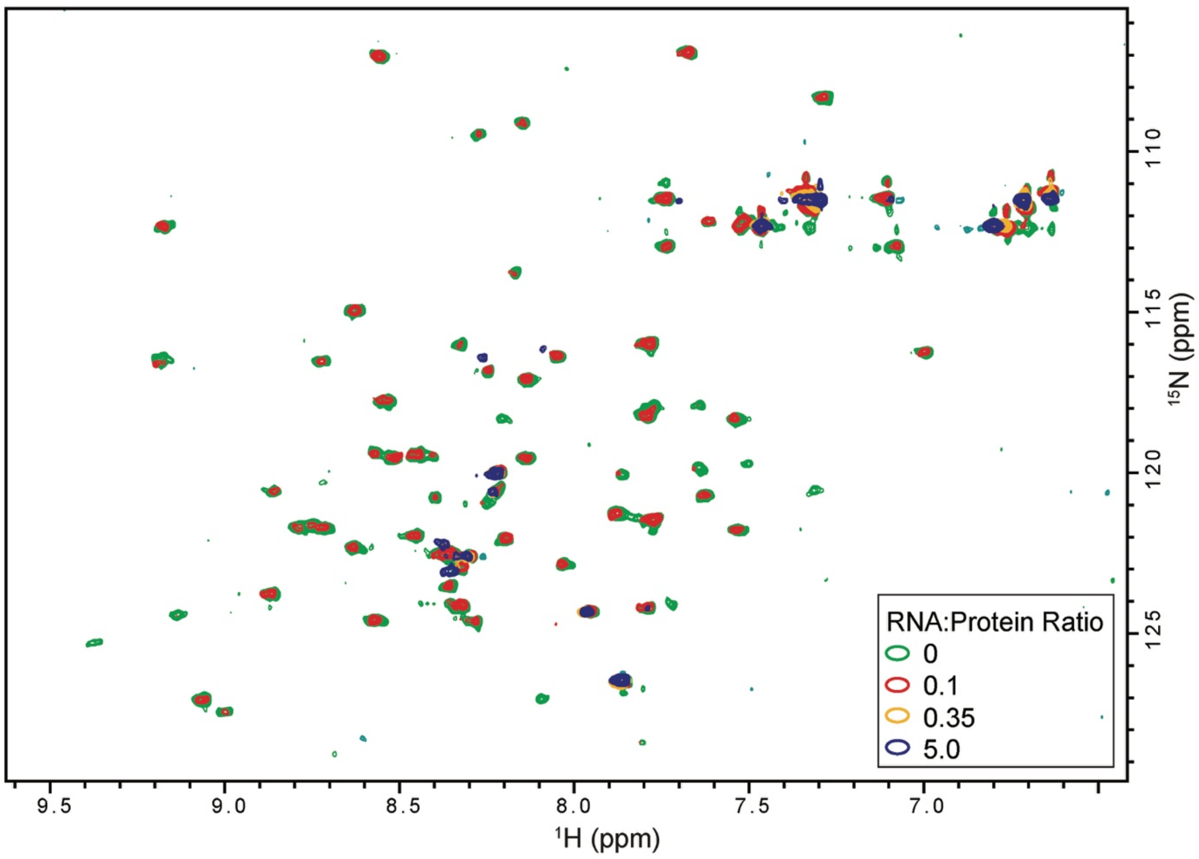


**Figure S8**: Titration of $^{15}$N-TRBP2-dsRBD2 with D12 RNA. An overlay of $^{1}$H-$^{15}$N -HSQC spectra of apo-protein (green), (R:P) = 0.1:1 (red), (R:P) = 0.35:1 (yellow), and (R:P) = 5:1 (blue) measured on 600 MHz NMR spectrometer at 298 K in buffer D, pH 6.4.

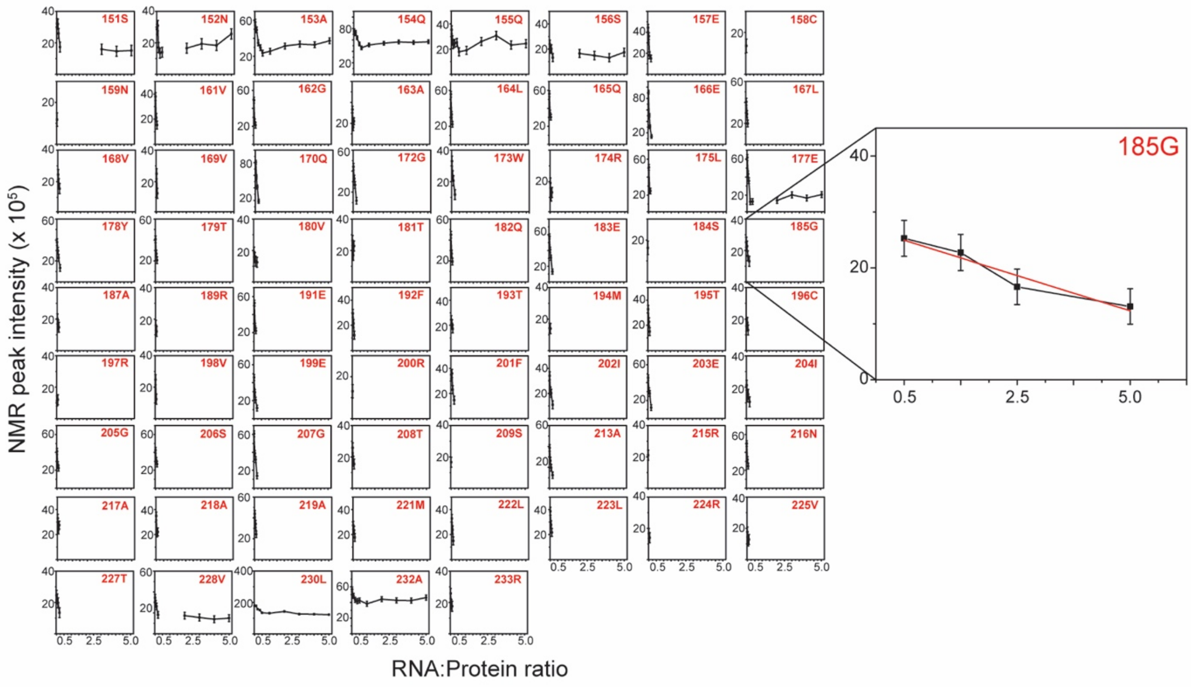


**Figure S9**: Normalized NMR peak intensity from $^{1}H$-$^{15}N$ HSQC spectra of TRBP2-dsRBD2 plotted against protein: D12 RNA ratio from the titration carried out in Figure S6. An example residue has been zoomed in to highlight the intensity decay.

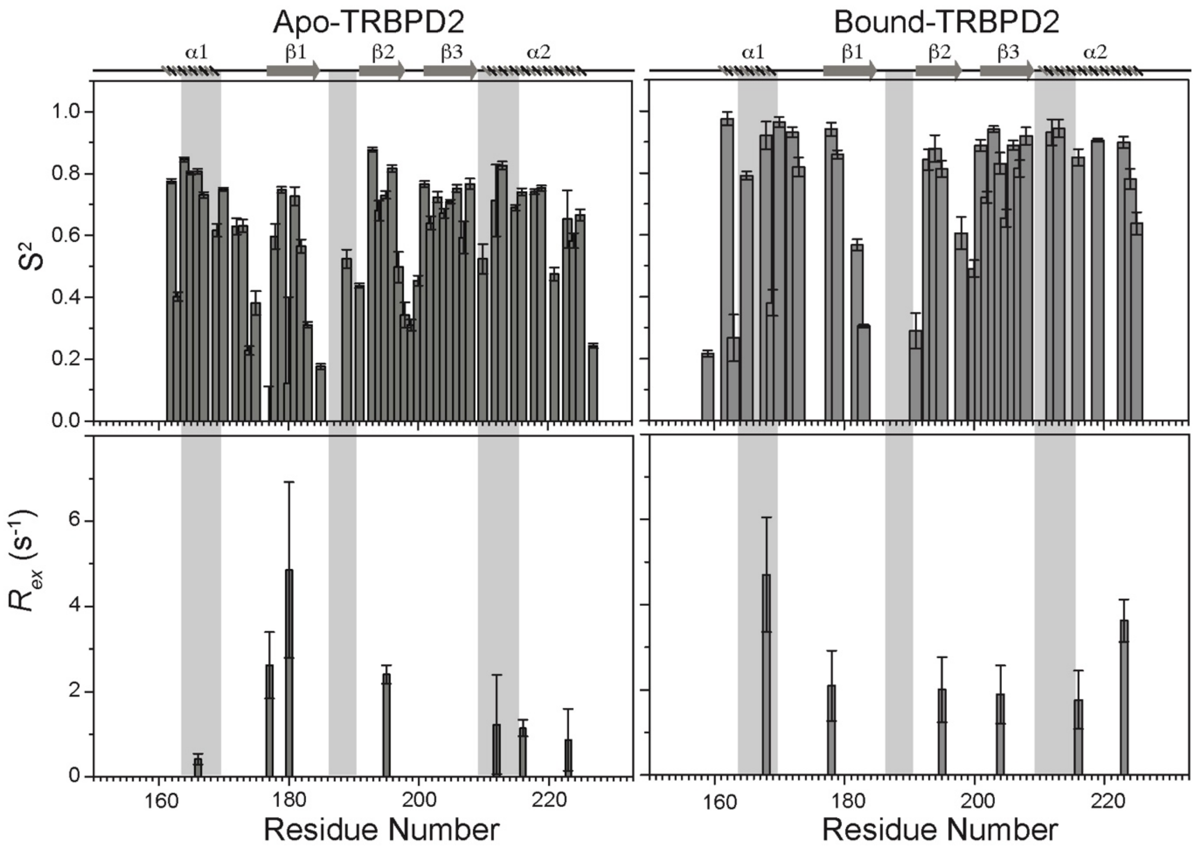


**Figure S10**: Top-panel: Order parameters ($S^2$), and bottom-panel: $R_{ex}$ as calculated using model-free fitting of the fast relaxation data for apo TRBP2-dsRBD2 (Left-panel) and D12 RNA-bound TRBP2-dsRBD2 (Right-panel) plotted against residue numbers. The secondary structure has been shown on the top, and three RNA-binding regions have been highlighted using vertical grey bars.

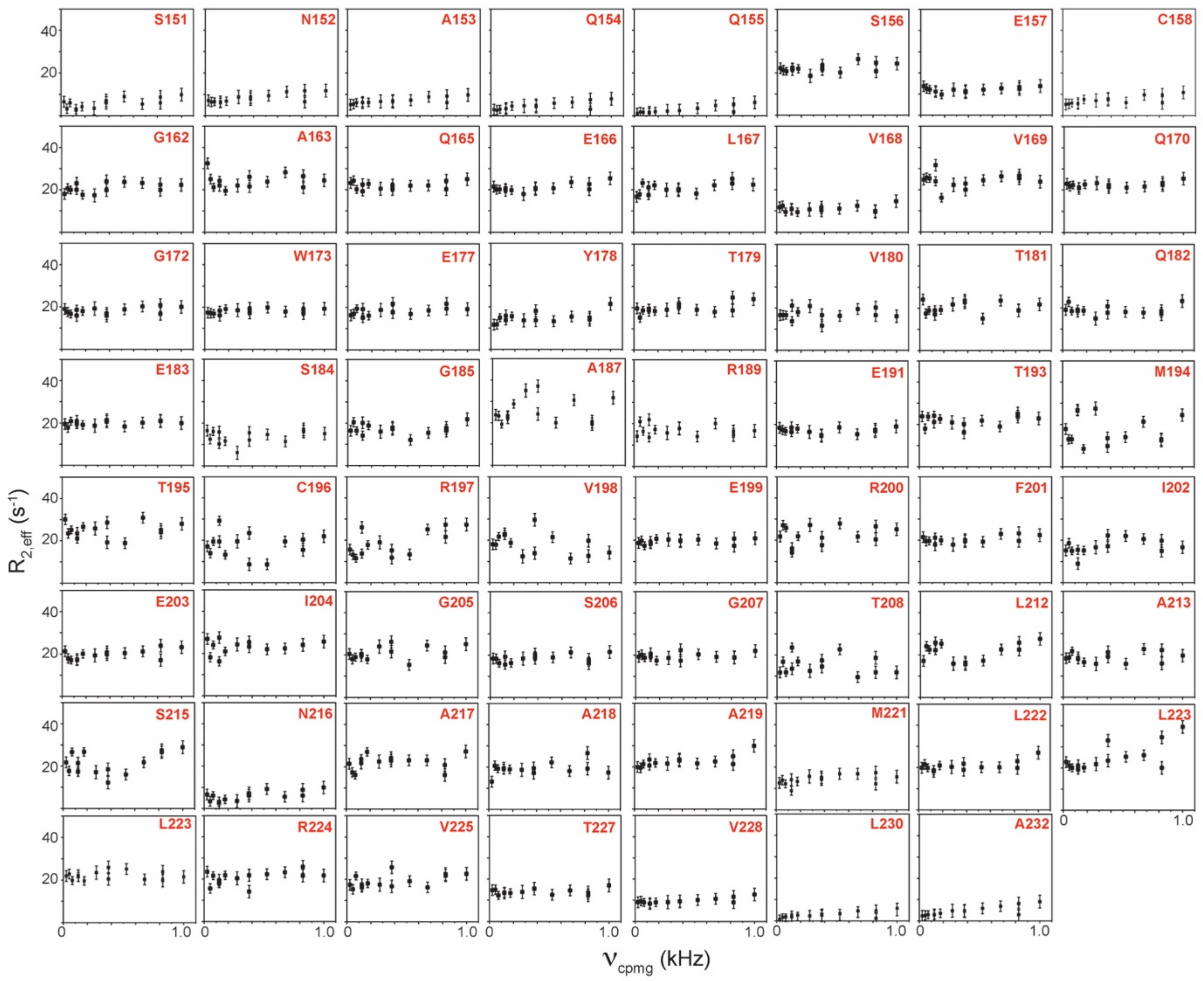


**Figure S11**: $R_{2,\text{ eff}}$ rates plotted against $\nu_{CPMG}$ for the 63 non-overlapping residues of TRBP2-dsRBD2 measured in the presence of D12 RNA at 600 MHz at 298 K. Residue names, along with respective positions in the TRBP2 protein, have been mentioned in each plot.

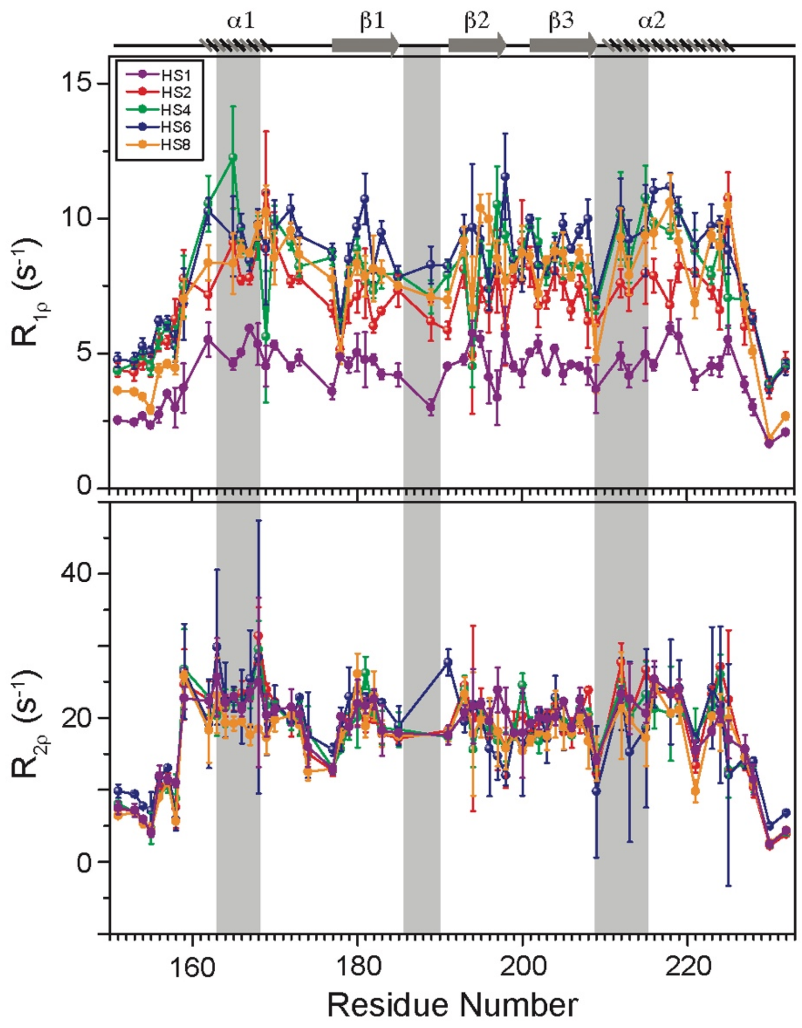


**Figure S12**: Heteronuclear Adiabatic Relaxation Dispersion (HARD) experiments recorded on $^{15}N$-TRBP2-dsRBD2 in the presence of D12-RNA on the 600 MHz NMR spectrometer at 298 K. Top-panel: $R_{1\rho}$, and Bottom-panel: $R_{2\rho}$ rates plotted against residue numbers. Increasing applied spin-lock field strength is denoted by increasing the stretching factor, n (in HSn). The secondary structure has been shown on the top, and three RNA-binding regions have been highlighted using vertical grey bars.

**Table S1**: RNA sequences used to study interaction with dsRBDs.

| Name | Strand | Sequence (5′ → 3′) |
|---|---|---|
| **miR-16-1-A** | Guide | UAGCAGCACGUAAAUAUUGGCG |
| | Passenger | CCAGUAUUAACUGUGCUGCUGAA |
| **miR-16-1-D** | Guide | UAGCAGCACGUAAAUAUUGGCG |
| | Passenger | CCAGUAUUUACGUGCUGCUGAA |
| **miR-16-1-M** | Guide | UAGCAGCACGUAAAUAUUGGCG |
| | Passenger | CCAGUAUUAACGUGCUGCUGAA |
| **miR-16-1-B** | Guide | UAGCAGCACGUAAAUAUUGGCG |
| | Passenger | CCAGUAUUAACGUGCUGCUGAA |
| **D12 RNA** | Guide | CGUAAAUAUUCG |
| | Passenger | CGAGUAUUUACG |

**Table S2**: ITC binding study of TRBP2-dsRBD2 and D12 RNA carried out in triplicate.

| Experiment No. | [Syr] (mM) TRBP2_dsRBD2 | [Cell] (μM) D12 RNA | n (stoichiometry) | $K_d$ (μM) | ΔH (kcal/mol) |
|---|---|---|---|---|---|
| 1) | 1.4 | 10 | 3.03 ± 0.06 | 1.13 ± 0.19 | -9.2 ± 0.23 |
| 2) | 0.8 | 10 | 2.84 ± 0.09 | 0.996 ± 0.31 | -9.16 ± 0.39 |
| 3) | 0.8 | 10 | 2.59 ± 0.11 | 1.41 ± 0.46 | -12 ± 0.67 |
| **Average** | - | - | **2.82 ± 0.09** | **1.18 ± 0.32** | **-10.12 ± 0.43** |

**Table S3**: Amino acid length of different secondary structures in TRBP2-dsRBD1 and TRBP2-dsRBD2 CS-ROSSETA structures. α represents an α-helix, L represents a loop, and β represents the β-strand.

| Domain Construct | α1 | L1 | β1 | L2 | β2 | L3 | β3 | L4 | α2 |
|---|---|---|---|---|---|---|---|---|---|
| TRBP2-dsRBD1 | 10 | 4 | 8 | 5 | 9 | 2 | 7 | 2 | 17 |
| TRBP2-dsRBD2 | 10 | 5 | 8 | 6 | 8 | 4 | 6 | 2 | 15 |

**Assignment Report of TRBP2-dsRBD2**
**(as obtained from CARA in NMR-STAR 3.1 format)**

```
data_starch_output
###################################################################
#       Chemical Shift Ambiguity Index Value Definitions          #
#                                                                 #
# The values other than 1 are used for those atoms with different #
# chemical shifts that cannot be assigned to stereospecific atoms #
# or to specific residues or chains.                              #
#                                                                 #
#   Index Value            Definition                             #
#                                                                 #
#      1             Unique (including isolated methyl protons,   #
#                         geminal atoms, and geminal methyl       #
#                         groups with identical chemical shifts)  #
#                         (e.g. ILE HD11, HD12, HD13 protons)     #
#      2             Ambiguity of geminal atoms or geminal methyl #
#                         proton groups (e.g. ASP HB2 and HB3     #
#                         protons, LEU CD1 and CD2 carbons, or    #
#                         LEU HD11, HD12, HD13 and HD21, HD22,    #
#                         HD23 methyl protons)                    #
#      3             Aromatic atoms on opposite sides of          #
#                         symmetrical rings (e.g. TYR HE1 and HE2 #
#                         protons)                                #
#      4             Intraresidue ambiguities (e.g. LYS HG and    #
#                         HD protons or TRP HZ2 and HZ3 protons)  #
#      5             Interresidue ambiguities (LYS 12 vs. LYS 27) #
#      6             Intermolecular ambiguities (e.g. ASP 31 CA   #
#                         in monomer 1 and ASP 31 CA in monomer 2 #
#                         of an asymmetrical homodimer, duplex    #
#                         DNA assignments, or other assignments   #
#                         that may apply to atoms in one or more  #
#                         molecule in the molecular assembly)     #
#      9             Ambiguous, specific ambiguity not defined    #
#                                                                 #
###################################################################
loop_
    _Atom_chem_shift.Atom_ID
    _Atom_chem_shift.Comp_index_ID
    _Atom_chem_shift.Comp_ID
    _Atom_chem_shift.Atom_ID
    _Atom_chem_shift.Atom_type
    _Atom_chem_shift.Val
    _Atom_chem_shift.Val_err
    _Atom_chem_shift.Ambiguity_code

C              1              SER            C              C
171.651        0.3            1
CA             1              SER            CA             C
59.107         0.3            1
CB             1              SER            CB             C
66.892         0.3            1
H              1              SER            H              H
8.050          0.020          1
HA             1              SER            HA             H
4.658          0.020          1
```

| | | | | |
|---|---|---|---|---|
| HB2 | 1 | SER | HB2 | H |
| 4.128 | 0.020 | 1 | | |
| HB3 | 1 | SER | HB3 | H |
| 4.128 | 0.020 | 1 | | |
| N | 1 | SER | N | N |
| 116.264 | 0.3 | 1 | | |
| C | 2 | ASN | C | C |
| 173.474 | 0.3 | 1 | | |
| CA | 2 | ASN | CA | C |
| 53.594 | 0.3 | 1 | | |
| CB | 2 | ASN | CB | C |
| 26.344 | 0.3 | 1 | | |
| H | 2 | ASN | H | H |
| 8.316 | 0.020 | 1 | | |
| HA | 2 | ASN | HA | H |
| 4.660 | 0.020 | 1 | | |
| HB2 | 2 | ASN | HB2 | H |
| 2.282 | 0.020 | 1 | | |
| HB3 | 2 | ASN | HB3 | H |
| 2.282 | 0.020 | 2 | | |
| N | 2 | ASN | N | N |
| 122.883 | 0.3 | 1 | | |
| C | 3 | ALA | C | C |
| 174.311 | 0.3 | 1 | | |
| CA | 3 | ALA | CA | C |
| 60.321 | 0.3 | 1 | | |
| CB | 3 | ALA | CB | C |
| 29.395 | 0.3 | 1 | | |
| H | 3 | ALA | H | H |
| 8.289 | 0.020 | 1 | | |
| HA | 3 | ALA | HA | H |
| 4.188 | 0.020 | 1 | | |
| HB | 3 | ALA | HB | H |
| 1.548 | 0.020 | 1 | | |
| N | 3 | ALA | N | N |
| 122.526 | 0.3 | 1 | | |
| C | 4 | GLN | C | C |
| 172.965 | 0.3 | 1 | | |
| CA | 4 | GLN | CA | C |
| 51.009 | 0.3 | 1 | | |
| CB | 4 | GLN | CB | C |
| 38.197 | 0.3 | 1 | | |
| H | 4 | GLN | H | H |
| 8.212 | 0.020 | 1 | | |
| HA | 4 | GLN | HA | H |
| 4.700 | 0.020 | 1 | | |
| HB2 | 4 | GLN | HB2 | H |
| 1.669 | 0.020 | 1 | | |
| HB3 | 4 | GLN | HB3 | H |
| 1.577 | 0.020 | 2 | | |
| HG2 | 4 | GLN | HG2 | H |
| 2.586 | 0.020 | 1 | | |
| HG3 | 4 | GLN | HG3 | H |
| 2.817 | 0.020 | 2 | | |
| N | 4 | GLN | N | N |
| 119.927 | 0.3 | 1 | | |
| C | 5 | GLN | C | C |
| 173.494 | 0.3 | 1 | | |

| | | | | |
|---|---|---|---|---|
| CA | 5 | GLN | CA | C |
| 53.100 | 0.3 | 1 | | |
| CB | 5 | GLN | CB | C |
| 26.290 | 0.3 | 1 | | |
| H | 5 | GLN | H | H |
| 8.214 | 0.020 | 1 | | |
| HA | 5 | GLN | HA | H |
| 4.687 | 0.020 | 1 | | |
| HB2 | 5 | GLN | HB2 | H |
| 1.585 | 0.020 | 1 | | |
| HB3 | 5 | GLN | HB3 | H |
| 1.585 | 0.020 | 2 | | |
| HG2 | 5 | GLN | HG2 | H |
| 2.551 | 0.020 | 1 | | |
| HG3 | 5 | GLN | HG3 | H |
| 2.831 | 0.020 | 2 | | |
| N | 5 | GLN | N | N |
| 120.433 | 0.3 | 1 | | |
| C | 6 | SER | C | C |
| 171.706 | 0.3 | 1 | | |
| CA | 6 | SER | CA | C |
| 56.270 | 0.3 | 1 | | |
| CB | 6 | SER | CB | C |
| 60.665 | 0.3 | 1 | | |
| H | 6 | SER | H | H |
| 8.241 | 0.020 | 1 | | |
| HA | 6 | SER | HA | H |
| 4.690 | 0.020 | 1 | | |
| HB2 | 6 | SER | HB2 | H |
| 3.730 | 0.020 | 1 | | |
| HB3 | 6 | SER | HB3 | H |
| 4.196 | 0.020 | 2 | | |
| N | 6 | SER | N | N |
| 116.723 | 0.3 | 1 | | |
| C | 7 | GLU | C | C |
| 173.718 | 0.3 | 1 | | |
| CA | 7 | GLU | CA | C |
| 53.829 | 0.3 | 1 | | |
| CB | 7 | GLU | CB | C |
| 27.232 | 0.3 | 1 | | |
| H | 7 | GLU | H | H |
| 8.197 | 0.020 | 1 | | |
| HA | 7 | GLU | HA | H |
| 4.676 | 0.020 | 1 | | |
| HB2 | 7 | GLU | HB2 | H |
| 1.905 | 0.020 | 1 | | |
| HB3 | 7 | GLU | HB3 | H |
| 1.905 | 0.020 | 1 | | |
| HG2 | 7 | GLU | HG2 | H |
| 2.083 | 0.020 | 1 | | |
| HG3 | 7 | GLU | HG3 | H |
| 2.083 | 0.020 | 1 | | |
| N | 7 | GLU | N | N |
| 121.958 | 0.3 | 1 | | |
| CA | 8 | CYS | CA | C |
| 50.998 | 0.3 | 1 | | |
| CB | 8 | CYS | CB | C |
| 38.589 | 0.3 | 1 | | |

| | | | | |
|---|---|---|---|---|
| H | 8 | CYS | H | H |
| 8.291 | 0.020 | 1 | | |
| HA | 8 | CYS | HA | H |
| 4.550 | 0.020 | 1 | | |
| HB2 | 8 | CYS | HB2 | H |
| 2.578 | 0.020 | 1 | | |
| HB3 | 8 | CYS | HB3 | H |
| 2.578 | 0.020 | 1 | | |
| N | 8 | CYS | N | N |
| 120.809 | 0.3 | 1 | | |
| C | 9 | ASN | C | C |
| 170.017 | 0.3 | 1 | | |
| CA | 9 | ASN | CA | C |
| 47.594 | 0.3 | 1 | | |
| CB | 9 | ASN | CB | C |
| 35.659 | 0.3 | 1 | | |
| H | 9 | ASN | H | H |
| 8.703 | 0.020 | 1 | | |
| HA | 9 | ASN | HA | H |
| 4.679 | 0.020 | 1 | | |
| HB2 | 9 | ASN | HB2 | H |
| 2.704 | 0.020 | 2 | | |
| HB3 | 9 | ASN | HB3 | H |
| 2.991 | 0.020 | 2 | | |
| N | 9 | ASN | N | N |
| 120.150 | 0.3 | 1 | | |
| C | 11 | VAL | C | C |
| 177.012 | 0.3 | 1 | | |
| CA | 11 | VAL | CA | C |
| 63.640 | 0.3 | 1 | | |
| CB | 11 | VAL | CB | C |
| 28.907 | 0.3 | 1 | | |
| H | 11 | VAL | H | H |
| 7.731 | 0.020 | 1 | | |
| HA | 11 | VAL | HA | H |
| 4.516 | 0.020 | 1 | | |
| HB | 11 | VAL | HB | H |
| 3.104 | 0.020 | 1 | | |
| HG1 | 11 | VAL | HG1 | H |
| 1.968 | 0.020 | 1 | | |
| HG2 | 11 | VAL | HG2 | H |
| 1.968 | 0.020 | 1 | | |
| N | 11 | VAL | N | N |
| 117.949 | 0.3 | 1 | | |
| C | 12 | GLY | C | C |
| 173.359 | 0.3 | 1 | | |
| CA | 12 | GLY | CA | C |
| 44.424 | 0.3 | 1 | | |
| H | 12 | GLY | H | H |
| 7.285 | 0.020 | 1 | | |
| HA2 | 12 | GLY | HA2 | H |
| 3.763 | 0.020 | 1 | | |
| HA3 | 12 | GLY | HA3 | H |
| 3.763 | 0.020 | 1 | | |
| N | 12 | GLY | N | N |
| 108.233 | 0.3 | 1 | | |
| C | 13 | ALA | C | C |
| 178.136 | 0.3 | 1 | | |

| | | | | |
|---|---|---|---|---|
| CA | 13 | ALA | CA | C |
| 52.023 | 0.3 | 1 | | |
| CB | 13 | ALA | CB | C |
| 15.946 | 0.3 | 1 | | |
| H | 13 | ALA | H | H |
| 8.273 | 0.020 | 1 | | |
| HA | 13 | ALA | HA | H |
| 4.023 | 0.020 | 1 | | |
| HB | 13 | ALA | HB | H |
| 1.339 | 0.020 | 1 | | |
| N | 13 | ALA | N | N |
| 124.528 | 0.3 | 1 | | |
| C | 14 | LEU | C | C |
| 178.136 | 0.3 | 1 | | |
| CA | 14 | LEU | CA | C |
| 51.916 | 0.3 | 1 | | |
| CB | 14 | LEU | CB | C |
| 15.823 | 0.3 | 1 | | |
| H | 14 | LEU | H | H |
| 8.436 | 0.020 | 1 | | |
| HA | 14 | LEU | HA | H |
| 4.016 | 0.020 | 1 | | |
| HB2 | 14 | LEU | HB2 | H |
| 2.000 | 0.020 | 1 | | |
| HB3 | 14 | LEU | HB3 | H |
| 2.000 | 0.020 | 1 | | |
| N | 14 | LEU | N | N |
| 119.386 | 0.3 | 1 | | |
| C | 15 | GLN | C | C |
| 174.032 | 0.3 | 1 | | |
| CA | 15 | GLN | CA | C |
| 57.538 | 0.3 | 1 | | |
| CB | 15 | GLN | CB | C |
| 25.246 | 0.3 | 1 | | |
| H | 15 | GLN | H | H |
| 7.540 | 0.020 | 1 | | |
| HA | 15 | GLN | HA | H |
| 3.634 | 0.020 | 1 | | |
| HB2 | 15 | GLN | HB2 | H |
| 2.154 | 0.020 | 1 | | |
| HB3 | 15 | GLN | HB3 | H |
| 2.154 | 0.020 | 1 | | |
| HG2 | 15 | GLN | HG2 | H |
| 2.397 | 0.020 | 1 | | |
| HG3 | 15 | GLN | HG3 | H |
| 2.397 | 0.020 | 1 | | |
| N | 15 | GLN | N | N |
| 118.263 | 0.3 | 1 | | |
| C | 16 | GLU | C | C |
| 176.016 | 0.3 | 1 | | |
| CA | 16 | GLU | CA | C |
| 56.389 | 0.3 | 1 | | |
| CB | 16 | GLU | CB | C |
| 26.832 | 0.3 | 1 | | |
| H | 16 | GLU | H | H |
| 7.775 | 0.020 | 1 | | |
| HA | 16 | GLU | HA | H |
| 3.866 | 0.020 | 1 | | |

| | | | | |
|---|---|---|---|---|
| HB2 | 16 | GLU | HB2 | H |
| 1.967 | 0.020 | 1 | | |
| HB3 | 16 | GLU | HB3 | H |
| 1.967 | 0.020 | 1 | | |
| N | 16 | GLU | N | N |
| 115.882 | 0.3 | 1 | | |
| CA | 17 | LEU | CA | C |
| 56.442 | 0.3 | 1 | | |
| H | 17 | LEU | H | H |
| 7.622 | 0.020 | 1 | | |
| HA | 17 | LEU | HA | H |
| 3.946 | 0.020 | 1 | | |
| HB2 | 17 | LEU | HB2 | H |
| 1.726 | 0.020 | 1 | | |
| HB3 | 17 | LEU | HB3 | H |
| 1.726 | 0.020 | 1 | | |
| N | 17 | LEU | N | N |
| 120.591 | 0.3 | 1 | | |
| C | 18 | VAL | C | C |
| 175.377 | 0.3 | 1 | | |
| CA | 18 | VAL | CA | C |
| 63.745 | 0.3 | 1 | | |
| CB | 18 | VAL | CB | C |
| 28.297 | 0.3 | 1 | | |
| H | 18 | VAL | H | H |
| 8.318 | 0.020 | 1 | | |
| HA | 18 | VAL | HA | H |
| 3.397 | 0.020 | 1 | | |
| HB | 18 | VAL | HB | H |
| 2.322 | 0.020 | 1 | | |
| N | 18 | VAL | N | N |
| 115.936 | 0.3 | 1 | | |
| C | 19 | VAL | C | C |
| 177.933 | 0.3 | 1 | | |
| CA | 19 | VAL | CA | C |
| 63.494 | 0.3 | 1 | | |
| CB | 19 | VAL | CB | C |
| 28.541 | 0.3 | 1 | | |
| H | 19 | VAL | H | H |
| 7.863 | 0.020 | 1 | | |
| HA | 19 | VAL | HA | H |
| 3.762 | 0.020 | 1 | | |
| HB | 19 | VAL | HB | H |
| 2.055 | 0.020 | 1 | | |
| HG1 | 19 | VAL | HG1 | H |
| 0.911 | 0.020 | 2 | | |
| HG2 | 19 | VAL | HG2 | H |
| 0.911 | 0.020 | 1 | | |
| N | 19 | VAL | N | N |
| 119.974 | 0.3 | 1 | | |
| C | 20 | GLN | C | C |
| 175.579 | 0.3 | 1 | | |
| CA | 20 | GLN | CA | C |
| 55.967 | 0.3 | 1 | | |
| CB | 20 | GLN | CB | C |
| 25.238 | 0.3 | 1 | | |
| H | 20 | GLN | H | H |
| 7.766 | 0.020 | 1 | | |

| | | | | |
|---|---|---|---|---|
| HA | 20 | GLN | HA | H |
| 3.883 | 0.020 | 1 | | |
| HB2 | 20 | GLN | HB2 | H |
| 2.198 | 0.020 | 1 | | |
| HB3 | 20 | GLN | HB3 | H |
| 2.198 | 0.020 | 1 | | |
| HG2 | 20 | GLN | HG2 | H |
| 2.389 | 0.020 | 1 | | |
| HG3 | 20 | GLN | HG3 | H |
| 2.389 | 0.020 | 1 | | |
| N | 20 | GLN | N | N |
| 121.355 | 0.3 | 1 | | |
| C | 22 | GLY | C | C |
| 172.249 | 0.3 | 1 | | |
| CA | 22 | GLY | CA | C |
| 42.640 | 0.3 | 1 | | |
| H | 22 | GLY | H | H |
| 7.670 | 0.020 | 1 | | |
| HA2 | 22 | GLY | HA2 | H |
| 3.988 | 0.020 | 1 | | |
| HA3 | 22 | GLY | HA3 | H |
| 3.988 | 0.020 | 2 | | |
| N | 22 | GLY | N | N |
| 106.857 | 0.3 | 1 | | |
| C | 23 | TRP | C | C |
| 172.485 | 0.3 | 1 | | |
| CA | 23 | TRP | CA | C |
| 50.567 | 0.3 | 1 | | |
| CB | 23 | TRP | CB | C |
| 28.907 | 0.3 | 1 | | |
| H | 23 | TRP | H | H |
| 7.536 | 0.020 | 1 | | |
| HA | 23 | TRP | HA | H |
| 4.968 | 0.020 | 1 | | |
| HB2 | 23 | TRP | HB2 | H |
| 3.277 | 0.020 | 1 | | |
| HB3 | 23 | TRP | HB3 | H |
| 3.223 | 0.020 | 2 | | |
| HD1 | 23 | TRP | HD1 | H |
| 7.195 | 0.020 | 1 | | |
| HE1 | 23 | TRP | HE1 | H |
| 9.950 | 0.020 | 1 | | |
| HZ2 | 23 | TRP | HZ2 | H |
| 6.757 | 0.020 | 1 | | |
| N | 23 | TRP | N | N |
| 121.648 | 0.3 | 1 | | |
| C | 24 | ARG | C | C |
| 173.124 | 0.3 | 1 | | |
| CA | 24 | ARG | CA | C |
| 53.780 | 0.3 | 1 | | |
| CB | 24 | ARG | CB | C |
| 27.809 | 0.3 | 1 | | |
| H | 24 | ARG | H | H |
| 8.401 | 0.020 | 1 | | |
| HA | 24 | ARG | HA | H |
| 4.056 | 0.020 | 1 | | |
| HB2 | 24 | ARG | HB2 | H |
| 1.761 | 0.020 | 1 | | |

| | | | | |
|---|---|---|---|---|
| HB3 | 24 | ARG | HB3 | H |
| 1.761 | 0.020 | 1 | | |
| HE | 24 | ARG | HE | H |
| 3.174 | 0.020 | 1 | | |
| N | 24 | ARG | N | N |
| 119.407 | 0.3 | 1 | | |
| CA | 25 | LEU | CA | C |
| 53.728 | 0.3 | 1 | | |
| CB | 25 | LEU | CB | C |
| 27.813 | 0.3 | 1 | | |
| H | 25 | LEU | H | H |
| 7.786 | 0.020 | 1 | | |
| HA | 25 | LEU | HA | H |
| 4.222 | 0.020 | 1 | | |
| N | 25 | LEU | N | N |
| 118.119 | 0.3 | 1 | | |
| C | 27 | GLU | C | C |
| 172.956 | 0.3 | 1 | | |
| CA | 27 | GLU | CA | C |
| 51.604 | 0.3 | 1 | | |
| CB | 27 | GLU | CB | C |
| 29.883 | 0.3 | 1 | | |
| H | 27 | GLU | H | H |
| 8.353 | 0.020 | 1 | | |
| HA | 27 | GLU | HA | H |
| 4.674 | 0.020 | 1 | | |
| HB2 | 27 | GLU | HB2 | H |
| 1.936 | 0.020 | 1 | | |
| HB3 | 27 | GLU | HB3 | H |
| 2.181 | 0.020 | 2 | | |
| HG2 | 27 | GLU | HG2 | H |
| 3.006 | 0.020 | 1 | | |
| HG3 | 27 | GLU | HG3 | H |
| 3.006 | 0.020 | 2 | | |
| N | 27 | GLU | N | N |
| 122.417 | 0.3 | 1 | | |
| CA | 28 | TYR | CA | C |
| 51.602 | 0.3 | 1 | | |
| H | 28 | TYR | H | H |
| 9.065 | 0.020 | 1 | | |
| HA | 28 | TYR | HA | H |
| 5.594 | 0.020 | 1 | | |
| HB2 | 28 | TYR | HB2 | H |
| 2.720 | 0.020 | 1 | | |
| HB3 | 28 | TYR | HB3 | H |
| 2.720 | 0.020 | 2 | | |
| HD1 | 28 | TYR | HD1 | H |
| 6.900 | 0.020 | 1 | | |
| HE1 | 28 | TYR | HE1 | H |
| 6.545 | 0.020 | 1 | | |
| N | 28 | TYR | N | N |
| 126.987 | 0.3 | 1 | | |
| C | 29 | THR | C | C |
| 170.601 | 0.3 | 1 | | |
| CA | 29 | THR | CA | C |
| 58.160 | 0.3 | 1 | | |
| CB | 29 | THR | CB | C |
| 68.776 | 0.3 | 1 | | |

| | | | | |
|---|---|---|---|---|
| H | 29 | THR | H | H |
| 8.531 | 0.020 | 1 | | |
| HA | 29 | THR | HA | H |
| 4.540 | 0.020 | 1 | | |
| HB | 29 | THR | HB | H |
| 3.840 | 0.020 | 1 | | |
| HG2 | 29 | THR | HG2 | H |
| 1.073 | 0.020 | 1 | | |
| N | 29 | THR | N | N |
| 117.650 | 0.3 | 1 | | |
| C | 30 | VAL | C | C |
| 173.258 | 0.3 | 1 | | |
| CA | 30 | VAL | CA | C |
| 60.094 | 0.3 | 1 | | |
| CB | 30 | VAL | CB | C |
| 29.319 | 0.3 | 1 | | |
| H | 30 | VAL | H | H |
| 8.990 | 0.020 | 1 | | |
| HA | 30 | VAL | HA | H |
| 4.595 | 0.020 | 1 | | |
| HB | 30 | VAL | HB | H |
| 2.116 | 0.020 | 1 | | |
| HG1 | 30 | VAL | HG1 | H |
| 1.068 | 0.020 | 1 | | |
| HG2 | 30 | VAL | HG2 | H |
| 1.068 | 0.020 | 1 | | |
| N | 30 | VAL | N | N |
| 127.369 | 0.3 | 1 | | |
| C | 31 | THR | C | C |
| 172.047 | 0.3 | 1 | | |
| CA | 31 | THR | CA | C |
| 59.394 | 0.3 | 1 | | |
| CB | 31 | THR | CB | C |
| 66.291 | 0.3 | 1 | | |
| H | 31 | THR | H | H |
| 8.705 | 0.020 | 1 | | |
| HA | 31 | THR | HA | H |
| 4.349 | 0.020 | 1 | | |
| HB | 31 | THR | HB | H |
| 4.074 | 0.020 | 1 | | |
| HG1 | 31 | THR | HG1 | H |
| 1.087 | 0.020 | 1 | | |
| N | 31 | THR | N | N |
| 121.646 | 0.3 | 1 | | |
| C | 32 | GLN | C | C |
| 170.904 | 0.3 | 1 | | |
| CA | 32 | GLN | CA | C |
| 53.471 | 0.3 | 1 | | |
| CB | 32 | GLN | CB | C |
| 28.907 | 0.3 | 1 | | |
| H | 32 | GLN | H | H |
| 7.885 | 0.020 | 1 | | |
| HA | 32 | GLN | HA | H |
| 4.685 | 0.020 | 1 | | |
| HB2 | 32 | GLN | HB2 | H |
| 1.935 | 0.020 | 1 | | |
| HB3 | 32 | GLN | HB3 | H |
| 1.935 | 0.020 | 1 | | |

| | | | | |
|---|---|---|---|---|
| HG2 | 32 | GLN | HG2 | H |
| 2.201 | 0.020 | 1 | | |
| HG3 | 32 | GLN | HG3 | H |
| 2.201 | 0.020 | 1 | | |
| HE21 | 32 | GLN | HE21 | H |
| 7.261 | 0.020 | 1 | | |
| HE22 | 32 | GLN | HE22 | H |
| 6.888 | 0.020 | 1 | | |
| N | 32 | GLN | N | N |
| 121.190 | 0.3 | 1 | | |
| C | 33 | GLU | C | C |
| 172.417 | 0.3 | 1 | | |
| CA | 33 | GLU | CA | C |
| 52.370 | 0.3 | 1 | | |
| CB | 33 | GLU | CB | C |
| 28.769 | 0.3 | 1 | | |
| H | 33 | GLU | H | H |
| 8.316 | 0.020 | 1 | | |
| HA | 33 | GLU | HA | H |
| 4.685 | 0.020 | 1 | | |
| HB2 | 33 | GLU | HB2 | H |
| 1.781 | 0.020 | 1 | | |
| N | 33 | GLU | N | N |
| 124.085 | 0.3 | 1 | | |
| C | 34 | SER | C | C |
| 170.904 | 0.3 | 1 | | |
| CA | 34 | SER | CA | C |
| 54.725 | 0.3 | 1 | | |
| CB | 34 | SER | CB | C |
| 62.221 | 0.3 | 1 | | |
| H | 34 | SER | H | H |
| 8.197 | 0.020 | 1 | | |
| HA | 34 | SER | HA | H |
| 4.687 | 0.020 | 1 | | |
| HB2 | 34 | SER | HB2 | H |
| 3.635 | 0.020 | 1 | | |
| HB3 | 34 | SER | HB3 | H |
| 3.635 | 0.020 | 1 | | |
| HG | 34 | SER | HG | H |
| 2.603 | 0.020 | 1 | | |
| N | 34 | SER | N | N |
| 118.212 | 0.3 | 1 | | |
| C | 35 | GLY | C | C |
| 169.424 | 0.3 | 1 | | |
| CA | 35 | GLY | CA | C |
| 41.480 | 0.3 | 1 | | |
| H | 35 | GLY | H | H |
| 8.147 | 0.020 | 1 | | |
| HA2 | 35 | GLY | HA2 | H |
| 3.837 | 0.020 | 2 | | |
| HA3 | 35 | GLY | HA3 | H |
| 4.692 | 0.020 | 2 | | |
| N | 35 | GLY | N | N |
| 109.048 | 0.3 | 1 | | |
| H | 37 | ALA | H | H |
| 8.246 | 0.020 | 1 | | |
| HA | 37 | ALA | HA | H |
| 4.254 | 0.020 | 1 | | |

| | | | | |
|---|---|---|---|---|
| HB | 37 | ALA | HB | H |
| 1.673 | 0.020 | 1 | | |
| N | 37 | ALA | N | N |
| 120.862 | 0.3 | 1 | | |
| C | 39 | ARG | C | C |
| 171.442 | 0.3 | 1 | | |
| CA | 39 | ARG | CA | C |
| 52.850 | 0.3 | 1 | | |
| CB | 39 | ARG | CB | C |
| 27.564 | 0.3 | 1 | | |
| H | 39 | ARG | H | H |
| 7.719 | 0.020 | 1 | | |
| HA | 39 | ARG | HA | H |
| 4.660 | 0.020 | 1 | | |
| HB2 | 39 | ARG | HB2 | H |
| 1.568 | 0.020 | 1 | | |
| HB3 | 39 | ARG | HB3 | H |
| 1.568 | 0.020 | 1 | | |
| HG2 | 39 | ARG | HG2 | H |
| 1.353 | 0.020 | 1 | | |
| HG3 | 39 | ARG | HG3 | H |
| 1.353 | 0.020 | 1 | | |
| HD2 | 39 | ARG | HD2 | H |
| 2.093 | 0.020 | 1 | | |
| HD3 | 39 | ARG | HD3 | H |
| 2.093 | 0.020 | 1 | | |
| HE | 39 | ARG | HE | H |
| 3.053 | 0.020 | 1 | | |
| N | 39 | ARG | N | N |
| 123.947 | 0.3 | 1 | | |
| C | 41 | GLU | C | C |
| 172.384 | 0.3 | 1 | | |
| CA | 41 | GLU | CA | C |
| 52.842 | 0.3 | 1 | | |
| CB | 41 | GLU | CB | C |
| 29.395 | 0.3 | 1 | | |
| H | 41 | GLU | H | H |
| 8.128 | 0.020 | 1 | | |
| HA | 41 | GLU | HA | H |
| 4.367 | 0.020 | 1 | | |
| HB2 | 41 | GLU | HB2 | H |
| 1.943 | 0.020 | 1 | | |
| HB3 | 41 | GLU | HB3 | H |
| 1.943 | 0.020 | 1 | | |
| HG2 | 41 | GLU | HG2 | H |
| 1.642 | 0.020 | 1 | | |
| HG3 | 41 | GLU | HG3 | H |
| 1.642 | 0.020 | 1 | | |
| N | 41 | GLU | N | N |
| 119.493 | 0.3 | 1 | | |
| C | 42 | PHE | C | C |
| 172.384 | 0.3 | 1 | | |
| CA | 42 | PHE | CA | C |
| 52.842 | 0.3 | 1 | | |
| CB | 42 | PHE | CB | C |
| 29.395 | 0.3 | 1 | | |
| H | 42 | PHE | H | H |
| 8.775 | 0.020 | 1 | | |

| | | | | |
|---|---|---|---|---|
| HA | 42 | PHE | HA | H |
| 4.880 | 0.020 | 1 | | |
| HB2 | 42 | PHE | HB2 | H |
| 2.553 | 0.020 | 1 | | |
| HB3 | 42 | PHE | HB3 | H |
| 2.553 | 0.020 | 1 | | |
| HD1 | 42 | PHE | HD1 | H |
| 6.881 | 0.020 | 1 | | |
| HE1 | 42 | PHE | HE1 | H |
| 7.236 | 0.020 | 1 | | |
| N | 42 | PHE | N | N |
| 121.600 | 0.3 | 1 | | |
| C | 43 | THR | C | C |
| 170.803 | 0.3 | 1 | | |
| CA | 43 | THR | CA | C |
| 58.772 | 0.3 | 1 | | |
| CB | 43 | THR | CB | C |
| 67.826 | 0.3 | 1 | | |
| H | 43 | THR | H | H |
| 8.717 | 0.020 | 1 | | |
| HA | 43 | THR | HA | H |
| 5.105 | 0.020 | 1 | | |
| HB | 43 | THR | HB | H |
| 3.836 | 0.020 | 1 | | |
| HG2 | 43 | THR | HG2 | H |
| 1.068 | 0.020 | 1 | | |
| N | 43 | THR | N | N |
| 116.346 | 0.3 | 1 | | |
| CA | 44 | MET | CA | C |
| 58.782 | 0.3 | 1 | | |
| H | 44 | MET | H | H |
| 9.360 | 0.020 | 1 | | |
| HA | 44 | MET | HA | H |
| 5.392 | 0.020 | 1 | | |
| HB2 | 44 | MET | HB2 | H |
| 1.896 | 0.020 | 1 | | |
| HB3 | 44 | MET | HB3 | H |
| 1.896 | 0.020 | 2 | | |
| HG2 | 44 | MET | HG2 | H |
| 2.327 | 0.020 | 1 | | |
| HG3 | 44 | MET | HG3 | H |
| 2.327 | 0.020 | 1 | | |
| N | 44 | MET | N | N |
| 125.128 | 0.3 | 1 | | |
| C | 45 | THR | C | C |
| 170.904 | 0.3 | 1 | | |
| CA | 45 | THR | CA | C |
| 58.162 | 0.3 | 1 | | |
| CB | 45 | THR | CB | C |
| 67.634 | 0.3 | 1 | | |
| H | 45 | THR | H | H |
| 9.190 | 0.020 | 1 | | |
| HA | 45 | THR | HA | H |
| 4.937 | 0.020 | 1 | | |
| HB | 45 | THR | HB | H |
| 3.910 | 0.020 | 1 | | |
| HG1 | 45 | THR | HG1 | H |
| 1.032 | 0.020 | 1 | | |

| | | | | |
|---|---|---|---|---|
| N | 45 | THR | N | N |
| 116.395 | 0.3 | 1 | | |
| C | 46 | CYS | C | C |
| 169.350 | 0.3 | 1 | | |
| CA | 46 | CYS | CA | C |
| 54.079 | 0.3 | 1 | | |
| CB | 46 | CYS | CB | C |
| 26.896 | 0.3 | 1 | | |
| H | 46 | CYS | H | H |
| 9.124 | 0.020 | 1 | | |
| HA | 46 | CYS | HA | H |
| 4.566 | 0.020 | 1 | | |
| HB2 | 46 | CYS | HB2 | H |
| 2.260 | 0.020 | 1 | | |
| HB3 | 46 | CYS | HB3 | H |
| 2.260 | 0.020 | 1 | | |
| N | 46 | CYS | N | N |
| 124.254 | 0.3 | 1 | | |
| C | 47 | ARG | C | C |
| 173.314 | 0.3 | 1 | | |
| CA | 47 | ARG | CA | C |
| 51.588 | 0.3 | 1 | | |
| CB | 47 | ARG | CB | C |
| 30.648 | 0.3 | 1 | | |
| H | 47 | ARG | H | H |
| 8.671 | 0.020 | 1 | | |
| HA | 47 | ARG | HA | H |
| 5.201 | 0.020 | 1 | | |
| HB2 | 47 | ARG | HB2 | H |
| 1.416 | 0.020 | 1 | | |
| HB3 | 47 | ARG | HB3 | H |
| 1.416 | 0.020 | 1 | | |
| HG2 | 47 | ARG | HG2 | H |
| 1.232 | 0.020 | 1 | | |
| HG3 | 47 | ARG | HG3 | H |
| 1.232 | 0.020 | 1 | | |
| HD2 | 47 | ARG | HD2 | H |
| 1.658 | 0.020 | 1 | | |
| HD3 | 47 | ARG | HD3 | H |
| 1.658 | 0.020 | 1 | | |
| N | 47 | ARG | N | N |
| 128.627 | 0.3 | 1 | | |
| C | 48 | VAL | C | C |
| 170.805 | 0.3 | 1 | | |
| CA | 48 | VAL | CA | C |
| 58.397 | 0.3 | 1 | | |
| CB | 48 | VAL | CB | C |
| 32.934 | 0.3 | 1 | | |
| H | 48 | VAL | H | H |
| 8.096 | 0.020 | 1 | | |
| HA | 48 | VAL | HA | H |
| 4.080 | 0.020 | 1 | | |
| HB | 48 | VAL | HB | H |
| 2.224 | 0.020 | 1 | | |
| HG1 | 48 | VAL | HG1 | H |
| 0.975 | 0.020 | 1 | | |
| HG2 | 48 | VAL | HG2 | H |
| 0.975 | 0.020 | 1 | | |

| | | | | |
|---|---|---|---|---|
| N | 48 | VAL | N | N |
| 126.933 | 0.3 | 1 | | |
| C | 49 | GLU | C | C |
| 173.140 | 0.3 | 1 | | |
| CA | 49 | GLU | CA | C |
| 55.357 | 0.3 | 1 | | |
| CB | 49 | GLU | CB | C |
| 23.782 | 0.3 | 1 | | |
| H | 49 | GLU | H | H |
| 8.859 | 0.020 | 1 | | |
| HA | 49 | GLU | HA | H |
| 3.259 | 0.020 | 1 | | |
| HB2 | 49 | GLU | HB2 | H |
| 1.237 | 0.020 | 1 | | |
| HB3 | 49 | GLU | HB3 | H |
| 1.237 | 0.020 | 1 | | |
| HG2 | 49 | GLU | HG2 | H |
| 1.493 | 0.020 | 1 | | |
| HG3 | 49 | GLU | HG3 | H |
| 1.493 | 0.020 | 1 | | |
| N | 49 | GLU | N | N |
| 123.705 | 0.3 | 1 | | |
| C | 50 | ARG | C | C |
| 173.133 | 0.3 | 1 | | |
| CA | 50 | ARG | CA | C |
| 53.146 | 0.3 | 1 | | |
| CB | 50 | ARG | CB | C |
| 26.588 | 0.3 | 1 | | |
| H | 50 | ARG | H | H |
| 7.312 | 0.020 | 1 | | |
| HA | 50 | ARG | HA | H |
| 4.135 | 0.020 | 1 | | |
| HB2 | 50 | ARG | HB2 | H |
| 1.300 | 0.020 | 1 | | |
| HG2 | 50 | ARG | HG2 | H |
| 1.011 | 0.020 | 1 | | |
| N | 50 | ARG | N | N |
| 120.474 | 0.3 | 1 | | |
| C | 51 | PHE | C | C |
| 173.133 | 0.3 | 1 | | |
| CA | 51 | PHE | CA | C |
| 53.146 | 0.3 | 1 | | |
| H | 51 | PHE | H | H |
| 8.628 | 0.020 | 1 | | |
| HA | 51 | PHE | HA | H |
| 4.660 | 0.020 | 1 | | |
| HB2 | 51 | PHE | HB2 | H |
| 2.811 | 0.020 | 1 | | |
| HB3 | 51 | PHE | HB3 | H |
| 3.153 | 0.020 | 2 | | |
| HD1 | 51 | PHE | HD1 | H |
| 7.068 | 0.020 | 1 | | |
| N | 51 | PHE | N | N |
| 122.221 | 0.3 | 1 | | |
| CA | 52 | ILE | CA | C |
| 53.634 | 0.3 | 1 | | |
| H | 52 | ILE | H | H |
| 8.452 | 0.020 | 1 | | |

| | | | | |
|---|---|---|---|---|
| HA | 52 | ILE | HA | H |
| 4.943 | 0.020 | 1 | | |
| HB | 52 | ILE | HB | H |
| 0.701 | 0.020 | 1 | | |
| HG12 | 52 | ILE | HG12 | H |
| 1.545 | 0.020 | 1 | | |
| HG13 | 52 | ILE | HG13 | H |
| 1.545 | 0.020 | 1 | | |
| N | 52 | ILE | N | N |
| 121.877 | 0.3 | 1 | | |
| C | 53 | GLU | C | C |
| 171.777 | 0.3 | 1 | | |
| CA | 53 | GLU | CA | C |
| 51.238 | 0.3 | 1 | | |
| CB | 53 | GLU | CB | C |
| 32.080 | 0.3 | 1 | | |
| H | 53 | GLU | H | H |
| 8.561 | 0.020 | 1 | | |
| HA | 53 | GLU | HA | H |
| 4.697 | 0.020 | 1 | | |
| HB2 | 53 | GLU | HB2 | H |
| 1.365 | 0.020 | 1 | | |
| HB3 | 53 | GLU | HB3 | H |
| 1.365 | 0.020 | 1 | | |
| HG2 | 53 | GLU | HG2 | H |
| 2.132 | 0.020 | 1 | | |
| HG3 | 53 | GLU | HG3 | H |
| 2.132 | 0.020 | 1 | | |
| N | 53 | GLU | N | N |
| 124.418 | 0.3 | 1 | | |
| CB | 54 | ILE | CB | C |
| 32.080 | 0.3 | 1 | | |
| H | 54 | ILE | H | H |
| 8.845 | 0.020 | 1 | | |
| HA | 54 | ILE | HA | H |
| 4.898 | 0.020 | 1 | | |
| HB | 54 | ILE | HB | H |
| 0.738 | 0.020 | 1 | | |
| HG2 | 54 | ILE | HG2 | H |
| 1.545 | 0.020 | 1 | | |
| HD1 | 54 | ILE | HD1 | H |
| 0.940 | 0.020 | 1 | | |
| N | 54 | ILE | N | N |
| 120.479 | 0.3 | 1 | | |
| C | 55 | GLY | C | C |
| 169.020 | 0.3 | 1 | | |
| CA | 55 | GLY | CA | C |
| 40.943 | 0.3 | 1 | | |
| H | 55 | GLY | H | H |
| 9.177 | 0.020 | 1 | | |
| HA2 | 55 | GLY | HA2 | H |
| 3.672 | 0.020 | 1 | | |
| HA3 | 55 | GLY | HA3 | H |
| 4.678 | 0.020 | 2 | | |
| N | 55 | GLY | N | N |
| 112.301 | 0.3 | 1 | | |
| C | 56 | SER | C | C |
| 171.341 | 0.3 | 1 | | |

| | | | | |
|---|---|---|---|---|
| CA | 56 | SER | CA | C |
| 53.780 | 0.3 | 1 | | |
| CB | 56 | SER | CB | C |
| 63.810 | 0.3 | 1 | | |
| H | 56 | SER | H | H |
| 8.625 | 0.020 | 1 | | |
| HA | 56 | SER | HA | H |
| 5.910 | 0.020 | 1 | | |
| HG | 56 | SER | HG | H |
| 3.727 | 0.020 | 1 | | |
| N | 56 | SER | N | N |
| 114.811 | 0.3 | 1 | | |
| C | 57 | GLY | C | C |
| 169.928 | 0.3 | 1 | | |
| CA | 57 | GLY | CA | C |
| 42.860 | 0.3 | 1 | | |
| H | 57 | GLY | H | H |
| 8.552 | 0.020 | 1 | | |
| HA2 | 57 | GLY | HA2 | H |
| 3.948 | 0.020 | 1 | | |
| HA3 | 57 | GLY | HA3 | H |
| 4.440 | 0.020 | 2 | | |
| N | 57 | GLY | N | N |
| 106.946 | 0.3 | 1 | | |
| C | 58 | THR | C | C |
| 170.467 | 0.3 | 1 | | |
| CA | 58 | THR | CA | C |
| 59.902 | 0.3 | 1 | | |
| CB | 58 | THR | CB | C |
| 66.004 | 0.3 | 1 | | |
| H | 58 | THR | H | H |
| 8.273 | 0.020 | 1 | | |
| HA | 58 | THR | HA | H |
| 4.638 | 0.020 | 1 | | |
| N | 58 | THR | N | N |
| 109.376 | 0.3 | 1 | | |
| CA | 59 | SER | CA | C |
| 39.410 | 0.3 | 1 | | |
| CB | 59 | SER | CB | C |
| 22.842 | 0.3 | 1 | | |
| H | 59 | SER | H | H |
| 7.322 | 0.020 | 1 | | |
| HA | 59 | SER | HA | H |
| 4.699 | 0.020 | 1 | | |
| HB2 | 59 | SER | HB2 | H |
| 2.920 | 0.020 | 1 | | |
| HB3 | 59 | SER | HB3 | H |
| 2.920 | 0.020 | 1 | | |
| N | 59 | SER | N | N |
| 112.329 | 0.3 | 1 | | |
| C | 60 | LYS | C | C |
| 172.956 | 0.3 | 1 | | |
| CB | 60 | LYS | CB | C |
| 28.348 | 0.3 | 1 | | |
| H | 60 | LYS | H | H |
| 8.408 | 0.020 | 1 | | |
| HA | 60 | LYS | HA | H |
| 4.695 | 0.020 | 1 | | |

| | | | | |
|---|---|---|---|---|
| HB2 | 60 | LYS | HB2 | H |
| 1.948 | 0.020 | 1 | | |
| HB3 | 60 | LYS | HB3 | H |
| 1.948 | 0.020 | 1 | | |
| N | 60 | LYS | N | N |
| 123.979 | 0.3 | 1 | | |
| C | 62 | LEU | C | C |
| 172.014 | 0.3 | 1 | | |
| CA | 62 | LEU | CA | C |
| 52.846 | 0.3 | 1 | | |
| CB | 62 | LEU | CB | C |
| 29.395 | 0.3 | 1 | | |
| H | 62 | LEU | H | H |
| 7.693 | 0.020 | 1 | | |
| HA | 62 | LEU | HA | H |
| 4.331 | 0.020 | 1 | | |
| HB2 | 62 | LEU | HB2 | H |
| 1.985 | 0.020 | 1 | | |
| HB3 | 62 | LEU | HB3 | H |
| 1.985 | 0.020 | 1 | | |
| HG | 62 | LEU | HG | H |
| 1.728 | 0.020 | 1 | | |
| N | 62 | LEU | N | N |
| 119.625 | 0.3 | 1 | | |
| C | 63 | ALA | C | C |
| 175.882 | 0.3 | 1 | | |
| CA | 63 | ALA | CA | C |
| 52.846 | 0.3 | 1 | | |
| CB | 63 | ALA | CB | C |
| 15.348 | 0.3 | 1 | | |
| H | 63 | ALA | H | H |
| 8.034 | 0.020 | 1 | | |
| HA | 63 | ALA | HA | H |
| 3.636 | 0.020 | 1 | | |
| HB | 63 | ALA | HB | H |
| 1.233 | 0.020 | 1 | | |
| N | 63 | ALA | N | N |
| 122.781 | 0.3 | 1 | | |
| C | 65 | ARG | C | C |
| 175.007 | 0.3 | 1 | | |
| CA | 65 | ARG | CA | C |
| 57.215 | 0.3 | 1 | | |
| CB | 65 | ARG | CB | C |
| 27.516 | 0.3 | 1 | | |
| H | 65 | ARG | H | H |
| 7.639 | 0.020 | 1 | | |
| HA | 65 | ARG | HA | H |
| 3.708 | 0.020 | 1 | | |
| HB2 | 65 | ARG | HB2 | H |
| 1.874 | 0.020 | 1 | | |
| HB3 | 65 | ARG | HB3 | H |
| 1.874 | 0.020 | 1 | | |
| N | 65 | ARG | N | N |
| 117.798 | 0.3 | 1 | | |
| C | 66 | ASN | C | C |
| 175.007 | 0.3 | 1 | | |
| CA | 66 | ASN | CA | C |
| 53.435 | 0.3 | 1 | | |

| | | | | |
|---|---|---|---|---|
| CB | 66 | ASN | CB | C |
| 35.946 | 0.3 | 1 | | |
| H | 66 | ASN | H | H |
| 8.132 | 0.020 | 1 | | |
| HA | 66 | ASN | HA | H |
| 4.532 | 0.020 | 1 | | |
| HB2 | 66 | ASN | HB2 | H |
| 2.880 | 0.020 | 1 | | |
| HB3 | 66 | ASN | HB3 | H |
| 2.880 | 0.020 | 1 | | |
| HD21 | 66 | ASN | HD21 | H |
| 7.094 | 0.020 | 1 | | |
| N | 66 | ASN | N | N |
| 117.005 | 0.3 | 1 | | |
| C | 67 | ALA | C | C |
| 176.184 | 0.3 | 1 | | |
| CA | 67 | ALA | CA | C |
| 52.533 | 0.3 | 1 | | |
| CB | 67 | ALA | CB | C |
| 15.362 | 0.3 | 1 | | |
| H | 67 | ALA | H | H |
| 8.345 | 0.020 | 1 | | |
| HA | 67 | ALA | HA | H |
| 4.660 | 0.020 | 1 | | |
| HB | 67 | ALA | HB | H |
| 1.435 | 0.020 | 1 | | |
| N | 67 | ALA | N | N |
| 123.376 | 0.3 | 1 | | |
| C | 68 | ALA | C | C |
| 176.891 | 0.3 | 1 | | |
| CA | 68 | ALA | CA | C |
| 52.223 | 0.3 | 1 | | |
| CB | 68 | ALA | CB | C |
| 15.664 | 0.3 | 1 | | |
| H | 68 | ALA | H | H |
| 8.568 | 0.020 | 1 | | |
| HA | 68 | ALA | HA | H |
| 4.058 | 0.020 | 1 | | |
| HB | 68 | ALA | HB | H |
| 1.527 | 0.020 | 1 | | |
| N | 68 | ALA | N | N |
| 119.272 | 0.3 | 1 | | |
| C | 69 | ALA | C | C |
| 178.438 | 0.3 | 1 | | |
| CA | 69 | ALA | CA | C |
| 52.524 | 0.3 | 1 | | |
| CB | 69 | ALA | CB | C |
| 15.365 | 0.3 | 1 | | |
| H | 69 | ALA | H | H |
| 8.514 | 0.020 | 1 | | |
| HA | 69 | ALA | HA | H |
| 3.966 | 0.020 | 1 | | |
| HB | 69 | ALA | HB | H |
| 1.527 | 0.020 | 1 | | |
| N | 69 | ALA | N | N |
| 119.440 | 0.3 | 1 | | |
| C | 71 | MET | C | C |
| 175.310 | 0.3 | 1 | | |

| | | | | |
|---|---|---|---|---|
| CA | 71 | MET | CA | C |
| 51.659 | 0.3 | 1 | | |
| CB | 71 | MET | CB | C |
| 15.670 | 0.3 | 1 | | |
| H | 71 | MET | H | H |
| 8.387 | 0.020 | 1 | | |
| HA | 71 | MET | HA | H |
| 4.660 | 0.020 | 1 | | |
| HB2 | 71 | MET | HB2 | H |
| 1.292 | 0.020 | 1 | | |
| HB3 | 71 | MET | HB3 | H |
| 1.292 | 0.020 | 1 | | |
| N | 71 | MET | N | N |
| 120.698 | 0.3 | 1 | | |
| CB | 72 | LEU | CB | C |
| 29.395 | 0.3 | 1 | | |
| H | 72 | LEU | H | H |
| 8.751 | 0.020 | 1 | | |
| HA | 72 | LEU | HA | H |
| 3.709 | 0.020 | 1 | | |
| N | 72 | LEU | N | N |
| 121.565 | 0.3 | 1 | | |
| H | 73 | LEU | H | H |
| 6.998 | 0.020 | 1 | | |
| HA | 73 | LEU | HA | H |
| 4.001 | 0.020 | 1 | | |
| HB2 | 73 | LEU | HB2 | H |
| 1.703 | 0.020 | 1 | | |
| HB3 | 73 | LEU | HB3 | H |
| 1.703 | 0.020 | 1 | | |
| HG | 73 | LEU | HG | H |
| 1.543 | 0.020 | 1 | | |
| N | 73 | LEU | N | N |
| 116.181 | 0.3 | 1 | | |
| C | 74 | ARG | C | C |
| 176.004 | 0.3 | 1 | | |
| CA | 74 | ARG | CA | C |
| 54.952 | 0.3 | 1 | | |
| CB | 74 | ARG | CB | C |
| 26.605 | 0.3 | 1 | | |
| H | 74 | ARG | H | H |
| 7.504 | 0.020 | 1 | | |
| HA | 74 | ARG | HA | H |
| 3.865 | 0.020 | 1 | | |
| HB2 | 74 | ARG | HB2 | H |
| 1.864 | 0.020 | 1 | | |
| HB3 | 74 | ARG | HB3 | H |
| 1.864 | 0.020 | 1 | | |
| N | 74 | ARG | N | N |
| 119.653 | 0.3 | 1 | | |
| C | 75 | VAL | C | C |
| 174.368 | 0.3 | 1 | | |
| CA | 75 | VAL | CA | C |
| 61.297 | 0.3 | 1 | | |
| CB | 75 | VAL | CB | C |
| 28.785 | 0.3 | 1 | | |
| H | 75 | VAL | H | H |
| 8.172 | 0.020 | 1 | | |

| | | | | |
|---|---|---|---|---|
| HA | 75 | VAL | HA | H |
| 3.764 | 0.020 | 1 | | |
| HB | 75 | VAL | HB | H |
| 2.058 | 0.020 | 1 | | |
| HG1 | 75 | VAL | HG1 | H |
| 0.830 | 0.020 | 1 | | |
| HG2 | 75 | VAL | HG2 | H |
| 0.830 | 0.020 | 1 | | |
| N | 75 | VAL | N | N |
| 113.698 | 0.3 | 1 | | |
| C | 77 | THR | C | C |
| 171.543 | 0.3 | 1 | | |
| CA | 77 | THR | CA | C |
| 59.712 | 0.3 | 1 | | |
| CB | 77 | THR | CB | C |
| 67.226 | 0.3 | 1 | | |
| H | 77 | THR | H | H |
| 7.614 | 0.020 | 1 | | |
| HA | 77 | THR | HA | H |
| 4.667 | 0.020 | 1 | | |
| HB | 77 | THR | HB | H |
| 4.219 | 0.020 | 1 | | |
| HG2 | 77 | THR | HG2 | H |
| 1.160 | 0.020 | 1 | | |
| N | 77 | THR | N | N |
| 112.134 | 0.3 | 1 | | |
| C | 78 | VAL | C | C |
| 175.982 | 0.3 | 1 | | |
| CA | 78 | VAL | CA | C |
| 57.525 | 0.3 | 1 | | |
| CB | 78 | VAL | CB | C |
| 29.720 | 0.3 | 1 | | |
| H | 78 | VAL | H | H |
| 7.783 | 0.020 | 1 | | |
| HA | 78 | VAL | HA | H |
| 4.312 | 0.020 | 1 | | |
| HB | 78 | VAL | HB | H |
| 2.079 | 0.020 | 1 | | |
| HG1 | 78 | VAL | HG1 | H |
| 0.976 | 0.020 | 1 | | |
| HG2 | 78 | VAL | HG2 | H |
| 0.976 | 0.020 | 1 | | |
| N | 78 | VAL | N | N |
| 124.088 | 0.3 | 1 | | |
| C | 80 | LEU | C | C |
| 178.080 | 0.3 | 1 | | |
| CA | 80 | LEU | CA | C |
| 52.947 | 0.3 | 1 | | |
| CB | 80 | LEU | CB | C |
| 39.401 | 0.3 | 1 | | |
| H | 80 | LEU | H | H |
| 7.851 | 0.020 | 1 | | |
| HA | 80 | LEU | HA | H |
| 4.267 | 0.020 | 1 | | |
| HB2 | 80 | LEU | HB2 | H |
| 2.538 | 0.020 | 1 | | |
| HB3 | 80 | LEU | HB3 | H |
| 2.538 | 0.020 | 1 | | |

| | | | | |
|---|---|---|---|---|
| N | 80 | LEU | N | N |
| 126.412 | 0.3 | 1 | | |
| H | 81 | ASP | H | H |
| 7.999 | 0.020 | 1 | | |
| HA | 81 | ASP | HA | H |
| 4.643 | 0.020 | 1 | | |
| HB2 | 81 | ASP | HB2 | H |
| 2.895 | 0.020 | 1 | | |
| HB3 | 81 | ASP | HB3 | H |
| 2.895 | 0.020 | 1 | | |
| N | 81 | ASP | N | N |
| 120.618 | 0.3 | 1 | | |
| C | 82 | ALA | C | C |
| 174.606 | 0.3 | 1 | | |
| CA | 82 | ALA | CA | C |
| 49.694 | 0.3 | 1 | | |
| CB | 82 | ALA | CB | C |
| 16.460 | 0.3 | 1 | | |
| H | 82 | ALA | H | H |
| 7.960 | 0.020 | 1 | | |
| HA | 82 | ALA | HA | H |
| 4.193 | 0.020 | 1 | | |
| HB | 82 | ALA | HB | H |
| 1.255 | 0.020 | 1 | | |
| N | 82 | ALA | N | N |
| 124.204 | 0.3 | 1 | | |
| H | 83 | ARG | H | H |
| 7.639 | 0.020 | 1 | | |
| HA | 83 | ARG | HA | H |
| 3.965 | 0.020 | 1 | | |
| HB2 | 83 | ARG | HB2 | H |
| 1.603 | 0.020 | 1 | | |
| HB3 | 83 | ARG | HB3 | H |
| 1.603 | 0.020 | 1 | | |
| HE | 83 | ARG | HE | H |
| 2.572 | 0.020 | 1 | | |
| N | 83 | ARG | N | N |
| 119.956 | 0.3 | 1 | | |
| CB | 84 | ASP | CB | C |
| 39.157 | 0.3 | 1 | | |
| H | 84 | ASP | H | H |
| 7.807 | 0.020 | 1 | | |
| HA | 84 | ASP | HA | H |
| 4.241 | 0.020 | 1 | | |
| HB2 | 84 | ASP | HB2 | H |
| 2.517 | 0.020 | 1 | | |
| HB3 | 84 | ASP | HB3 | H |
| 2.517 | 0.020 | 1 | | |
| N | 84 | ASP | N | N |
| 126.487 | 0.3 | 1 | | |

stop_

**Table S4**: Nuclear spin relaxation data for apo TRBP2-dsRBD2 recorded at 600 MHz and 800 MHz NMR spectrometer. Data for some residues is missing in this table due to line-broadening issues in the corresponding experiments.

| Residue Number | 600 MHz | | | | | | 800 MHz | | | | | |
|---|---|---|---|---|---|---|---|---|---|---|---|---|
| | $R_1$ ($s^{-1}$) | | $R_2$ ($s^{-1}$) | | NOE | | $R1$ ($s^{-1}$) | | $R_2$ ($s^{-1}$) | | NOE | |
| | Value | Error | Value | Error | Value | Error | Value | Error | Value | Error | Value | Error |
| S151 | 1.39 | 0.04 | 3.32 | 0.05 | -0.31 | 0.01 | 1.34 | 0.04 | 4.98 | 1.40 | -0.07 | 0.01 |
| N152 | 1.41 | 0.07 | 4.23 | 0.09 | -13 | 0.01 | 1.41 | 0.05 | 5.67 | 1.41 | 0.38 | 0.01 |
| A153 | 1.38 | 0.07 | 5.36 | 0.08 | 0.12 | 0.01 | 1.42 | 0.02 | 5.91 | 0.33 | 0.38 | 0.01 |
| Q154 | 1.45 | 0.05 | 4.02 | 0.05 | 0.09 | 0.01 | 1.48 | 0.04 | 4.73 | 0.62 | 0.40 | 0.01 |
| Q155 | 1.33 | 0.05 | 3.60 | 0.1 | 0.14 | 0.01 | 1.37 | 0.03 | 4.5 | 0.89 | 0.77 | 0.01 |
| S156 | 1.45 | 0.06 | 4.49 | 0.06 | -0.05 | 0.01 | 1.46 | 0.03 | 6.24 | 1.42 | 0.25 | 0.01 |
| S157 | 1.51 | 0.06 | 5.22 | 0.13 | 0.23 | 0.01 | 1.51 | 0.05 | 6.57 | 1.27 | 0.39 | 0.01 |
| N159 | 1.38 | 0.04 | 7.64 | 0.18 | 0.48 | 0.03 | 1.26 | 0.06 | 8.61 | 1.59 | 0.61 | 0.03 |
| G162 | 1.45 | 0.02 | 10.86 | 0.14 | 0.78 | 0.02 | 1.20 | 0.03 | 10.20 | 0.74 | 0.84 | 0.02 |
| A163 | 1.32 | 0.04 | 11.55 | 0.28 | 0.81 | 0.04 | 1.22 | 0.02 | 12.27 | 0.47 | 0.82 | 0.02 |
| L164 | 1.40 | 0.02 | 12.54 | 0.19 | 0.83 | 0.02 | 1.11 | 0.03 | 12.25 | 0.38 | 0.85 | 0.02 |
| Q165 | 1.28 | 0.06 | 12.66 | 0.11 | 0.82 | 0.02 | 1.24 | 0.01 | 12.03 | 0.46 | 0.84 | 0.01 |
| E166 | 1.43 | 0.02 | 11.88 | 0.11 | 0.81 | 0.01 | 1.12 | 0.02 | 11.40 | 0.24 | 0.84 | 0.01 |
| L167 | 1.35 | 0.06 | 12.41 | 0.45 | 0.76 | 0.04 | 1.12 | 0.01 | 12.18 | 0.11 | 0.79 | 0.02 |
| V168 | 1.37 | 0.06 | 21.02 | 1.17 | 0.92 | 0.06 | 1.10 | 0.03 | - | - | 0.88 | 0.04 |
| V169 | 1.46 | 0.09 | 10.85 | 0.73 | 0.69 | 0.04 | 1.17 | 0.02 | 11.68 | 0.31 | 0.87 | 0.04 |
| Q170 | 1.45 | 0.02 | 12.11 | 0.25 | 0.80 | 0.01 | 1.10 | 0.02 | 11.69 | 0.10 | 0.82 | 0.01 |
| G172 | 1.44 | 0.05 | 10.56 | 0.89 | 0.76 | 0.02 | 1.21 | 0.01 | 10.80 | 0.32 | 0.81 | 0.02 |
| W173 | 1.29 | 0.10 | 10.44 | 0.30 | 0.80 | 0.03 | 1.22 | 0.01 | 10.15 | 0.16 | 0.84 | 0.02 |
| R174 | 1.36 | 0.03 | 9.48 | 0.32 | 0.8 | 0.05 | 1.11 | 0.03 | 10.16 | 0.13 | 0.78 | 0.03 |
| L175 | 1.46 | 0.04 | 10.70 | 0.1 | 0.80 | 0.02 | 1.18 | 0.02 | 10.09 | 0.13 | 0.80 | 0.02 |
| E177 | 1.39 | 0.02 | 4.99 | 0.39 | 0.57 | 0.01 | 1.15 | 0.04 | 9.65 | 0.56 | 0.59 | 0.01 |
| Y178 | 1.45 | 0.07 | 9.60 | 0.66 | 0.74 | 0.03 | 1.32 | 0.02 | 10.13 | 0.60 | 0.80 | 0.02 |
| T179 | 1.50 | 0.03 | 10.54 | 00.12 | 0.72 | 0.02 | 1.25 | 0.03 | 10.69 | 0.69 | 0.80 | 0.02 |
| V180 | 1.54 | 0.09 | 9.89 | 1.48 | 0.68 | 0.04 | 1.42 | 0.05 | 12.93 | 1.34 | 0.88 | 0.05 |
| T181 | 1.56 | 0.02 | 10.22 | 0.31 | 0.78 | 0.02 | 1.28 | 0.04 | 10.42 | 0.64 | 0.85 | 0.02 |
| Q182 | 1.54 | 0.03 | 11.54 | 0.37 | 0.71 | 0.02 | 1.32 | 0.04 | 12.09 | 0.85 | 0.79 | 0.02 |
| E183 | 1.43 | 0.03 | 10.61 | 0.08 | 0.65 | 0.01 | 1.28 | 0.04 | 10.30 | 0.76 | 0.70 | 0.01 |
| S184 | 1.41 | 0.04 | - | - | 0.52 | 0.05 | 1.35 | 0.03 | - | - | 0.65 | 0.04 |
| G185 | 1.44 | 0.03 | 8.30 | 0.18 | 0.57 | 0.02 | 1.29 | 0.05 | 8.71 | 1.11 | 0.64 | 0.02 |
| R189 | 1.58 | 0.05 | 15.44 | 0.69 | 0.49 | 0.05 | 1.40 | 0.06 | 16.65 | 1.49 | 0.51 | 0.04 |
| E191 | 1.41 | 0.02 | 9.07 | 0.06 | 0.54 | 0.01 | 1.18 | 0.03 | 9.23 | 0.49 | 0.60 | 0.01 |
| F192 | 1.40 | 0.03 | 12.40 | 0.29 | - | - | 1.22 | 0.03 | 12.16 | 0.53 | 0.79 | 0.02 |
| T193 | 1.42 | 0.02 | 12.39 | 0.14 | 0.78 | 0.03 | 1.13 | 0.03 | 11.67 | 0.67 | 0.84 | 0.02 |
| M194 | 1.52 | 0.08 | 9.92 | 0.56 | 0.72 | 0.07 | 1.31 | 0.04 | 10.71 | 0.50 | 0.80 | 0.06 |
| T195 | 1.51 | 0.03 | 13.36 | 0.15 | 0.78 | 0.03 | 1.18 | 0.04 | 13.47 | 0.70 | 0.85 | 0.02 |
| C196 | 1.56 | 0.03 | 11.00 | 0.28 | 0.82 | 0.04 | 1.23 | 0.04 | 10.85 | 0.52 | 0.78 | 0.02 |
| R197 | 1.51 | 0.03 | 8.40 | 0.94 | 0.84 | 0.05 | 1.22 | 0.04 | 9.64 | 0.50 | 0.81 | 0.04 |

| | | | | | | | | | | | | |
|---|---|---|---|---|---|---|---|---|---|---|---|---|
| V198 | 1.54 | 0.11 | 9.52 | 0.64 | 0.73 | 0.04 | 1.20 | 0.03 | 10.49 | 0.61 | 0.82 | 0.05 |
| E199 | 1.39 | 0.09 | 10.27 | 0.15 | 0.68 | 0.02 | 1.22 | 0.02 | 9.77 | 0.29 | 0.76 | 0.01 |
| R200 | 1.31 | 0.07 | 11.83 | 0.57 | 0.63 | 0.05 | 1.07 | 0.04 | 10.69 | 0.38 | 0.82 | 0.05 |
| F201 | 1.59 | 0.04 | 10.67 | 0.13 | 0.72 | 0.01 | 1.27 | 0.03 | 10.73 | 0.38 | 0.79 | 0.02 |
| I202 | 1.55 | 0.04 | 9.82 | 0.21 | 0.74 | 0.02 | 1.19 | 0.03 | 9.86 | 0.41 | 0.86 | 0.02 |
| E203 | 1.62 | 0.08 | 10.21 | 0.13 | 0.77 | 0.01 | 1.29 | 0.02 | 9.83 | 0.44 | 0.82 | 0.02 |
| I204 | 1.33 | 0.05 | 9.87 | 0.22 | 0.68 | 0.04 | 1.11 | 0.04 | 9.54 | 0.33 | 0.72 | 0.03 |
| G205 | 1.56 | 0.03 | 10.33 | 0.10 | 0.77 | 0.02 | 1.22 | 0.04 | 10.27 | 0.39 | 0.83 | 0.02 |
| S206 | 1.50 | 0.02 | 10.37 | 0.13 | 0.72 | 0.01 | 1.16 | 0.03 | 10.55 | 0.43 | 0.80 | 0.02 |
| G207 | 1.55 | 0.07 | 10.48 | 0.66 | 0.79 | 0.01 | 1.32 | 0.01 | 11.35 | 0.72 | 0.80 | 0.02 |
| T208 | 1.54 | 0.10 | 10.31 | 0.14 | 0.75 | 0.02 | 1.37 | 0.04 | 10.68 | 1.25 | 0.83 | 0.02 |
| S209 | 1.29 | 0.09 | 6.42 | 0.69 | - | - | - | - | - | - | - | - |
| K210 | 1.60 | 0.14 | 8.69 | 0.7 | 0.75 | 0.06 | 1.30 | 0.06 | 9.23 | 1.02 | 0.85 | 0.12 |
| L212 | 1.28 | 0.05 | 12.31 | 0.44 | 0.72 | 0.04 | 1.15 | 0.04 | 12.44 | 0.88 | 0.81 | 0.02 |
| A213 | 1.50 | 0.05 | 12.52 | 0.35 | 0.88 | 0.03 | 1.22 | 0.03 | 12.33 | 0.59 | 0.82 | 0.03 |
| R215 | 1.27 | 0.04 | 12.93 | 0.41 | 0.66 | 0.06 | 1.10 | 0.02 | 12.33 | 0.34 | 0.81 | 0.03 |
| N216 | 1.37 | 0.04 | 12.83 | 0.15 | 0.83 | 0.01 | 1.13 | 0.02 | 12.74 | 0.44 | 0.83 | 0.01 |
| A217 | 1.48 | 0.02 | 11.11 | 0.36 | - | - | 1.19 | 0.03 | 12.01 | 0.73 | 0.88 | 0.03 |
| A218 | 1.45 | 0.04 | 12.37 | 0.40 | 0.79 | 0.02 | 1.19 | 0.02 | 12.47 | 0.40 | 0.81 | 0.04 |
| A219 | 1.44 | 0.04 | 12.61 | 0.18 | 0.84 | 0.02 | 1.13 | 0.03 | 12.36 | 0.31 | 0.88 | 0.03 |
| M221 | 1.46 | 0.03 | 8.57 | 0.22 | 0.5 | 0.02 | 1.25 | 0.06 | 9.11 | 1.29 | 0.49 | 0.02 |
| L222 | 1.56 | 0.02 | - | - | 0.79 | 0.03 | 1.13 | 0.02 | 12.70 | 0.26 | 0.86 | 0.02 |
| L223 | 1.31 | 0.03 | 12.49 | 0.53 | 0.65 | 0.03 | 1.07 | 0.01 | 12.44 | 0.24 | 0.87 | 0.02 |
| R224 | 1.20 | 0.06 | 11.11 | 0.90 | 0.79 | 0.05 | 1.15 | 0.03 | 11.33 | 0.36 | 0.84 | 0.03 |
| V225 | 1.30 | 0.09 | 12.12 | 0.32 | 0.76 | 0.05 | 1.16 | 0.01 | 11.29 | 0.38 | 0.79 | 0.04 |
| T227 | 1.50 | 0.04 | 8.77 | 0.16 | 0.54 | 0.02 | 1.39 | 0.04 | 9.60 | 1.33 | 0.66 | 0.02 |
| V228 | 1.57 | 0.04 | 7.61 | 0.19 | 0.57 | 0.02 | 1.36 | 0.05 | 7.88 | 0.59 | 0.64 | 0.02 |
| L230 | 1.01 | 0.08 | 1.62 | 0.41 | -0.65 | 0.00 | 0.95 | 0.02 | 1.97 | 0.37 | -0.81 | 0.01 |
| A232 | 1.44 | 0.05 | 4.08 | 0.06 | -0.36 | -0.02 | 1.31 | 0.04 | 4.54 | 0.61 | 0.29 | 0.02 |

**Table S5**: *R₂eff* values measured at different CPMG frequencies from CPMG relaxation dispersion experiment for apo TRBP2-dsRBD2 at 600 MHz NMR spectrometer.

| Residue Number | $R_{2eff}$ at CPMG frequency ($s^{-1}$) | | | | | | | | | | | | | |
|---|---|---|---|---|---|---|---|---|---|---|---|---|---|---|
| | | | | | | | | | | | | Repeat | | |
| | 25 | 50 | 75 | 125 | 175 | 275 | 375 | 525 | 675 | 825 | 1000 | 125 | 375 | 825 |
| error | 0.52 | 0.48 | 0.43 | 0.44 | 0.33 | 0.33 | 0.30 | 0.30 | 0.25 | 0.25 | 0.28 | 0.38 | 0.28 | 0.28 |
| S151 | 2.95 | 5.77 | 1.83 | 5.00 | 6.54 | 4.92 | 4.35 | 6.83 | 6.54 | 7.41 | 8.30 | 4.55 | 7.14 | 7.45 |
| N152 | 1.00 | 1.58 | 1.58 | 2.70 | 1.58 | 3.48 | 4.64 | 5.68 | 6.02 | 6.59 | 6.41 | 2.64 | 2.75 | 6.75 |
| A153 | 2.84 | 3.59 | 3.67 | 3.32 | 3.83 | 4.44 | 4.50 | 4.44 | 5.03 | 5.33 | 8.28 | 3.25 | 4.97 | 5.36 |
| Q154 | 6.27 | 6.07 | 5.86 | 6.07 | 6.76 | 6.34 | 6.63 | 7.11 | 7.38 | 7.58 | 4.76 | 5.97 | 6.33 | 7.59 |
| Q155 | 2.47 | 1.93 | 2.26 | 2.56 | 2.74 | 2.79 | 3.21 | 3.27 | 4.02 | 4.46 | 5.62 | 3.07 | 2.79 | 4.51 |
| E157 | 2.47 | 2.30 | 2.41 | 2.21 | 2.07 | 4.02 | 4.26 | 4.10 | 7.20 | 5.18 | 4.97 | 2.34 | 3.93 | 5.25 |
| C158 | 1.79 | 2.10 | 3.75 | 5.53 | 4.45 | 5.53 | 6.20 | 4.11 | 5.29 | 7.71 | 8.22 | 3.24 | 3.91 | 8.05 |
| N159 | 12.80 | 14.90 | 15.90 | 3.78 | 8.93 | 7.97 | 6.66 | 12.30 | 12.60 | 14.00 | 10.70 | 6.62 | 7.44 | 14.20 |
| L167 | 9.37 | 9.58 | 9.83 | 9.79 | 9.98 | 10.80 | 11.40 | 11.70 | 11.60 | 13.30 | 13.80 | 9.82 | 11.10 | 13.40 |
| V168 | 15.40 | 15.40 | 15.90 | 16.00 | 15.70 | 16.70 | 17.50 | 17.50 | 16.50 | 18.00 | 18.50 | 15.50 | 17.80 | 17.30 |
| V169 | 8.40 | 9.12 | 8.68 | 9.27 | 8.55 | 9.03 | 9.91 | 10.00 | 10.50 | 10.70 | 11.60 | 8.85 | 9.84 | 10.70 |
| Q170 | 8.42 | 8.27 | 8.42 | 9.89 | 9.64 | 8.62 | 9.11 | 8.90 | 9.31 | 9.95 | 11.00 | 9.33 | 8.70 | 10.70 |
| G172 | 8.20 | 8.31 | 8.22 | 7.95 | 8.52 | 8.64 | 8.97 | 9.56 | 10.20 | 10.30 | 11.30 | 8.04 | 9.18 | 10.80 |
| W173 | 7.12 | 8.34 | 7.89 | 7.02 | 7.31 | 8.09 | 8.59 | 9.11 | 9.34 | 10.00 | 10.90 | 7.42 | 8.84 | 10.10 |
| L175 | 7.80 | 7.73 | 8.49 | 7.88 | 8.26 | 8.15 | 9.51 | 9.51 | 10.30 | 10.40 | 10.70 | 8.20 | 8.70 | 10.50 |
| E177 | 8.14 | 7.62 | 7.66 | 8.44 | 7.80 | 8.46 | 9.07 | 9.34 | 9.39 | 10.70 | 11.70 | 8.58 | 8.60 | 10.80 |
| Y178 | 7.78 | 6.44 | 7.96 | 7.64 | 7.72 | 8.28 | 7.62 | 84.20 | 8.77 | 11.50 | 11.20 | 8.04 | 9.44 | 11.20 |
| T179 | 8.50 | 7.54 | 8.68 | 8.93 | 9.16 | 9.30 | 9.41 | 9.64 | 11.20 | 11.00 | 11.10 | 8.91 | 8.63 | 10.90 |
| V180 | 11.20 | 10.60 | 10.10 | 9.96 | 8.91 | 9.77 | 10.10 | 7.47 | 11.20 | 11.20 | 14.40 | 9.25 | 9.55 | 11.30 |
| T181 | 8.24 | 8.59 | 9.01 | 7.14 | 8.89 | 8.04 | 8.63 | 9.86 | 9.81 | 9.64 | 10.30 | 7.46 | 8.48 | 9.27 |
| Q182 | 8.16 | 7.68 | 8.23 | 8.21 | 8.27 | 7.83 | 9.46 | 9.08 | 10.20 | 9.32 | 11.00 | 8.00 | 8.86 | 10.30 |
| E183 | 7.30 | 7.43 | 7.30 | 7.44 | 7.49 | 7.74 | 7.89 | 8.82 | 9.20 | 9.42 | 9.78 | 7.18 | 7.80 | 9.49 |
| G185 | 6.24 | 4.69 | 5.88 | 6.62 | 4.94 | 6.76 | 7.73 | 6.29 | 6.47 | 7.35 | 7.69 | 5.37 | 5.90 | 7.49 |
| A187 | 10.40 | 8.74 | 9.48 | 10.20 | 9.64 | 10.00 | 11.70 | 10.70 | 12.00 | 10.50 | 11.70 | 8.94 | 10.60 | 11.20 |
| R189 | 7.06 | 8.56 | 7.36 | 7.14 | 9.54 | 9.29 | 9.45 | 9.41 | 11.30 | 9.73 | 12.90 | 9.44 | 11.70 | 8.70 |
| E191 | 6.67 | 6.48 | 6.26 | 6.47 | 7.39 | 6.75 | 7.04 | 7.52 | 7.78 | 7.98 | 8.69 | 6.38 | 6.74 | 7.99 |
| F192 | 9.20 | 8.03 | 8.28 | 9.72 | 8.96 | 9.93 | 10.20 | 10.00 | 9.87 | 10.90 | 10.80 | 9.05 | 9.29 | 11.30 |
| T193 | 8.84 | 9.17 | 9.26 | 9.33 | 9.55 | 10.50 | 10.70 | 11.70 | 11.20 | 12.10 | 13.10 | 9.62 | 10.40 | 12.10 |
| M194 | 9.13 | 10.50 | 8.68 | 9.42 | 8.96 | 9.68 | 10.80 | 10.40 | 10.30 | 12.10 | 12.30 | 7.61 | 10.10 | 12.20 |
| T195 | 9.39 | 9.39 | 9.06 | 10.30 | 9.57 | 9.66 | 9.94 | 10.80 | 10.30 | 11.50 | 11.60 | 9.90 | 10.90 | 11.60 |
| C196 | 8.38 | 8.57 | 9.57 | 8.55 | 8.33 | 9.19 | 9.86 | 9.08 | 8.63 | 9.77 | 11.10 | 9.33 | 7.91 | 9.86 |
| R197 | 8.81 | 8.49 | 7.95 | 7.72 | 8.00 | 8.13 | 8.61 | 10.30 | 10.30 | 12.10 | 10.90 | 7.54 | 8.15 | 12.40 |
| V198 | 8.49 | 6.46 | 8.07 | 7.74 | 7.01 | 8.63 | 8.56 | 9.51 | 7.88 | 10.10 | 9.46 | 8.57 | 9.46 | 10.10 |
| E199 | 7.56 | 7.65 | 7.70 | 8.07 | 8.07 | 8.54 | 8.86 | 9.56 | 10.30 | 11.70 | 10.90 | 8.26 | 9.45 | 11.70 |
| R200 | 8.18 | 9.23 | 8.39 | 8.31 | 9.41 | 9.50 | 9.71 | 10.90 | 10.90 | 11.40 | 9.85 | 8.59 | 10.60 | 12.40 |
| F201 | 8.27 | 8.35 | 8.87 | 8.58 | 8.67 | 9.06 | 8.87 | 9.82 | 10.30 | 11.10 | 10.90 | 8.49 | 8.94 | 11.20 |
| I202 | 7.81 | 8.12 | 7.34 | 7.02 | 11.00 | 7.51 | 8.35 | 9.49 | 9.50 | 10.10 | 11.00 | 7.61 | 9.28 | 10.20 |
| E203 | 7.90 | 8.48 | 6.33 | 8.48 | 8.35 | 9.09 | 8.56 | 9.52 | 10.10 | 10.40 | 11.40 | 8.69 | 8.45 | 10.50 |

| | | | | | | | | | | | | | | |
|---|---|---|---|---|---|---|---|---|---|---|---|---|---|---|
| I204 | 9.02 | 9.29 | 8.84 | 9.87 | 9.76 | 10.90 | 10.10 | 11.30 | 10.30 | 12.40 | 12.30 | 9.58 | 11.40 | 12.40 |
| G205 | 8.43 | 8.20 | 8.01 | 8.21 | 8.36 | 9.58 | 9.18 | 9.32 | 9.99 | 10.90 | 11.30 | 8.41 | 9.27 | 11.00 |
| S206 | 8.84 | 9.17 | 9.26 | 9.33 | 9.55 | 10.30 | 10.70 | 11.70 | 11.20 | 12.10 | 12.90 | 9.62 | 10.40 | 12.60 |
| G207 | 8.30 | 7.89 | 8.00 | 8.60 | 8.09 | 9.37 | 9.90 | 9.92 | 10.40 | 11.50 | 11.90 | 8.80 | 9.72 | 11.50 |
| T208 | 9.89 | 5.44 | 7.69 | 8.33 | 6.76 | 9.71 | 7.99 | 10.60 | 9.83 | 10.10 | 11.60 | 8.63 | 10.20 | 10.20 |
| K210 | 6.15 | 7.06 | 7.04 | 11.10 | 2.35 | 6.02 | 5.54 | 12.20 | 6.76 | 10.60 | 5.55 | 10.40 | 4.12 | 11.90 |
| L212 | 9.36 | 9.27 | 8.34 | 9.36 | 10.10 | 10.10 | 10.10 | 9.88 | 11.50 | 11.90 | 13.10 | 8.94 | 10.10 | 11.80 |
| A213 | 9.71 | 9.31 | 9.59 | 9.19 | 10.10 | 10.40 | 9.98 | 11.90 | 11.80 | 12.40 | 14.00 | 9.57 | 10.70 | 12.50 |
| R215 | 10.40 | 9.84 | 9.99 | 9.94 | 10.60 | 10.40 | 10.90 | 10.30 | 11.90 | 11.50 | 12.40 | 10.00 | 9.22 | 11.40 |
| N216 | 9.56 | 9.68 | 10.40 | 10.40 | 9.96 | 10.70 | 11.10 | 10.90 | 11.10 | 12.00 | 12.90 | 9.93 | 10.80 | 12.10 |
| A217 | 8.19 | 9.23 | 8.85 | 7.99 | 7.99 | 8.92 | 9.23 | 10.80 | 10.30 | 9.47 | 11.70 | 8.14 | 9.08 | 9.52 |
| A218 | 10.20 | 9.14 | 9.03 | 11.30 | 9.65 | 10.70 | 10.50 | 10.70 | 11.00 | 12.50 | 12.50 | 10.70 | 10.30 | 12.70 |
| A219 | 10.20 | 9.39 | 9.66 | 10.80 | 9.85 | 10.90 | 11.30 | 11.50 | 11.80 | 13.90 | 13.10 | 10.20 | 10.80 | 14.30 |
| M221 | 8.32 | 7.06 | 6.00 | 7.55 | 5.65 | 9.94 | 8.42 | 10.30 | 8.80 | 9.61 | 11.80 | 6.80 | 9.88 | 9.84 |
| L222 | 9.58 | 9.89 | 9.35 | 9.40 | 9.27 | 10.00 | 10.20 | 10.70 | 10.80 | 10.30 | 10.90 | 9.33 | 9.90 | 10.50 |
| L223 | 10.40 | 9.88 | 9.78 | 10.70 | 10.50 | 11.30 | 11.00 | 11.50 | 12.90 | 13.00 | 13.30 | 10.10 | 10.90 | 13.20 |
| R224 | 8.10 | 8.61 | 8.10 | 7.94 | 2.34 | 7.70 | 7.57 | 8.59 | 10.60 | 10.40 | 10.90 | 8.99 | 8.19 | 10.30 |
| V225 | 9.13 | 9.04 | 9.91 | 8.86 | 8.92 | 11.10 | 10.10 | 9.56 | 10.70 | 12.10 | 11.30 | 9.69 | 9.63 | 12.10 |
| T227 | 6.36 | 5.50 | 5.59 | 7.49 | 6.03 | 9.35 | 8.86 | 7.62 | 7.32 | 8.20 | 9.06 | 7.06 | 7.74 | 8.43 |
| V228 | 4.41 | 4.03 | 5.10 | 4.15 | 4.63 | 5.91 | 4.19 | 4.36 | 6.78 | 6.53 | 7.35 | 4.45 | 5.22 | 6.44 |
| L230 | 0.15 | 1.66 | 0.21 | 0.85 | 0.56 | 1.48 | 1.59 | 2.60 | 3.21 | 4.06 | 4.38 | 1.81 | 2.66 | 4.39 |
| A232 | 0.91 | 0.82 | 0.50 | 1.24 | 1.04 | 1.98 | 2.04 | 2.67 | 3.21 | 5.87 | 5.78 | 0.57 | 2.64 | 4.22 |

**Table S6**: $R_{1\rho}$ relaxation rates measured using HSn pulses (n=1,2,4,6,8) from HARD experiment for apo TRBP2-dsRBD2 at 600 MHz NMR spectrometer.

| Residue Number | $R_{1\rho}$ ($s^{-1}$) | | | | | | | | | |
|---|---|---|---|---|---|---|---|---|---|---|
| | HS1 | | HS2 | | HS4 | | HS6 | | HS8 | |
| | Value | Error | Value | Error | Value | Error | Value | Error | Value | Error |
| S151 | 3.76 | 0.39 | 3.67 | 0.29 | 4.18 | 0.25 | 4.01 | 0.12 | 4.86 | 0.54 |
| Q154 | 4.78 | 0.30 | 4.40 | 0.30 | 4.93 | 0.36 | 5.19 | 0.13 | 6.07 | 0.42 |
| Q155 | 5.14 | 0.23 | 5.02 | 0.32 | 5.13 | 0.43 | 5.51 | 0.10 | 6.06 | 0.28 |
| S156 | 4.37 | 0.53 | 4.26 | 0.33 | 5.08 | 0.32 | 4.78 | 0.18 | 5.87 | 0.44 |
| E157 | 4.68 | 0.38 | 4.34 | 0.25 | 4.94 | 0.31 | 5.02 | 0.11 | 6.02 | 0.48 |
| N159 | 4.12 | 0.56 | 4.06 | 0.45 | 4.42 | 0.19 | 5.01 | 0.25 | 5.92 | 0.50 |
| G162 | 4.75 | 0.27 | 5.34 | 0.29 | 6.05 | 0.14 | 6.31 | 0.13 | 7.00 | 0.39 |
| A163 | 5.54 | 0.44 | 5.55 | 0.42 | 6.81 | 0.29 | 6.80 | 0.16 | 8.30 | 0.64 |
| Q165 | 4.84 | 0.39 | 5.16 | 0.32 | 6.06 | 0.16 | 6.32 | 0.15 | 7.44 | 0.43 |
| E166 | 4.86 | 0.34 | 5.54 | 0.24 | 6.35 | 0.23 | 6.44 | 0.10 | 7.60 | 0.40 |
| L167 | 5.35 | 0.23 | 5.87 | 0.29 | 6.88 | 0.35 | 6.99 | 0.19 | 7.95 | 0.43 |
| V168 | 5.82 | 0.46 | 6.73 | 0.35 | 8.11 | 0.27 | 8.62 | 0.25 | 9.90 | 0.49 |
| V169 | 4.79 | 0.28 | 5.05 | 0.34 | 6.09 | 0.27 | 6.57 | 0.21 | 7.40 | 0.29 |
| Q170 | 5.78 | 0.42 | 6.29 | 0.23 | 6.94 | 0.33 | 7.34 | 0.15 | 8.34 | 0.39 |
| G172 | 4.34 | 0.32 | 4.35 | 0.31 | 5.27 | 0.21 | 5.57 | 0.09 | 6.42 | 0.34 |
| W173 | 4.70 | 0.35 | 4.97 | 0.17 | 5.91 | 0.23 | 5.88 | 0.13 | 6.71 | 0.43 |
| E177 | 3.96 | 0.25 | 4.01 | 0.25 | 4.89 | 0.22 | 4.92 | 0.10 | 5.70 | 0.36 |
| Y178 | 4.05 | 0.29 | 4.33 | 0.23 | 4.88 | 0.16 | 5.27 | 0.11 | 5.95 | 0.37 |
| T179 | 4.31 | 0.32 | 5.02 | 0.25 | 5.53 | 0.14 | 5.86 | 0.13 | 6.65 | 0.35 |
| V180 | 4.67 | 0.33 | 4.57 | 0.19 | 5.94 | 0.29 | 6.00 | 0.15 | 6.69 | 0.47 |
| T181 | 3.86 | 0.26 | 4.35 | 0.22 | 5.09 | 0.17 | 5.18 | 0.11 | 5.91 | 0.29 |
| Q182 | 5.05 | 0.34 | 5.59 | 0.24 | 6.12 | 0.27 | 6.42 | 0.10 | 7.41 | 0.36 |
| E183 | 4.67 | 0.32 | 4.72 | 0.36 | 5.66 | 0.26 | 5.57 | 0.11 | 6.49 | 0.41 |
| G185 | 5.01 | 0.25 | 5.02 | 0.30 | 5.75 | 0.32 | 5.56 | 0.27 | 6.71 | 0.35 |
| R189 | 4.72 | 0.45 | 4.91 | 0.27 | 6.38 | 0.13 | 6.23 | 0.18 | 7.60 | 0.50 |
| E191 | 4.06 | 0.32 | 4.03 | 0.26 | 4.88 | 0.25 | 5.07 | 0.10 | 5.95 | 0.40 |
| T193 | 4.49 | 0.31 | 5.08 | 0.23 | 5.85 | 0.24 | 5.99 | 0.11 | 6.91 | 0.33 |
| M194 | 4.88 | 0.18 | 4.94 | 0.36 | 6.04 | 0.26 | 6.12 | 0.26 | 6.44 | 0.58 |
| T195 | 4.96 | 0.29 | 5.35 | 0.19 | 6.21 | 0.21 | 6.31 | 0.07 | 7.24 | 0.34 |
| C196 | 4.51 | 0.36 | 5.17 | 0.33 | 5.68 | 0.14 | 5.93 | 0.18 | 6.79 | 0.37 |
| R197 | 5.29 | 0.32 | 5.36 | 0.20 | 6.01 | 0.36 | 6.43 | 0.22 | 6.90 | 0.34 |
| V198 | 3.99 | 0.48 | 4.03 | 0.10 | 4.92 | 0.22 | 5.12 | 0.24 | 5.77 | 0.33 |
| E199 | 4.53 | 0.37 | 4.68 | 0.28 | 5.62 | 0.26 | 5.62 | 0.12 | 6.74 | 0.41 |
| R200 | 4.33 | 0.29 | 5.36 | 0.36 | 6.01 | 0.35 | 6.09 | 0.23 | 7.10 | 0.29 |
| F201 | 4.96 | 0.35 | 5.18 | 0.25 | 6.02 | 0.28 | 6.26 | 0.13 | 6.96 | 0.35 |
| I202 | 4.49 | 0.24 | 4.98 | 0.23 | 5.67 | 0.19 | 5.89 | 0.20 | 6.54 | 0.32 |
| E203 | 5.04 | 0.29 | 5.33 | 0.24 | 5.89 | 0.22 | 6.00 | 0.12 | 6.95 | 0.34 |
| I204 | 5.08 | 0.27 | 5.51 | 0.17 | 6.11 | 0.37 | 6.07 | 0.11 | 7.13 | 0.24 |
| G205 | 4.37 | 0.27 | 4.91 | 0.19 | 5.68 | 0.25 | 6.04 | 0.14 | 6.68 | 0.32 |

| | | | | | | | | | | |
|---|---|---|---|---|---|---|---|---|---|---|
| S206 | 4.29 | 0.36 | 4.36 | 0.27 | 5.13 | 0.14 | 5.37 | 0.12 | 6.08 | 0.38 |
| G207 | 4.48 | 0.34 | 4.76 | 0.32 | 5.42 | 0.20 | 5.87 | 0.11 | 6.73 | 0.45 |
| T208 | 5.53 | 0.47 | 5.54 | 0.46 | 6.29 | 0.27 | 6.33 | 0.33 | 7.28 | 0.49 |
| S209 | 3.64 | 0.36 | 3.38 | 0.50 | 4.04 | 0.59 | 4.15 | 0.46 | 5.07 | 0.44 |
| L212 | 4.76 | 0.34 | 4.97 | 0.35 | 6.13 | 0.23 | 6.20 | 0.26 | 7.52 | 0.35 |
| A213 | 4.74 | 0.46 | 4.83 | 0.26 | 5.81 | 0.19 | 6.16 | 0.14 | 7.21 | 0.54 |
| R215 | 4.81 | 0.46 | 5.38 | 0.35 | 6.39 | 0.21 | 6.76 | 0.29 | 7.82 | 0.58 |
| N216 | 4.95 | 0.40 | 5.55 | 0.36 | 6.48 | 0.24 | 6.64 | 0.19 | 7.76 | 0.46 |
| A217 | 4.34 | 0.44 | 4.93 | 0.27 | 6.06 | 0.22 | 5.91 | 0.22 | 7.05 | 0.40 |
| A218 | 4.93 | 0.36 | 5.26 | 0.35 | 6.16 | 0.21 | 6.64 | 0.29 | 7.68 | 0.54 |
| A219 | 4.67 | 0.36 | 5.03 | 0.41 | 6.01 | 0.20 | 6.26 | 0.19 | 7.42 | 0.44 |
| L223 | 4.88 | 0.35 | 5.64 | 0.18 | 6.50 | 0.31 | 6.74 | 0.20 | 7.79 | 0.34 |
| R224 | 4.56 | 0.22 | 4.66 | 0.34 | 5.56 | 0.21 | 5.91 | 0.21 | 6.93 | 0.58 |
| V225 | 5.57 | 0.24 | 5.62 | 0.32 | 6.48 | 0.36 | 6.74 | 0.21 | 7.56 | 0.63 |
| T227 | 4.44 | 0.25 | 4.70 | 0.19 | 5.44 | 0.34 | 5.72 | 0.14 | 6.65 | 0.44 |
| V228 | 5.38 | 0.24 | 5.29 | 0.24 | 5.89 | 0.31 | 5.83 | 0.08 | 6.96 | 0.46 |
| L230 | 3.33 | 0.43 | 2.93 | 0.33 | 3.54 | 0.19 | 3.51 | 0.18 | 4.33 | 0.57 |
| A232 | 4.65 | 0.40 | 4.27 | 0.31 | 4.83 | 0.25 | 4.83 | 0.17 | 5.84 | 0.57 |

**Table S7**: $R_{2\rho}$ relaxation rates measured using HSn pulses (n=1,2,4,6,8) from HARD experiment for apo TRBP2-dsRBD2 at 600 MHz NMR spectrometer.

| Residue Number | $R_{2\rho}$ ($s^{-1}$) | | | | | | | | | |
|---|---|---|---|---|---|---|---|---|---|---|
| | HS1 | | HS2 | | HS4 | | HS6 | | HS8 | |
| | Value | Error | Value | Error | Value | Error | Value | Error | Value | Error |
| S151 | 7.22 | 0.25 | 7.20 | 0.35 | 6.78 | 0.07 | 7.27 | 0.19 | 7.30 | 0.64 |
| Q154 | 6.78 | 0.24 | 6.98 | 0.15 | 6.55 | 0.36 | 6.54 | 0.35 | 7.20 | 0.22 |
| Q155 | 5.75 | 0.74 | 5.06 | 0.11 | 4.86 | 0.18 | 6.35 | 1.02 | 6.09 | 0.41 |
| S156 | 9.58 | 0.68 | 9.00 | 0.63 | 8.43 | 0.51 | 7.46 | 0.45 | 8.63 | 0.49 |
| E157 | 7.68 | 0.72 | 7.75 | 0.13 | 7.05 | 0.45 | 7.44 | 0.81 | 7.78 | 0.40 |
| N159 | 12.73 | 0.23 | 14.44 | 0.79 | 12.52 | 1.01 | 10.49 | 0.77 | 14.41 | 0.99 |
| G162 | 12.32 | 0.37 | 12.14 | 0.46 | 12.59 | 0.03 | 12.88 | 0.09 | 12.30 | 0.35 |
| A163 | 13.65 | 0.91 | 13.41 | 0.62 | 11.97 | 0.31 | 12.47 | 1.28 | 14.41 | 0.35 |
| L164 | 14.55 | 0.24 | 13.28 | 0.30 | 13.33 | 0.16 | 13.83 | 0.03 | 12.48 | 0.32 |
| Q165 | 14.05 | 0.55 | 13.32 | 0.41 | 12.93 | 0.65 | 13.00 | 0.78 | 13.00 | 0.18 |
| E166 | 12.61 | 0.35 | 12.51 | 0.03 | 11.74 | 0.28 | 12.00 | 0.51 | 12.45 | 0.12 |
| L167 | 13.39 | 0.76 | 12.91 | 0.35 | 12.96 | 0.25 | 13.32 | 0.41 | 12.99 | 0.86 |
| V168 | 20.39 | 1.92 | 19.34 | 0.25 | 16.87 | 0.79 | 18.63 | 2.04 | 19.05 | 0.87 |
| V169 | 14.19 | 0.95 | 13.55 | 0.54 | 12.16 | 0.88 | 12.45 | 0.95 | 14.13 | 0.70 |
| Q170 | 13.85 | 0.07 | 13.71 | 0.50 | 13.03 | 0.30 | 13.10 | 0.39 | 13.28 | 0.49 |
| G172 | 12.51 | 0.49 | 11.90 | 0.31 | 12.13 | 0.76 | 11.81 | 0.96 | 11.88 | 0.14 |
| W173 | 11.85 | 0.39 | 11.63 | 0.59 | 11.42 | 0.25 | 10.98 | 0.69 | 11.25 | 0.26 |
| R174 | 11.73 | 0.24 | 10.27 | 0.78 | 10.40 | 0.68 | 10.23 | 0.49 | 9.29 | 0.15 |
| E177 | 10.83 | 0.46 | 10.81 | 0.18 | 10.60 | 0.31 | 10.36 | 0.32 | 10.61 | 0.33 |
| Y178 | 11.61 | 0.04 | 10.91 | 0.37 | 10.97 | 0.47 | 10.72 | 0.55 | 10.75 | 0.39 |
| T179 | 11.63 | 0.34 | 11.39 | 0.59 | 11.19 | 0.21 | 11.14 | 0.27 | 11.19 | 0.37 |
| V180 | 15.96 | 0.78 | 14.19 | 1.10 | 13.22 | 0.40 | 13.27 | 0.85 | 13.50 | 0.63 |
| T181 | 11.86 | 0.29 | 11.37 | 0.22 | 10.92 | 0.12 | 10.72 | 0.37 | 11.02 | 0.37 |
| Q182 | 12.69 | 0.18 | 12.46 | 0.52 | 12.48 | 0.22 | 12.44 | 0.27 | 12.36 | 0.44 |
| E183 | 11.39 | 0.60 | 10.75 | 0.20 | 10.48 | 0.48 | 10.83 | 0.68 | 10.99 | 0.46 |
| G185 | 9.43 | 0.58 | 9.59 | 0.34 | 8.82 | 0.46 | 9.58 | 0.32 | 10.15 | 0.25 |
| R189 | 15.64 | 1.98 | 14.98 | 0.94 | 14.67 | 0.66 | 13.13 | 1.48 | 13.28 | 1.04 |
| E191 | 10.71 | 0.42 | 9.96 | 0.37 | 9.94 | 0.21 | 10.04 | 0.31 | 9.51 | 0.12 |
| F192 | 13.38 | 0.07 | 12.92 | 0.36 | 13.11 | 0.10 | 12.54 | 0.36 | 12.90 | 0.12 |
| T193 | 12.72 | 0.50 | 12.80 | 0.72 | 12.45 | 0.48 | 12.58 | 0.32 | 12.45 | 0.66 |
| M194 | 8.43 | 0.49 | 9.32 | 1.43 | 8.84 | 0.51 | 9.21 | 0.28 | 10.98 | 1.37 |
| T195 | 13.09 | 0.38 | 13.14 | 0.79 | 13.18 | 0.39 | 13.38 | 0.18 | 12.72 | 0.47 |
| C196 | 12.58 | 0.35 | 12.68 | 0.32 | 11.60 | 0.80 | 10.74 | 0.74 | 12.52 | 0.86 |
| R197 | 10.37 | 0.70 | 11.05 | 1.01 | 10.43 | 0.25 | 10.60 | 0.96 | 11.38 | 0.36 |
| V198 | 13.26 | 1.36 | 11.51 | 1.40 | 12.37 | 1.02 | 10.36 | 1.86 | 10.64 | 0.94 |
| E199 | 11.88 | 0.19 | 11.36 | 0.19 | 11.15 | 0.13 | 11.10 | 0.51 | 11.18 | 0.29 |
| R200 | 12.68 | 2.54 | 10.82 | 1.71 | 10.99 | 1.47 | 11.10 | 1.08 | 11.60 | 2.00 |
| F201 | 11.48 | 0.45 | 11.38 | 0.11 | 10.67 | 0.38 | 10.77 | 0.58 | 11.34 | 0.09 |
| I202 | 11.36 | 0.79 | 11.19 | 0.64 | 10.99 | 0.51 | 10.34 | 0.47 | 10.68 | 0.16 |

| | | | | | | | | | | |
|---|---|---|---|---|---|---|---|---|---|---|
| E203 | 11.67 | 0.56 | 12.18 | 0.20 | 10.82 | 0.26 | 10.92 | 0.62 | 11.83 | 0.32 |
| I204 | 10.53 | 0.64 | 11.15 | 0.29 | 10.62 | 0.32 | 10.66 | 0.48 | 10.75 | 0.29 |
| G205 | 11.98 | 0.40 | 11.92 | 0.21 | 11.30 | 0.23 | 11.16 | 0.48 | 11.37 | 0.07 |
| S206 | 11.12 | 0.13 | 10.59 | 0.33 | 10.53 | 0.05 | 11.01 | 0.14 | 10.66 | 0.15 |
| G207 | 13.23 | 0.39 | 12.35 | 0.42 | 12.69 | 0.53 | 12.58 | 0.86 | 12.33 | 0.29 |
| T208 | 12.86 | 0.74 | 12.63 | 0.55 | 11.74 | 0.46 | 12.86 | 0.66 | 11.34 | 0.23 |
| S209 | 9.60 | 1.46 | 9.31 | 0.23 | 11.16 | 0.32 | 10.41 | 0.63 | 11.80 | 1.09 |
| L212 | 14.93 | 1.52 | 14.22 | 0.75 | 13.55 | 0.97 | 12.83 | 0.96 | 13.68 | 0.85 |
| A213 | 12.78 | 0.53 | 11.71 | 0.50 | 12.23 | 0.90 | 12.87 | 0.12 | 12.54 | 0.72 |
| R215 | 14.78 | 1.64 | 13.74 | 1.27 | 14.28 | 0.93 | 13.86 | 0.40 | 14.89 | 1.57 |
| N216 | 14.09 | 0.92 | 14.09 | 0.39 | 12.96 | 0.44 | 12.77 | 0.70 | 13.37 | 0.41 |
| A217 | 13.10 | 0.42 | 12.05 | 0.69 | 12.37 | 0.47 | 12.54 | 0.89 | 12.05 | 0.10 |
| A218 | 14.84 | 0.80 | 13.03 | 0.29 | 13.80 | 0.43 | 14.71 | 0.56 | 13.03 | 0.47 |
| A219 | 14.93 | 0.28 | 13.18 | 0.34 | 13.53 | 0.40 | 13.84 | 0.49 | 12.58 | 0.37 |
| L222 | 13.60 | 0.24 | 13.10 | 0.38 | 12.83 | 0.05 | 12.41 | 0.37 | 12.52 | 0.13 |
| L223 | 13.32 | 0.20 | 13.13 | 1.16 | 12.79 | 0.21 | 13.05 | 0.35 | 13.32 | 0.69 |
| R224 | 15.29 | 2.02 | 14.00 | 1.41 | 14.32 | 0.30 | 13.10 | 0.98 | 12.28 | 1.27 |
| V225 | 12.72 | 1.02 | 12.95 | 0.62 | 12.63 | 0.80 | 12.77 | 0.43 | 12.13 | 1.31 |
| T227 | 10.96 | 1.32 | 11.29 | 0.74 | 10.76 | 1.18 | 10.05 | 1.02 | 10.45 | 0.41 |
| V228 | 9.46 | 0.64 | 9.48 | 0.49 | 8.37 | 0.61 | 8.64 | 0.76 | 9.16 | 0.61 |
| L230 | 4.07 | 0.15 | 3.97 | 0.17 | 4.03 | 0.18 | 4.51 | 0.46 | 4.65 | 0.14 |
| A232 | 6.18 | 0.31 | 6.18 | 0.21 | 5.37 | 0.30 | 5.70 | 0.80 | 6.94 | 0.23 |

**Table S8**: $R_1$ relaxation rates measured from HARD experiment for apo TRBP2-dsRBD2 at 600 MHz NMR spectrometer.

| Residue Number | $R_1$ ($s^{-1}$) | |
|---|---|---|
| | Value | Error |
| S151 | 1.50 | 0.04 |
| Q154 | 1.57 | 0.07 |
| Q155 | 1.41 | 0.06 |
| S156 | 1.60 | 0.05 |
| E157 | 1.65 | 0.04 |
| N159 | 1.54 | 0.11 |
| G162 | 1.54 | 0.05 |
| A163 | 1.49 | 0.08 |
| Q165 | 1.50 | 0.05 |
| E166 | 1.49 | 0.05 |
| L167 | 1.44 | 0.07 |
| V168 | 1.42 | 0.08 |
| V169 | 1.51 | 0.05 |
| Q170 | 1.48 | 0.06 |
| G172 | 1.50 | 0.06 |
| W173 | 1.50 | 0.06 |
| E177 | 1.46 | 0.06 |
| Y178 | 1.55 | 0.05 |
| T179 | 1.56 | 0.04 |
| V180 | 1.71 | 0.08 |
| T181 | 1.62 | 0.04 |
| Q182 | 1.63 | 0.05 |
| E183 | 1.57 | 0.07 |
| G185 | 1.57 | 0.05 |
| R189 | 1.72 | 0.07 |
| E191 | 1.46 | 0.05 |
| T193 | 1.49 | 0.04 |
| M194 | 1.58 | 0.07 |
| T195 | 1.57 | 0.04 |
| C196 | 1.62 | 0.04 |
| R197 | 1.63 | 0.09 |
| V198 | 1.55 | 0.08 |
| E199 | 1.56 | 0.06 |
| R200 | 1.44 | 0.05 |
| F201 | 1.60 | 0.05 |
| I202 | 1.52 | 0.07 |
| E203 | 1.61 | 0.04 |
| I204 | 1.47 | 0.07 |
| G205 | 1.62 | 0.04 |
| S206 | 1.50 | 0.04 |

| | | |
|---|---|---|
| G207 | 1.55 | 0.04 |
| T208 | 1.71 | 0.04 |
| S209 | 1.35 | 0.05 |
| L212 | 1.42 | 0.04 |
| A213 | 1.53 | 0.06 |
| R215 | 1.42 | 0.08 |
| N216 | 1.44 | 0.06 |
| A217 | 1.53 | 0.09 |
| A218 | 1.45 | 0.06 |
| A219 | 1.42 | 0.05 |
| L223 | 1.35 | 0.06 |
| R224 | 1.45 | 0.08 |
| V225 | 1.33 | 0.10 |
| T227 | 1.63 | 0.05 |
| V228 | 1.67 | 0.07 |
| L230 | 1.06 | 0.06 |
| A232 | 1.47 | 0.06 |

**Table S9**: Nuclear spin relaxation data for RNA-bound TRBP2-dsRBD2 recorded at 600 MHz and 800 MHz NMR spectrometer. Data for some residues is missing in this table due to line-broadening issues in the corresponding experiments.

| Residue Number | 600 MHz | | | | | | 800 MHz | | | | | |
|---|---|---|---|---|---|---|---|---|---|---|---|---|
| | $R_1$ ($s^{-1}$) | | $R_2$ ($s^{-1}$) | | NOE | | $R_1$ ($s^{-1}$) | | $R_2$ ($s^{-1}$) | | NOE | |
| | Value | Error | Value | Error | Value | Error | Value | Error | Value | Error | Value | Error |
| S151 | 1.48 | 0.10 | 7.55 | 0.29 | -0.26 | -0.02 | 1.32 | 0.25 | 7.24 | 0.30 | 0.10 | 0.03 |
| Q154 | 1.57 | 0.02 | 6.12 | 0.06 | - | - | 1.40 | 0.13 | 5.76 | 0.25 | 0.38 | 0.01 |
| Q155 | 1.39 | 0.03 | 4.68 | 0.10 | - | - | 1.25 | 0.14 | 5.54 | 0.09 | - | - |
| S156 | 1.50 | 0.13 | 10.32 | 0.26 | 0.04 | 0.02 | 1.39 | 0.27 | 11.19 | 0.26 | 0.33 | 0.03 |
| E157 | 1.74 | 0.11 | 11.13 | 0.28 | 0.26 | 0.02 | 1.45 | 0.22 | 11.59 | 0.41 | 0.39 | 0.03 |
| N159 | 1.39 | 0.15 | 17.27 | 1.13 | 0.38 | 0.07 | 1.54 | 0.27 | 21.17 | 1.04 | 0.55 | 0.14 |
| G162 | 1.32 | 0.07 | 23.58 | 0.97 | 0.85 | 0.07 | 1.19 | 0.10 | 24.71 | 2.09 | 0.81 | 0.15 |
| A163 | 1.28 | 0.10 | 22.33 | 2.50 | 0.86 | 0.09 | 0.88 | 0.12 | 25.06 | 2.61 | 1.01 | 0.16 |
| Q165 | 1.29 | 0.06 | 24.36 | 0.97 | 0.76 | 0.05 | 1.02 | 0.10 | 24.87 | 0.54 | 0.85 | 0.07 |
| E166 | 1.35 | 0.04 | 23.64 | 0.74 | 0.74 | 0.03 | 1.04 | 0.09 | 24.39 | 0.35 | 0.77 | 0.06 |
| L167 | 1.34 | 0.06 | 22.40 | 0.39 | 0.76 | 0.07 | 0.86 | 0.07 | 26.05 | 0.62 | 0.73 | 0.13 |
| V168 | 1.27 | 0.07 | 24.52 | 3.18 | 0.71 | 0.11 | 0.96 | 0.17 | 33.08 | 2.23 | 0.63 | 0.22 |
| V169 | 1.27 | 0.10 | 14.23 | 1.55 | 0.77 | 0.17 | 1.16 | 0.10 | 12.32 | 1.64 | 0.69 | 0.22 |
| Q170 | 1.29 | 0.05 | 22.32 | 0.55 | 0.74 | 0.03 | 1.01 | 0.07 | 23.28 | 0.82 | 0.76 | 0.06 |
| G172 | 1.38 | 0.04 | 23.28 | 0.60 | 0.77 | 0.04 | 1.00 | 0.09 | 27.34 | 1.36 | 0.68 | 0.07 |
| W173 | 1.41 | 0.03 | 19.33 | 0.84 | 0.74 | 0.05 | 1.01 | 0.07 | 21.83 | 0.84 | 0.82 | 0.07 |
| E177 | 1.30 | 0.05 | 7.35 | 0.79 | 0.31 | 0.02 | 1.11 | 0.15 | 8.38 | 0.93 | 0.66 | 0.05 |
| Y178 | 1.43 | 0.04 | 21.96 | 1.13 | 0.74 | 0.06 | 0.89 | 0.08 | 22.49 | 2.11 | 0.93 | 0.11 |
| T179 | 1.38 | 0.06 | 20.72 | 0.42 | 0.82 | 0.05 | 0.90 | 0.17 | 23.41 | 0.64 | 0.68 | 0.09 |
| V180 | 1.77 | 0.14 | 20.46 | 1.68 | 0.67 | 0.10 | 1.16 | 0.30 | 25.52 | 2.19 | 1.22 | 0.30 |
| Q182 | 1.47 | 0.05 | 19.91 | 0.63 | 0.77 | 0.04 | 1.26 | 0.12 | 25.21 | 1.67 | 0.74 | 0.09 |
| E183 | 1.45 | 0.04 | 17.79 | 0.36 | 0.67 | 0.03 | 1.24 | 0.14 | 20.92 | 0.40 | 0.71 | 0.05 |
| G185 | 1.64 | 0.04 | 15.33 | 0.53 | 0.57 | 0.05 | 1.20 | 0.19 | 20.94 | 0.54 | 0.68 | 0.07 |
| R189 | 1.74 | 0.10 | - | - | 0.34 | 0.08 | 1.41 | 0.09 | 26.32 | 1.69 | 0.90 | 0.25 |
| E191 | 1.33 | 0.07 | 17.75 | 0.33 | 0.56 | 0.04 | 0.94 | 0.14 | 19.38 | 0.68 | 0.52 | 0.06 |
| T193 | 1.20 | 0.08 | 21.32 | 1.07 | 0.69 | 0.07 | 1.05 | 0.15 | 22.19 | 0.95 | 0.67 | 0.10 |
| M194 | 1.49 | 0.13 | 22.38 | 2.67 | 0.89 | 0.18 | 1.28 | 0.06 | 21.07 | 4.10 | 0.77 | 0.26 |
| T195 | 1.33 | 0.05 | 23.04 | 0.64 | 0.84 | 0.07 | 1.06 | 0.15 | 27.43 | 1.58 | 0.87 | 0.14 |
| C196 | 1.30 | 0.15 | 20.41 | 1.26 | 0.59 | 0.10 | 1.20 | 0.14 | 25.47 | 1.26 | 0.55 | 0.11 |
| R197 | 1.20 | 0.14 | 22.37 | 2.66 | 1.05 | 0.19 | 1.38 | 0.13 | 26.97 | 1.80 | 0.76 | 0.21 |
| V198 | 1.15 | 0.10 | 19.37 | 1.51 | 0.77 | 0.14 | 1.03 | 0.20 | 24.74 | 3.10 | 1.08 | 0.29 |
| E199 | 1.39 | 0.05 | 18.73 | 0.55 | 0.70 | 0.04 | 1.02 | 0.07 | 20.50 | 0.76 | 0.78 | 0.08 |
| R200 | 1.30 | 0.11 | 17.60 | 0.98 | 0.80 | 0.13 | 1.16 | 0.10 | 20.62 | 2.50 | 0.59 | 0.18 |
| F201 | 1.41 | 0.06 | 20.46 | 0.67 | 0.74 | 0.04 | 1.00 | 0.08 | 22.79 | 0.92 | 0.76 | 0.06 |
| I202 | 1.33 | 0.05 | 19.62 | 0.78 | 0.69 | 0.05 | 1.00 | 0.09 | 20.53 | 0.59 | 0.84 | 0.10 |
| E203 | 1.46 | 0.02 | 19.29 | 0.65 | 0.77 | 0.04 | 0.99 | 0.10 | 24.02 | 0.46 | 0.77 | 0.08 |
| I204 | 1.22 | 0.05 | 20.52 | 1.00 | 0.76 | 0.08 | 0.86 | 0.09 | 22.86 | 0.81 | 0.87 | 0.15 |
| G205 | 1.52 | 0.05 | 19.70 | 0.73 | 0.85 | 0.06 | 1.11 | 0.16 | 19.37 | 0.87 | 1.02 | 0.13 |

| | | | | | | | | | | | | |
|---|---|---|---|---|---|---|---|---|---|---|---|---|
| S206 | 1.36 | 0.04 | 20.38 | 0.54 | 0.74 | 0.03 | 0.95 | 0.13 | 20.90 | 0.65 | 0.80 | 0.06 |
| G207 | 1.35 | 0.04 | 22.78 | 0.80 | 0.77 | 0.04 | 1.00 | 0.12 | 22.56 | 0.79 | 0.97 | 0.10 |
| T208 | 1.61 | 0.10 | 19.56 | 0.63 | 0.69 | 0.05 | 1.31 | 0.24 | 22.45 | 1.75 | 0.94 | 0.12 |
| S209 | 1.32 | 0.07 | 10.29 | 2.68 | 0.11 | 0.04 | 1.06 | 0.15 | 7.85 | 5.43 | 0.49 | 0.17 |
| L212 | 1.07 | 0.14 | 22.60 | 2.24 | 0.97 | 0.11 | 0.76 | 0.22 | 23.43 | 1.98 | 0.86 | 0.17 |
| A213 | 1.43 | 0.05 | 17.74 | 1.45 | 0.82 | 0.09 | 0.98 | 0.11 | 20.26 | 1.86 | 0.95 | 0.17 |
| R215 | 1.37 | 0.12 | 20.01 | 2.07 | 1.03 | 0.15 | 0.92 | 0.05 | 25.59 | 1.33 | 0.77 | 0.15 |
| N216 | 1.26 | 0.05 | 24.39 | 0.72 | 0.75 | 0.04 | 0.87 | 0.10 | 24.44 | 0.92 | 0.77 | 0.07 |
| A217 | 1.39 | 0.03 | 18.58 | 1.56 | 0.64 | 0.08 | 1.06 | 0.10 | 23.62 | 1.23 | 1.08 | 0.21 |
| A219 | 1.37 | 0.05 | 24.46 | 0.93 | 0.83 | 0.06 | 1.01 | 0.01 | 27.16 | 0.21 | 0.72 | 0.13 |
| L223 | 1.18 | 0.03 | 24.26 | 0.56 | 0.74 | 0.07 | 0.92 | 0.06 | 27.62 | 0.81 | 0.85 | 0.10 |
| R224 | 1.47 | 0.13 | 19.59 | 1.09 | 0.84 | 0.14 | 0.95 | 0.16 | 25.31 | 2.80 | 0.74 | 0.18 |
| V225 | 1.43 | 0.09 | 18.70 | 1.00 | 0.75 | 0.14 | 0.95 | 0.14 | 23.69 | 2.26 | 0.81 | 0.22 |
| T227 | 1.70 | 0.10 | 15.33 | 0.31 | 0.53 | 0.03 | 1.36 | 0.18 | 16.08 | 0.24 | 0.64 | 0.05 |
| V228 | 1.52 | 0.05 | 11.29 | 0.20 | 0.48 | 0.04 | 1.31 | 0.12 | 13.59 | 0.38 | 0.63 | 0.06 |
| L230 | 0.97 | 0.01 | 4.70 | 0.09 | -0.93 | -0.01 | 1.02 | 0.04 | 2.33 | 0.60 | -0.63 | -0.01 |
| A232 | 1.35 | 0.04 | 4.85 | 0.19 | -0.25 | -0.02 | 1.29 | 0.14 | 5.94 | 0.37 | 0.29 | 0.03 |

**Table S10**: $R_{2eff}$ values measured at different CPMG frequencies from CPMG relaxation dispersion experiment for bound TRBP2-dsRBD2 at 600 MHz NMR spectrometer.

| Residue Number | $R_{2eff}$ at CPMG frequency ($s^{-1}$) | | | | | | | | | | | | | |
|---|---|---|---|---|---|---|---|---|---|---|---|---|---|---|
| | | | | | | | | | | | | Repeat | | |
| | 25 | 50 | 75 | 125 | 175 | 275 | 375 | 525 | 675 | 825 | 1000 | 125 | 375 | 825 |
| error | 2.52 | 2.21 | 1.84 | 2.15 | 1.98 | 3.09 | 2.93 | 2.52 | 2.52 | 2.93 | 2.97 | 2.73 | 2.90 | 2.99 |
| S151 | 6.51 | 3.25 | 6.04 | 3.11 | 1.98 | 3.44 | 7.08 | 8.93 | 5.47 | 6.04 | 9.84 | 2.54 | 6.00 | 8.69 |
| N152 | 7.15 | 6.40 | 6.58 | 6.09 | 4.19 | 8.78 | 8.75 | 9.35 | 11.15 | 6.55 | 11.67 | 6.47 | 7.74 | 11.67 |
| A153 | 5.19 | 5.37 | 6.15 | 6.21 | 6.76 | 6.68 | 6.51 | 7.28 | 8.83 | 6.15 | 9.70 | 6.11 | 6.98 | 9.14 |
| Q154 | 2.87 | 2.51 | 2.99 | 3.31 | 6.21 | 4.60 | 4.25 | 5.85 | 6.27 | 2.99 | 7.90 | 3.32 | 4.93 | 7.49 |
| Q155 | 0.96 | 1.65 | 1.59 | 1.30 | 4.47 | 2.14 | 2.30 | 3.45 | 4.79 | 1.59 | 6.20 | 1.56 | 2.44 | 5.42 |
| S156 | 22.20 | 21.30 | 20.80 | 22.50 | 22.00 | 18.60 | 23.50 | 20.20 | 26.50 | 20.80 | 24.40 | 21.60 | 21.80 | 24.80 |
| E157 | 13.70 | 12.60 | 12.20 | 11.20 | 9.84 | 12.10 | 11.40 | 12.10 | 12.70 | 12.40 | 13.80 | 11.20 | 10.90 | 13.30 |
| C158 | 5.30 | 5.51 | 5.94 | 5.51 | 1.87 | 7.03 | 7.48 | 6.13 | 9.75 | 5.93 | 10.90 | 5.72 | 7.944 | 9.75 |
| G162 | 18.00 | 20.60 | 19.90 | 20.00 | 17.60 | 17.40 | 19.80 | 23.70 | 23.20 | 19.90 | 22.30 | 23.10 | 24.00 | 22.40 |
| A163 | 32.50 | 25.00 | 21.20 | 24.20 | 19.50 | 22.00 | 26.10 | 23.80 | 28.20 | 21.20 | 24.40 | 22.00 | 21.60 | 26.40 |
| L164 | 21.87 | 22.89 | 19.81 | 21.48 | 19.63 | 23.48 | 25.89 | 25.21 | 20.26 | 19.81 | 21.48 | 22.07 | 20.54 | 23.90 |
| Q165 | 23.20 | 24.30 | 20.20 | 19.30 | 22.80 | 20.50 | 20.30 | 21.90 | 22.00 | 20.20 | 25.00 | 22.50 | 22.10 | 24.20 |
| E166 | 21.50 | 20.10 | 20.20 | 20.70 | 19.80 | 18.00 | 20.30 | 20.70 | 23.60 | 20.20 | 25.40 | 19.80 | 20.80 | 22.70 |
| L167 | 16.70 | 17.80 | 23.20 | 17.50 | 22.10 | 20.00 | 20.30 | 18.20 | 22.10 | 23.00 | 22.40 | 21.10 | 19.70 | 25.20 |
| V168 | 11.70 | 12.40 | 9.58 | 11.00 | 9.46 | 10.70 | 10.30 | 11.10 | 12.40 | 9.58 | 14.60 | 10.80 | 11.70 | 9.98 |
| V169 | 24.80 | 25.70 | 25.40 | 24.00 | 16.20 | 22.30 | 20.00 | 24.50 | 26.30 | 25.40 | 23.80 | 31.60 | 22.90 | 26.70 |
| Q170 | 23.00 | 21.80 | 22.40 | 21.30 | 22.70 | 23.30 | 21.40 | 21.20 | 21.70 | 22.40 | 25.40 | 20.90 | 22.40 | 23.40 |
| G172 | 19.00 | 17.50 | 16.70 | 18.60 | 18.00 | 19.20 | 15.80 | 18.80 | 20.20 | 16.70 | 20.00 | 15.90 | 16.90 | 20.80 |
| W173 | 17.40 | 17.10 | 16.90 | 18.30 | 19.10 | 18.50 | 19.20 | 19.80 | 17.90 | 17.20 | 19.20 | 16.10 | 17.40 | 18.90 |
| E177 | 16.30 | 17.00 | 19.40 | 15.10 | 16.10 | 18.80 | 21.70 | 16.90 | 18.60 | 19.40 | 19.20 | 19.20 | 17.70 | 21.70 |
| Y178 | 11.60 | 11.70 | 14.90 | 13.70 | 15.60 | 13.60 | 13.60 | 13.20 | 15.40 | 14.90 | 21.40 | 15.80 | 18.10 | 14.00 |
| T179 | 19.60 | 15.30 | 18.70 | 19.50 | 18.40 | 19.00 | 20.20 | 19.00 | 18.00 | 18.70 | 24.00 | 18.70 | 21.70 | 24.80 |
| V180 | 16.60 | 16.60 | 16.60 | 13.80 | 18.10 | 20.90 | 16.60 | 16.30 | 19.50 | 16.60 | 16.10 | 21.10 | 11.70 | 20.20 |
| T181 | 24.10 | 17.50 | 18.90 | 19.00 | 19.20 | 21.80 | 22.80 | 15.10 | 23.60 | 18.90 | 21.80 | 17.10 | 23.60 | 19.00 |
| Q182 | 19.10 | 22.80 | 18.50 | 19.10 | 18.70 | 15.00 | 17.70 | 18.10 | 17.70 | 18.50 | 23.20 | 18.50 | 20.70 | 17.10 |
| E183 | 19.50 | 17.80 | 20.80 | 19.50 | 19.10 | 18.70 | 21.40 | 18.30 | 20.10 | 20.80 | 19.90 | 20.70 | 20.60 | 21.10 |
| S184 | 16.87 | 12.49 | 16.08 | 10.31 | 10.31 | 6.15 | 12.11 | 14.72 | 11.43 | 16.07 | 15.11 | 15.76 | 15.51 | 17.08 |
| G185 | 16.40 | 20.40 | 16.40 | 14.10 | 18.80 | 16.00 | 17.20 | 12.10 | 15.40 | 16.40 | 21.80 | 20.10 | 17.90 | 17.70 |
| A187 | 23.70 | 23.21 | 19.29 | 23.37 | 23.37 | 35.06 | 24.14 | 20.03 | 30.58 | 19.81 | 31.74 | 22.32 | 37.16 | 20.48 |
| R189 | 14.07 | 21.04 | 16.42 | 13.58 | 13.58 | 15.71 | 17.90 | 13.99 | 20.05 | 16.07 | 16.74 | 21.90 | 17.66 | 14.72 |
| E191 | 18.40 | 17.40 | 16.80 | 16.40 | 18.00 | 16.50 | 14.60 | 18.60 | 15.40 | 16.80 | 19.00 | 18.50 | 14.90 | 17.50 |
| T193 | 23.80 | 18.00 | 23.70 | 24.10 | 22.60 | 21.00 | 16.20 | 21.80 | 19.00 | 23.70 | 22.80 | 21.10 | 20.20 | 25.10 |
| M194 | 17.80 | 13.00 | 13.00 | 26.50 | 8.49 | 27.50 | 13.50 | 14.00 | 21.30 | 13.00 | 24.40 | 26.80 | 9.79 | 12.50 |
| T195 | 30.10 | 23.50 | 25.10 | 21.10 | 26.60 | 25.80 | 19.20 | 18.90 | 30.90 | 25.10 | 28.00 | 23.50 | 28.60 | 24.00 |
| C196 | 17.10 | 13.90 | 19.40 | 29.20 | 13.20 | 19.50 | 23.40 | 8.60 | 19.40 | 20.30 | 21.90 | 19.40 | 8.65 | 15.50 |
| R197 | 15.60 | 13.10 | 11.50 | 13.70 | 17.80 | 19.00 | 15.20 | 13.30 | 25.10 | 21.60 | 27.40 | 26.10 | 11.80 | 27.40 |
| V198 | 18.10 | 17.90 | 21.70 | 22.50 | 18.80 | 12.40 | 13.90 | 21.50 | 11.40 | 12.60 | 14.20 | 23.00 | 29.70 | 19.80 |
| E199 | 18.70 | 19.80 | 17.40 | 18.20 | 20.70 | 20.30 | 19.60 | 20.30 | 18.50 | 17.40 | 20.90 | 19.10 | 20.00 | 20.80 |

| | | | | | | | | | | | | | | |
|---|---|---|---|---|---|---|---|---|---|---|---|---|---|---|
| R200 | 21.80 | 27.10 | 25.80 | 14.30 | 21.90 | 27.20 | 17.70 | 27.90 | 21.70 | 26.60 | 25.20 | 16.10 | 21.20 | 20.40 |
| F201 | 21.50 | 19.60 | 19.70 | 21.20 | 20.00 | 17.90 | 19.50 | 19.30 | 23.00 | 19.70 | 22.40 | 17.90 | 20.10 | 23.30 |
| I202 | 15.20 | 18.60 | 15.00 | 15.50 | 15.20 | 16.70 | 17.20 | 22.00 | 20.60 | 15.00 | 16.70 | 8.95 | 22.20 | 19.80 |
| E203 | 21.30 | 17.90 | 17.10 | 17.10 | 20.10 | 19.30 | 20.80 | 20.40 | 21.20 | 17.10 | 23.20 | 17.40 | 19.80 | 23.90 |
| I204 | 26.90 | 18.20 | 24.00 | 16.30 | 21.00 | 24.30 | 23.30 | 22.00 | 22.40 | 24.00 | 25.60 | 27.50 | 25.30 | 24.10 |
| G205 | 20.50 | 18.00 | 19.00 | 20.50 | 17.90 | 24.00 | 21.60 | 15.30 | 24.50 | 19.00 | 25.10 | 19.70 | 26.20 | 21.20 |
| S206 | 18.40 | 18.30 | 16.20 | 15.50 | 16.30 | 18.40 | 18.70 | 18.90 | 21.30 | 16.20 | 21.50 | 19.20 | 20.30 | 17.80 |
| G207 | 19.30 | 20.20 | 18.80 | 19.30 | 17.50 | 18.70 | 17.40 | 20.40 | 19.20 | 18.80 | 22.00 | 20.70 | 22.40 | 19.00 |
| T208 | 11.70 | 16.80 | 11.80 | 23.50 | 16.90 | 12.50 | 14.50 | 22.60 | 9.57 | 11.80 | 11.80 | 13.50 | 17.40 | 18.60 |
| L212 | 17.20 | 24.10 | 22.50 | 22.30 | 25.30 | 15.90 | 16.40 | 17.30 | 22.60 | 22.50 | 27.60 | 25.80 | 15.60 | 25.80 |
| A213 | 18.30 | 18.70 | 21.70 | 18.00 | 16.50 | 15.70 | 21.20 | 15.70 | 22.70 | 22.20 | 19.60 | 18.50 | 19.10 | 15.90 |
| R215 | 21.70 | 17.70 | 26.60 | 17.30 | 26.70 | 17.10 | 18.50 | 16.00 | 21.80 | 26.60 | 28.90 | 21.40 | 12.20 | 27.40 |
| N216 | 6.54 | 3.31 | 6.10 | 3.17 | 4.25 | 3.50 | 7.14 | 9.19 | 5.51 | 6.10 | 9.90 | 2.60 | 6.06 | 8.75 |
| A217 | 21.40 | 17.30 | 16.10 | 23.50 | 27.00 | 22.40 | 24.10 | 23.00 | 23.00 | 16.10 | 27.20 | 21.70 | 22.90 | 20.80 |
| A218 | 13.10 | 20.60 | 19.20 | 19.00 | 18.90 | 18.70 | 19.40 | 22.20 | 18.10 | 19.20 | 17.40 | 19.40 | 17.20 | 26.60 |
| A219 | 19.90 | 19.50 | 21.30 | 20.60 | 21.80 | 21.50 | 22.90 | 21.60 | 22.50 | 21.30 | 29.90 | 23.40 | 23.50 | 25.00 |
| M221 | 12.58 | 13.92 | 12.18 | 8.97 | 13.30 | 15.67 | 15.22 | 25.21 | 16.92 | 12.18 | 15.53 | 14.11 | 14.37 | 17.40 |
| L222 | 19.90 | 21.50 | 19.80 | 18.70 | 21.00 | 20.30 | 18.90 | 20.10 | 20.10 | 19.80 | 27.00 | 18.20 | 21.80 | 23.00 |
| R224 | 23.50 | 15.60 | 21.60 | 18.10 | 21.80 | 20.40 | 14.10 | 22.30 | 23.20 | 21.60 | 21.70 | 19.50 | 21.80 | 25.80 |
| L223 | 22.80 | 21.10 | 20.10 | 19.30 | 20.20 | 21.80 | 23.50 | 25.30 | 26.00 | 20.10 | 39.40 | 21.10 | 33.00 | 34.50 |
| V225 | 17.40 | 15.30 | 21.60 | 16.20 | 18.10 | 17.50 | 25.70 | 19.10 | 16.20 | 21.80 | 22.60 | 17.50 | 16.70 | 22.50 |
| T227 | 14.70 | 15.00 | 12.30 | 13.60 | 13.40 | 13.90 | 15.50 | 12.60 | 14.60 | 12.30 | 17.00 | 13.40 | 15.40 | 13.60 |
| V228 | 9.09 | 9.59 | 8.98 | 8.41 | 9.01 | 9.09 | 9.48 | 10.10 | 10.70 | 8.98 | 12.70 | 8.36 | 9.54 | 11.60 |
| L230 | 0.68 | 1.61 | 1.85 | 1.85 | 2.58 | 2.34 | 3.14 | 3.35 | 4.65 | 1.14 | 5.83 | 2.34 | 2.58 | 4.94 |
| A232 | 2.43 | 2.63 | 3.01 | 3.01 | 3.60 | 4.82 | 4.82 | 5.94 | 6.94 | 3.01 | 9.06 | 2.86 | 4.60 | 8.17 |

**Table S11**: $R_{1\rho}$ relaxation rates measured using HSn pulses (n=1,2,4,6,8) from HARD experiment for RNA-bound TRBP2-dsRBD2 at 600 MHz NMR spectrometer.

| Residue Number | $R_{1\rho}$ ($s^{-1}$) | | | | | | | | | |
|---|---|---|---|---|---|---|---|---|---|---|
| | HS1 | | HS2 | | HS4 | | HS6 | | HS8 | |
| | Value | Error | Value | Error | Value | Error | Value | Error | Value | Error |
| A153 | 2.45 | 0.11 | 4.29 | 0.30 | 4.62 | 0.18 | 4.73 | 0.28 | 3.58 | 0.11 |
| Q154 | 2.68 | 0.03 | 4.68 | 0.28 | 4.94 | 0.14 | 5.27 | 0.30 | 3.41 | 0.09 |
| Q155 | 2.35 | 0.12 | 4.54 | 0.39 | 4.48 | 0.27 | 5.06 | 0.20 | 2.93 | 0.16 |
| S156 | 2.73 | 0.28 | 5.41 | 0.32 | 5.56 | 0.38 | 6.20 | 0.17 | 4.38 | 0.23 |
| E157 | 3.52 | 0.10 | 5.42 | 0.23 | 6.00 | 0.15 | 6.14 | 0.25 | 4.62 | 0.13 |
| C158 | 2.99 | 0.74 | 6.28 | 0.74 | 5.45 | 0.56 | 5.27 | 0.56 | 4.47 | 0.25 |
| N159 | 3.72 | 0.91 | 7.78 | 1.05 | 7.49 | 1.12 | 6.83 | 1.32 | 7.02 | 0.73 |
| G162 | 5.52 | 0.64 | 7.17 | 0.55 | 10.55 | 1.03 | 10.27 | 0.49 | 8.35 | 0.67 |
| Q165 | 4.61 | 0.22 | 9.11 | 1.27 | 12.24 | 1.89 | 9.34 | 1.48 | 8.33 | 1.12 |
| E166 | 5.04 | 0.08 | 7.73 | 0.16 | 9.16 | 0.45 | 9.67 | 0.51 | 8.88 | 0.31 |
| L167 | 5.93 | 0.04 | 7.80 | 0.25 | 8.47 | 0.19 | 8.43 | 0.39 | 8.74 | 0.13 |
| V168 | 5.34 | 0.77 | 8.95 | 0.46 | 9.29 | 0.81 | 9.78 | 0.58 | 9.73 | 0.50 |
| V169 | 4.53 | 0.76 | 10.95 | 2.28 | 5.62 | 2.43 | 8.88 | 2.08 | 10.24 | 0.97 |
| Q170 | 5.32 | 0.17 | 9.17 | 1.07 | 9.96 | 0.50 | 9.62 | 1.06 | 8.55 | 1.00 |
| G172 | 4.52 | 0.17 | 7.66 | 0.21 | 9.12 | 0.29 | 10.36 | 0.53 | 9.55 | 0.29 |
| W173 | 4.85 | 0.29 | 7.79 | 0.19 | 8.23 | 0.38 | 9.37 | 0.31 | 8.66 | 0.61 |
| E177 | 3.59 | 0.28 | 6.66 | 0.29 | 8.56 | 0.29 | 8.78 | 0.30 | 7.75 | 0.39 |
| Y178 | 4.88 | 0.12 | 5.15 | 0.39 | 6.02 | 0.24 | 6.20 | 0.47 | 4.98 | 0.37 |
| T179 | 4.57 | 0.28 | 6.85 | 0.72 | 7.93 | 0.34 | 8.49 | 0.59 | 7.59 | 0.62 |
| V180 | 5.04 | 0.70 | 7.11 | 0.60 | 8.87 | 0.39 | 9.68 | 0.35 | 8.34 | 0.42 |
| T181 | 4.77 | 1.02 | 7.60 | 1.28 | 7.97 | 0.38 | 10.72 | 0.94 | 7.86 | 0.76 |
| Q182 | 4.78 | 0.19 | 6.02 | 0.25 | 7.32 | 0.26 | 8.07 | 0.43 | 8.15 | 1.22 |
| E183 | 4.23 | 0.17 | 6.58 | 0.13 | 7.93 | 0.52 | 9.48 | 0.41 | 8.03 | 0.41 |
| G185 | 4.20 | 0.42 | 7.33 | 0.59 | 7.93 | 0.31 | 7.82 | 0.41 | 7.52 | 0.07 |
| R189 | 3.01 | 0.30 | 6.19 | 0.73 | 7.10 | 0.62 | 8.28 | 0.69 | 7.09 | 0.29 |
| E191 | 4.53 | 0.05 | 5.86 | 0.32 | 7.91 | 0.54 | 8.25 | 0.23 | 6.99 | 0.30 |
| T193 | 4.78 | 0.19 | 8.12 | 0.68 | 8.69 | 0.68 | 9.54 | 0.61 | 9.16 | 0.54 |
| M194 | 5.75 | 0.46 | 4.95 | 2.17 | 4.54 | 0.79 | 9.66 | 2.35 | 6.67 | 1.91 |
| T195 | 5.53 | 0.28 | 7.36 | 0.45 | 8.92 | 0.23 | 9.22 | 0.50 | 10.38 | 0.52 |
| C196 | 4.12 | 0.96 | 6.61 | 0.82 | 7.87 | 0.95 | 7.39 | 0.77 | 9.97 | 0.94 |
| R197 | 3.37 | 1.01 | 8.05 | 1.54 | 10.52 | 1.42 | 8.28 | 0.30 | 8.51 | 0.67 |
| V198 | 5.68 | 1.05 | 5.97 | 1.41 | 9.38 | 1.48 | 11.55 | 1.59 | 7.68 | 1.36 |
| E199 | 4.52 | 0.18 | 7.68 | 0.20 | 8.30 | 0.29 | 8.48 | 0.36 | 8.16 | 0.12 |
| 200 | 4.27 | 0.52 | 9.13 | 1.56 | 8.82 | 0.70 | 7.76 | 0.48 | 8.73 | 1.00 |
| F201 | 5.03 | 0.12 | 8.21 | 0.73 | 9.68 | 0.11 | 9.98 | 0.17 | 8.59 | 0.24 |
| I202 | 5.35 | 0.34 | 6.76 | 0.79 | 9.12 | 0.87 | 8.26 | 0.70 | 7.22 | 0.51 |
| E203 | 4.32 | 0.11 | 7.00 | 0.33 | 7.90 | 0.37 | 8.19 | 0.27 | 8.46 | 0.13 |
| I204 | 5.17 | 0.15 | 8.06 | 0.71 | 8.92 | 0.52 | 8.63 | 0.69 | 8.78 | 0.33 |
| G205 | 4.24 | 0.37 | 7.66 | 0.67 | 8.34 | 0.57 | 9.79 | 0.38 | 8.71 | 0.78 |

| | | | | | | | | | | |
|---|---|---|---|---|---|---|---|---|---|---|
| S206 | 4.59 | 0.10 | 6.61 | 0.35 | 8.13 | 0.22 | 8.90 | 0.14 | 7.85 | 0.16 |
| G207 | 4.53 | 0.12 | 7.51 | 0.40 | 8.30 | 0.15 | 9.52 | 0.25 | 8.73 | 0.29 |
| T208 | 4.35 | 0.51 | 6.20 | 0.98 | 7.47 | 0.85 | 9.98 | 0.72 | 8.05 | 0.59 |
| S209 | 3.69 | 0.89 | 6.11 | 0.96 | 6.73 | 0.25 | 7.06 | 0.17 | 4.79 | 1.26 |
| L212 | 4.91 | 0.50 | 7.60 | 0.51 | 10.10 | 1.63 | 10.33 | 1.17 | 9.31 | 1.06 |
| A213 | 4.17 | 0.43 | 7.24 | 0.67 | 8.28 | 0.65 | 9.23 | 0.71 | 7.89 | 0.70 |
| R215 | 4.97 | 0.97 | 7.98 | 1.13 | 10.78 | 1.19 | 9.56 | 1.69 | 9.36 | 1.12 |
| N216 | 4.56 | 0.19 | 7.88 | 0.63 | 9.82 | 0.16 | 11.03 | 0.42 | 9.44 | 0.56 |
| A218 | 5.93 | 0.22 | 6.79 | 0.93 | 9.53 | 0.27 | 11.19 | 0.49 | 10.61 | 1.04 |
| A219 | 5.63 | 0.63 | 8.24 | 0.59 | 10.28 | 0.22 | 10.26 | 0.54 | 9.15 | 0.41 |
| M221 | 4.02 | 0.37 | 8.04 | 0.77 | 8.79 | 0.78 | 9.02 | 1.18 | 6.86 | 0.56 |
| L223 | 4.53 | 0.32 | 7.39 | 0.36 | 7.90 | 0.31 | 9.71 | 0.80 | 9.38 | 0.22 |
| R224 | 4.50 | 0.36 | 6.61 | 0.73 | 9.11 | 0.44 | 9.80 | 0.29 | 8.96 | 0.87 |
| V225 | 5.51 | 0.54 | 10.74 | 0.96 | 7.06 | 1.11 | 8.82 | 0.89 | 10.48 | 0.44 |
| T227 | 3.86 | 0.29 | 5.99 | 0.65 | 7.00 | 0.26 | 7.24 | 0.31 | 6.81 | 0.40 |
| V228 | 3.02 | 0.30 | 6.23 | 0.37 | 6.32 | 0.18 | 6.30 | 0.19 | 5.06 | 0.22 |
| L230 | 1.66 | 0.03 | 3.69 | 0.36 | 3.90 | 0.11 | 3.84 | 0.31 | 1.85 | 0.05 |
| A232 | 2.09 | 0.09 | 4.69 | 0.37 | 4.64 | 0.12 | 4.49 | 0.28 | 2.68 | 0.13 |

**Table S12**: $R_{2\rho}$ relaxation rates measured using HSn pulses (n=1,2,4,6,8) from HARD experiment for RNA-bound TRBP2-dsRBD2 at 600 MHz NMR spectrometer.

| Residue Number | $R_{2\rho}$ ($s^{-1}$) | | | | | | | | | |
|---|---|---|---|---|---|---|---|---|---|---|
| | HS1 | | HS2 | | HS4 | | HS6 | | HS8 | |
| | Value | Error | Value | Error | Value | Error | Value | Error | Value | Error |
| S151 | 7.52 | 0.88 | 7.36 | 0.08 | 8.09 | 0.39 | 9.85 | 0.93 | 6.44 | 0.17 |
| A153 | 7.20 | 0.93 | 7.03 | 0.12 | 6.90 | 0.66 | 9.42 | 0.20 | 6.89 | 0.38 |
| Q154 | 5.95 | 0.19 | 5.43 | 0.29 | 5.39 | 0.12 | 7.74 | 0.00 | 5.41 | 0.54 |
| Q155 | 4.06 | 0.56 | 4.54 | 0.50 | 4.69 | 2.13 | 7.14 | 2.55 | 4.43 | 0.96 |
| S156 | 11.98 | 1.44 | 10.14 | 0.77 | 9.29 | 0.40 | 11.78 | 0.66 | 8.98 | 0.10 |
| E157 | 11.42 | 0.63 | 11.44 | 0.90 | 10.55 | 0.93 | 13.03 | 0.04 | 11.13 | 0.32 |
| C158 | 10.99 | 0.56 | 7.60 | 2.88 | 8.80 | 3.36 | 5.86 | 1.50 | 5.66 | 0.04 |
| N159 | 22.76 | 1.58 | 24.66 | 4.86 | 26.77 | 5.51 | 26.46 | 6.61 | 25.79 | 0.72 |
| G162 | 22.34 | 0.84 | 22.53 | 2.55 | 22.89 | 2.43 | 19.21 | 6.19 | 18.27 | 4.48 |
| A163 | 25.68 | 5.44 | 20.88 | 0.59 | 20.24 | 2.68 | 29.83 | 10.75 | 23.13 | 5.24 |
| L164 | 22.11 | 1.08 | 21.66 | 1.43 | 21.37 | 2.58 | 22.51 | 5.18 | 19.30 | 2.13 |
| Q165 | 22.98 | 0.15 | 22.62 | 1.66 | 22.82 | 1.03 | 22.58 | 1.74 | 19.23 | 0.98 |
| E166 | 21.15 | 0.97 | 23.39 | 1.79 | 21.98 | 0.91 | 21.58 | 5.12 | 19.53 | 0.44 |
| L167 | 23.56 | 1.19 | 22.80 | 0.47 | 22.95 | 1.58 | 25.44 | 6.71 | 17.64 | 1.41 |
| V168 | 24.93 | 11.77 | 31.39 | 3.93 | 29.52 | 3.91 | 28.40 | 18.94 | 18.46 | 0.43 |
| V169 | 20.39 | 2.98 | 23.93 | 0.90 | 21.70 | 0.94 | 17.28 | 2.39 | 17.25 | 2.25 |
| Q170 | 21.15 | 0.74 | 22.25 | 0.06 | 21.78 | 0.45 | 22.11 | 2.39 | 19.74 | 1.63 |
| G172 | 21.47 | 2.52 | 19.56 | 2.17 | 19.37 | 1.05 | 19.66 | 0.29 | 20.35 | 0.38 |
| W173 | 20.46 | 1.15 | 19.23 | 1.52 | 19.90 | 2.37 | 22.82 | 0.06 | 19.05 | 0.04 |
| R174 | 15.98 | 2.78 | 15.03 | 0.86 | 18.07 | 3.56 | 17.64 | 5.95 | 12.51 | 1.18 |
| E177 | 12.92 | 0.77 | 12.70 | 0.80 | 13.02 | 0.65 | 15.59 | 0.79 | 12.96 | 0.30 |
| Y178 | 20.19 | 0.27 | 16.56 | 0.74 | 15.63 | 0.37 | 18.41 | 3.01 | 17.56 | 0.94 |
| T179 | 18.75 | 1.11 | 19.94 | 0.53 | 18.82 | 1.83 | 22.96 | 4.09 | 18.83 | 0.36 |
| V180 | 22.01 | 3.27 | 21.46 | 2.48 | 20.08 | 4.19 | 23.91 | 0.50 | 26.12 | 2.81 |
| T181 | 21.68 | 1.07 | 18.85 | 0.86 | 26.22 | 2.28 | 22.72 | 0.69 | 18.85 | 1.54 |
| Q182 | 22.61 | 1.38 | 20.11 | 0.82 | 20.71 | 0.04 | 22.64 | 1.13 | 23.06 | 3.42 |
| E183 | 18.22 | 3.48 | 17.77 | 1.30 | 18.62 | 2.35 | 22.15 | 0.40 | 17.84 | 1.83 |
| G185 | 17.93 | 0.46 | 17.18 | 0.71 | 18.36 | 2.42 | 18.95 | 2.71 | 17.48 | 0.21 |
| E191 | 17.60 | 1.33 | 18.25 | 0.17 | 17.37 | 1.06 | 27.71 | 1.83 | 17.71 | 0.94 |
| T193 | 20.75 | 1.00 | 24.53 | 1.32 | 23.89 | 1.95 | 19.42 | 1.43 | 23.31 | 2.38 |
| M194 | 21.64 | 5.16 | 19.92 | 12.84 | 15.64 | 2.44 | 21.92 | 0.14 | 17.78 | 8.54 |
| T195 | 22.00 | 0.70 | 21.89 | 2.18 | 20.70 | 0.25 | 20.08 | 1.54 | 19.74 | 2.44 |
| C196 | 19.54 | 1.76 | 17.44 | 0.32 | 18.67 | 0.14 | 15.72 | 6.59 | 20.17 | 2.55 |
| R197 | 23.88 | 3.14 | 18.10 | 5.39 | 16.82 | 0.17 | 14.73 | 3.25 | 18.00 | 4.74 |
| V198 | 21.00 | 3.24 | 15.94 | 5.64 | 15.97 | 2.03 | 12.01 | 1.39 | 15.78 | 2.91 |
| E199 | 17.92 | 0.09 | 19.06 | 1.85 | 18.11 | 1.08 | 17.92 | 0.24 | 17.62 | 0.03 |
| 200 | 17.91 | 6.24 | 20.25 | 1.94 | 24.53 | 1.66 | 16.34 | 7.13 | 15.38 | 0.16 |
| F201 | 19.22 | 0.12 | 17.94 | 0.24 | 18.44 | 1.67 | 18.23 | 1.48 | 16.84 | 0.65 |
| I202 | 20.00 | 1.61 | 19.45 | 0.39 | 16.81 | 1.66 | 20.95 | 0.46 | 17.96 | 0.02 |

| | | | | | | | | | | |
|---|---|---|---|---|---|---|---|---|---|---|
| E203 | 19.80 | 1.40 | 20.52 | 0.20 | 18.91 | 1.30 | 17.10 | 3.39 | 17.46 | 1.94 |
| I204 | 20.08 | 1.11 | 21.68 | 0.27 | 21.36 | 1.32 | 22.82 | 3.02 | 20.50 | 3.25 |
| G205 | 22.26 | 0.40 | 18.03 | 0.24 | 19.48 | 0.28 | 18.40 | 2.92 | 18.65 | 3.01 |
| S206 | 18.69 | 0.50 | 17.92 | 2.08 | 17.61 | 0.08 | 18.80 | 0.02 | 17.81 | 0.10 |
| G207 | 22.45 | 1.73 | 20.50 | 2.59 | 21.27 | 1.82 | 21.01 | 2.15 | 20.08 | 1.31 |
| T208 | 19.50 | 0.64 | 23.86 | 0.48 | 20.49 | 0.79 | 17.94 | 0.23 | 16.75 | 3.78 |
| S209 | 14.26 | 0.66 | 12.31 | 0.20 | 14.55 | 1.79 | 9.77 | 9.13 | 14.36 | 3.10 |
| L212 | 23.28 | 1.15 | 27.70 | 2.68 | 24.48 | 0.90 | 23.33 | 5.02 | 21.71 | 7.43 |
| A213 | 22.66 | 4.90 | 20.19 | 1.04 | 20.21 | 0.53 | 15.27 | 12.49 | 19.65 | 1.12 |
| R215 | 20.61 | 1.26 | 26.71 | 1.26 | 23.27 | 4.53 | 18.58 | 11.00 | 17.22 | 3.91 |
| N216 | 25.37 | 2.54 | 23.62 | 0.11 | 21.27 | 2.74 | 24.28 | 2.86 | 24.34 | 0.35 |
| A218 | 23.38 | 0.51 | 23.58 | 0.66 | 20.59 | 6.55 | 23.59 | 7.26 | 20.66 | 0.21 |
| A219 | 24.02 | 1.34 | 22.84 | 0.63 | 21.45 | 1.20 | 24.16 | 3.88 | 21.11 | 1.95 |
| M221 | 15.46 | 1.37 | 13.39 | 1.01 | 17.72 | 3.58 | 16.92 | 0.88 | 9.83 | 1.58 |
| L223 | 18.21 | 0.54 | 23.29 | 1.03 | 21.64 | 0.76 | 24.19 | 8.45 | 20.25 | 0.39 |
| R224 | 20.78 | 1.38 | 27.11 | 0.01 | 26.05 | 2.72 | 21.81 | 10.88 | 19.37 | 3.91 |
| V225 | 17.02 | 2.64 | 22.54 | 9.64 | 12.75 | 3.81 | 12.07 | 15.39 | 19.89 | 1.51 |
| T227 | 15.66 | 1.96 | 13.51 | 0.75 | 13.80 | 0.66 | 14.58 | 1.57 | 14.46 | 0.17 |
| V228 | 11.44 | 2.07 | 10.40 | 1.40 | 11.43 | 1.75 | 13.93 | 0.57 | 10.89 | 0.65 |
| L230 | 2.51 | 0.23 | 2.28 | 0.13 | 2.56 | 0.51 | 5.01 | 0.36 | 2.66 | 0.00 |
| A232 | 4.35 | 0.21 | 3.89 | 0.21 | 4.25 | 0.55 | 6.80 | 0.24 | 4.39 | 0.47 |

**Table S13**: $R_1$ relaxation rates measured from HARD experiment for RNA-bound TRBP2-dsRBD2 at 600 MHz NMR spectrometer.

| Residue Number | $R_1$ ($s^{-1}$) | |
|---|---|---|
| | Value | Error |
| S151 | 1.36 | 0.10 |
| A153 | 1.48 | 0.02 |
| Q154 | 1.47 | 0.03 |
| Q155 | 1.17 | 0.04 |
| S156 | 1.41 | 0.13 |
| E157 | 1.55 | 0.11 |
| C158 | 1.51 | 0.17 |
| N159 | 1.67 | 0.24 |
| G162 | 1.21 | 0.21 |
| A163 | 1.28 | 0.11 |
| Q165 | 1.30 | 0.15 |
| E166 | 1.35 | 0.10 |
| L167 | 1.15 | 0.18 |
| V168 | 1.36 | 0.30 |
| V169 | 1.40 | 0.12 |
| Q170 | 1.29 | 0.05 |
| G172 | 1.35 | 0.05 |
| W173 | 1.33 | 0.11 |
| E177 | 1.26 | 0.08 |
| Y178 | 1.43 | 0.08 |
| T179 | 1.39 | 0.09 |
| V180 | 1.50 | 0.23 |
| T181 | 1.46 | 0.13 |
| Q182 | 1.56 | 0.08 |
| E183 | 1.41 | 0.10 |
| G185 | 1.46 | 0.14 |
| R189 | 1.51 | 0.22 |
| E191 | 1.41 | 0.07 |
| T193 | 1.44 | 0.18 |
| M194 | 1.37 | 0.42 |
| T195 | 1.34 | 0.18 |
| C196 | 1.61 | 0.09 |
| R197 | 1.36 | 0.19 |
| V198 | 1.54 | 0.38 |
| E199 | 1.46 | 0.08 |
| R200 | 0.96 | 0.23 |
| F201 | 1.45 | 0.04 |
| I202 | 1.35 | 0.10 |
| E203 | 1.52 | 0.06 |
| I204 | 1.22 | 0.11 |

| | | |
|---|---|---|
| G205 | 1.52 | 0.07 |
| S206 | 1.38 | 0.03 |
| G207 | 1.35 | 0.04 |
| T208 | 1.51 | 0.15 |
| S209 | 1.25 | 0.10 |
| L212 | 1.35 | 0.20 |
| A213 | 1.14 | 0.19 |
| R215 | 1.07 | 0.12 |
| N216 | 1.37 | 0.07 |
| A218 | 1.36 | 0.16 |
| A219 | 1.25 | 0.10 |
| M221 | 1.27 | 0.15 |
| L223 | 1.31 | 0.14 |
| R224 | 1.40 | 0.09 |
| V225 | 1.27 | 0.33 |
| T227 | 1.56 | 0.10 |
| V228 | 1.57 | 0.06 |
| L230 | 0.96 | 0.02 |

**Table S14**: Order parameter ($S^2$) extracted from Model-free analysis of the Nuclear Spin Relaxation data for apo TRBP2-dsRBD2 recorded at 600 MHz and 800 MHz NMR spectrometer.

| Residue Number | $S^2$ | |
|---|---|---|
| | Value | Error |
| G162 | 0.78 | 0.01 |
| A163 | 0.40 | 0.01 |
| L164 | 0.85 | 0.01 |
| Q165 | 0.80 | 0.00 |
| E166 | 0.81 | 0.01 |
| L167 | 0.73 | 0.01 |
| V169 | 0.62 | 0.02 |
| Q170 | 0.75 | 0.00 |
| G172 | 0.63 | 0.03 |
| W173 | 0.63 | 0.02 |
| R174 | 0.23 | 0.01 |
| L175 | 0.38 | 0.04 |
| E177 | 0.00 | 0.11 |
| Y178 | 0.60 | 0.04 |
| T179 | 0.75 | 0.01 |
| V180 | 0.12 | 0.28 |
| T181 | 0.73 | 0.03 |
| Q182 | 0.56 | 0.02 |
| E183 | 0.31 | 0.01 |
| G185 | 0.18 | 0.01 |
| R189 | 0.52 | 0.03 |
| E191 | 0.44 | 0.01 |
| T193 | 0.88 | 0.01 |
| M194 | 0.68 | 0.03 |
| T195 | 0.73 | 0.01 |
| C196 | 0.82 | 0.01 |
| R197 | 0.50 | 0.05 |
| V198 | 0.34 | 0.04 |
| R200 | 0.45 | 0.02 |
| F201 | 0.77 | 0.01 |
| I202 | 0.64 | 0.02 |
| E203 | 0.72 | 0.02 |
| I204 | 0.67 | 0.02 |
| G205 | 0.71 | 0.01 |
| S206 | 0.75 | 0.01 |
| G207 | 0.59 | 0.05 |
| T208 | 0.77 | 0.02 |
| K210 | 0.52 | 0.05 |

| | | |
|---|---|---|
| L212 | 0.71 | 0.12 |
| A213 | 0.83 | 0.01 |
| R215 | 0.69 | 0.01 |
| N216 | 0.74 | 0.01 |
| A218 | 0.74 | 0.01 |
| A219 | 0.91 | 0.01 |
| M221 | 0.47 | 0.02 |
| L223 | 0.65 | 0.09 |
| R224 | 0.58 | 0.02 |
| V225 | 0.67 | 0.02 |
| T227 | 0.24 | 0.01 |

**Table S15**: Order parameter ($S^2$) extracted from Model-free analysis of the Nuclear Spin Relaxation data for RNA-bound TRBP2-dsRBD2 recorded at 600 MHz and 800 MHz NMR spectrometer.

| Residue Number | $S^2$ | |
|---|---|---|
| | Value | Error |
| N159 | 0.22 | 0.01 |
| G162 | 0.98 | 0.02 |
| A163 | 0.27 | 0.08 |
| Q165 | 0.79 | 0.01 |
| V168 | 0.92 | 0.04 |
| V169 | 0.38 | 0.04 |
| Q170 | 0.96 | 0.02 |
| G172 | 0.93 | 0.02 |
| W173 | 0.82 | 0.03 |
| Y178 | 0.94 | 0.02 |
| T179 | 0.86 | 0.01 |
| Q182 | 0.57 | 0.02 |
| E183 | 0.31 | 0.01 |
| E191 | 0.29 | 0.06 |
| T193 | 0.84 | 0.03 |
| M194 | 0.88 | 0.04 |
| T195 | 0.81 | 0.03 |
| V198 | 0.61 | 0.05 |
| R200 | 0.49 | 0.03 |
| F201 | 0.89 | 0.02 |
| I202 | 0.72 | 0.02 |
| E203 | 0.94 | 0.01 |
| I204 | 0.83 | 0.03 |
| G205 | 0.65 | 0.03 |
| S206 | 0.89 | 0.02 |
| G207 | 0.81 | 0.03 |
| T208 | 0.92 | 0.03 |
| L212 | 0.93 | 0.04 |
| A213 | 0.94 | 0.03 |
| N216 | 0.85 | 0.03 |
| A219 | 0.91 | 0.01 |
| L223 | 0.90 | 0.02 |
| R224 | 0.78 | 0.03 |
| V225 | 0.64 | 0.04 |

**Table S16**: $R_{ex}$ of residues extracted from Model-free analysis of the Nuclear Spin Relaxation data for apo TRBP2-dsRBD2 recorded at 600 MHz and 800 MHz NMR spectrometer.

| Residue Number | $R_{ex}$ ($s^{-1}$) | |
|---|---|---|
| | Value | Error |
| E166 | 0.42 | 0.12 |
| E177 | 2.62 | 0.78 |
| V180 | 4.85 | 2.06 |
| T195 | 2.4 | 0.22 |
| L212 | 1.22 | 1.17 |
| N216 | 1.14 | 0.19 |
| L223 | 0.87 | 0.73 |

**Table S17**: $R_{ex}$ of residues extracted from Model-free analysis of Nuclear Spin Relaxation data for RNA-bound TRBP2-dsRBD2 recorded at 600 MHz and 800 MHz NMR spectrometer.

| Residue Number | $R_{ex}$ ($s^{-1}$) | |
|---|---|---|
| | Value | Error |
| V168 | 4.70 | 1.35 |
| Y178 | 2.10 | 0.83 |
| T195 | 2.01 | 0.77 |
| I204 | 1.89 | 0.69 |
| N216 | 1.77 | 0.68 |
| L223 | 3.62 | 0.50 |

**Table S18:** Dynamics parameters extracted from HARD experimental data from geoHARD method for apo TRBP2-dsRBD2.

| Residue Number | $k_{ex}$ (Hz) | | $p_B$ (%) |
|---|---|---|---|
| | Value | Error | |
| N159 | 193.02 | 74.20 | 2.18 |
| G162 | 108.91 | 9.01 | 1.39 |
| A163 | 143.20 | 40.06 | 1.10 |
| Q165 | 145.51 | 32.02 | 1.60 |
| E166 | 113.27 | 11.37 | 0.94 |
| L167 | 153.11 | 78.02 | 0.63 |
| V169 | 181.02 | 63.29 | 1.41 |
| G172 | 149.65 | 32.10 | 1.64 |
| W173 | 111.43 | 13.12 | 0.99 |
| E177 | 144.32 | 24.23 | 1.43 |
| Y178 | 149.17 | 45.08 | 1.34 |
| T179 | 116.90 | 6.58 | 1.05 |
| V180 | 30162.81 | 2556.08 | 4.98 |
| T181 | 225.86 | 113.77 | 1.16 |
| Q182 | 104.47 | 6.91 | 1.12 |
| E183 | 151.01 | 54.24 | 0.92 |
| R189 | 31748.72 | 955.20 | 9.37 |
| E191 | 185.94 | 58.22 | 0.89 |
| T193 | 113.50 | 14.73 | 1.51 |
| T195 | 110.02 | 13.94 | 1.60 |
| C196 | 164.04 | 75.77 | 1.18 |
| V198 | 30554.87 | 1687.74 | 3.05 |
| E199 | 115.53 | 14.31 | 1.31 |
| R200 | 53667.71 | 2331.35 | 15.82 |
| F201 | 148.85 | 63.15 | 0.56 |
| I202 | 153.01 | 64.86 | 0.74 |
| E203 | 129.64 | 34.33 | 0.84 |
| 205 | 153.79 | 40.99 | 0.96 |
| S206 | 137.90 | 32.54 | 1.26 |
| G207 | 204.01 | 64.95 | 1.34 |
| T208 | 173.36 | 69.90 | 0.81 |
| S209 | 110.46 | 12.18 | 2.16 |
| L212 | 36081.01 | 4503.77 | 10.56 |
| A213 | 121.17 | 17.10 | 1.53 |
| N216 | 195.81 | 71.97 | 1.07 |
| A217 | 160.24 | 74.26 | 1.40 |
| A218 | 162.39 | 53.99 | 1.45 |
| A219 | 138.44 | 43.89 | 1.74 |
| L223 | 107.89 | 13.48 | 0.95 |
| R224 | 31519.02 | 1155.44 | 15.57 |

| | | | |
|---|---|---|---|
| T227 | 52411.81 | 1883.58 | 7.57 |

**Table S19:** Dynamics parameters extracted from HARD experimental data from geoHARD method for RNA-bound TRBP2-dsRBD2.

| Residue Number | $k_{ex}$ (Hz) | | $p_B$ (%) |
|---|---|---|---|
| | Value | Error | |
| N159 | 109.84 | 6.09 | 8.18 |
| G162 | 33694.84 | 4498.63 | 14.58 |
| Q165 | 144.71 | 39.88 | 3.07 |
| E166 | 117.25 | 18.81 | 3.31 |
| L167 | 33136.76 | 1028.77 | 13.75 |
| V168 | 31361.87 | 381.57 | 23.84 |
| V169 | 157.41 | 44.83 | 3.44 |
| Q170 | 116.96 | 11.92 | 2.92 |
| G172 | 175.47 | 86.44 | 2.47 |
| W173 | 114.07 | 8.10 | 3.51 |
| T179 | 109.72 | 7.08 | 4.01 |
| V180 | 143.37 | 29.57 | 5.40 |
| T181 | 114.44 | 6.67 | 4.89 |
| Q182 | 137.39 | 19.91 | 4.48 |
| E183 | 119.61 | 13.26 | 4.15 |
| G185 | 130.95 | 25.10 | 2.92 |
| E191 | 110.37 | 8.93 | 4.66 |
| T193 | 111.15 | 9.42 | 3.73 |
| C196 | 117.34 | 15.76 | 3.48 |
| R197 | 38548.93 | 3434.10 | 6.46 |
| V198 | 73599.49 | 5723.40 | 27.41 |
| E199 | 114.28 | 11.25 | 2.60 |
| R200 | 34143.21 | 678.61 | 19.36 |
| F201 | 171.94 | 65.01 | 1.11 |
| E203 | 34467.84 | 1310.56 | 21.48 |
| I204 | 111.56 | 15.16 | 3.32 |
| G205 | 154.09 | 34.56 | 2.51 |
| S206 | 120.58 | 16.23 | 3.03 |
| G207 | 29981.15 | 4358.72 | 4.63 |
| T208 | 148.23 | 48.27 | 3.66 |
| S209 | 125.06 | 16.57 | 1.77 |
| L212 | 120.49 | 18.57 | 4.42 |
| A213 | 27995.83 | 4031.29 | 14.63 |
| R215 | 136.63 | 33.31 | 2.96 |
| N216 | 111.11 | 5.78 | 5.34 |
| A219 | 34874.65 | 4100.95 | 15.65 |
| M221 | 61676.83 | 1371.30 | 24.59 |
| L223 | 101.90 | 2.14 | 3.50 |
| R224 | 109.67 | 9.36 | 5.47 |
| V225 | 125.89 | 18.09 | 1.15 |
